# The dual function receptor kinase, OsWAKL21.2, is involved in elaboration of lipaseA/esterase induced immune responses in rice

**DOI:** 10.1101/754234

**Authors:** Kamal Kumar Malukani, Ashish Ranjan, Hota Shiva Jyothi, Hitendra Kumar Patel, Ramesh V. Sonti

## Abstract

Plant pathogens secrete cell wall degrading enzymes (CWDEs) to degrade various components of the plant cell wall. Plants sense this cell wall damage as a mark of infection and induce immune responses. Little is known about the plant functions that are involved in the elaboration of cell wall damage-induced immune responses. Transcriptome analysis revealed that a rice receptor kinase, *WALL-ASSOCIATED KINASE-LIKE 21* (*OsWAKL21.2*), is upregulated following treatment with either *Xanthomonas oryzae* pv. *oryzae* (*Xoo*, a bacterial pathogen) or lipaseA/esterase (LipA: a CWDE of *Xoo*). Downregulation of *OsWAKL21.2* attenuates LipA mediated immune responses. Overexpression of *OsWAKL21.2* in rice mimics LipA treatment mediated induction of immune responses and enhanced expression of defence related genes, indicating it could be involved in the perception of LipA induced cell wall damage in rice. OsWAKL21.2 is a dual function kinase having *in-vitro* kinase and guanylate cyclase (GC) activities. Ectopic expression of *OsWAKL21.2* in Arabidopsis also activates plant immune responses. Interestingly, OsWAKL21.2 needs kinase activity to activate rice immune responses while in Arabidopsis it needs GC activity. Our study reveals a novel receptor kinase involved in elaboration of cell wall damage induced rice immune responses that can activate similar immune responses in two different species via two different mechanisms.

**One sentence Summary:** A novel rice receptor WAKL21 that sense cell wall damage caused by Xanthomonas secreted cell wall degrading enzyme to induce immune responses.

## Introduction

The plant cell wall acts as a formidable barrier for pathogens. Plant pathogens secrete a battery of cell wall degrading enzymes (CWDEs) to degrade different components of the plant cell wall (Albersheim and Anderson-Prouty, 1975, Hématy et al., 2009). CWDEs act as a double-edged sword for pathogens as on one hand the activity of these enzymes leads to cell wall degradation, on the other hand, it releases cell wall degradation products that can elicit plant immune responses (Jha et al., 2007). Such host derived molecules that can elicit immune responses are called damage associated molecular patterns (DAMPs). Some known cell wall degradation products that act as DAMPs include pectin degradation products oligogalacturonide (OG), hemicellulose degradation products such as xyloglucan oligomers, and cellulose degradation products such as cellobiose and cellotriose (Gust et al., 2017, de Azevedo Souza et al., 2017, Claverie et al., 2018). These DAMPs are sensed by membrane-localised receptor-like kinases (RLKs) that activate the signaling cascade. Some known receptors of the DAMPs are AtPEPR1/2 for plant elicitor peptides (Pep), AtDORN1 for eATP, SYR1 for systemins and AtWAK1/2 for oligogalacturonide (OG) (Brutus et al., 2010, Gust et al., 2017, Wang et al., 2018).

The wall-associated kinases (WAKs) constitute a unique class of receptor kinases which are known to be closely associated with the plant cell wall (Verica and He, 2002). WAKs are known to be involved in many physiological processes including cell elongation, pollen development and abiotic and biotic stress tolerance (Kohorn, 2015). Members of the WAK gene family have been known to interact with pectin and pectin degradation products (OGs). AtWAK1 and AtWAK2 have been reported to interact with pectin and OGs *in vitro* (Kohorn et al., 2006, Kohorn et al., 2009). Some proteins of the WAK gene family have also been known to be involved in immune responses in many plant species such as Arabidopsis, rice, maize and wheat (He et al., 1998, Li et al., 2009, Zhang et al., 2017, Zuo et al., 2015, Hurni et al., 2015, Harkenrider et al., 2016, Hu et al., 2017, Saintenac et al., 2018). In most of the cases, a receptor kinase or receptor-coreceptor complex recognises the ligand and triggers phosphorylation events leading to activation of MAP kinase signaling and its downstream targets (Meng and Zhang, 2013). However, some recent studies also indicate the presence of an alternate signaling system in plants which is mediated by cyclic nucleotides such as cyclic guanosine monophosphate (cGMP) and cyclic adenosine monophosphate (cAMP) (Gehring and Turek, 2017). cGMP is generated by guanylate cyclases (GCs) and most of the reported plant GCs are membrane localised receptor kinases that also contain a functional GC motif inside the kinase domain (Gehring and Turek, 2017). Such kinases showing these dual activities are called moonlighting kinases (Wong et al., 2015). In Arabidopsis, some receptor kinases including a wall associated kinase like gene (*AtWAKL10*) are reported as moonlighting kinases (Meier et al., 2010).

Rice serves as a staple food for more than half of the world population. *Xanthomonas oryzae* pv. *oryzae* (*Xoo*) causes the serious bacterial blight disease of rice. CWDEs secreted by *Xoo* include cellulases, xylanases and lipases/esterases (LipA) (Rajeshwari et al., 2005, Jha et al., 2007). LipA is an important CWDE of *Xoo* and deletion of the LipA gene results in a significant reduction in the virulence of *Xoo* in rice (Jha et al., 2007). Treatment of rice tissue with purified LipA leads to the activation of plant immune responses including callose deposition, programmed cell death and an enhanced tolerance towards *Xoo* (Aparna et al., 2009). The mechanism of action of LipA on the cell wall is still not clear, but it has been predicted that it acts by cleaving ester linkages in the rice cell wall (Aparna et al., 2009). Heat inactivation or mutation of the active site residues of LipA abolishes the biochemical activity as well as the ability to induce immune responses in rice, indicating that the enzymatic activity of LipA is essential for the induction of immune response (Jha et al., 2007, Aparna et al., 2009). However, the process through which rice senses the cell wall damage caused by LipA and further activates immune responses is not clear.

In this study, transcriptome analysis was initially performed to identify gene expression changes that occur during LipA induced immune responses in rice. An enhanced transcript level of a wall-associated kinase like gene, *OsWAKL21.2* was observed after treatment of rice leaves with either purified LipA or the pathogen, *Xoo*, but not after treatment with a LipA mutant of *Xoo*. Sequence alignment and biochemical studies indicate that OsWAKL21.2 is a dual function receptor kinase that has an *in vitro* kinase as well as a GC activity. OsWAKL21.2 is a key component of signaling involved in LipA induced immunity as its downregulation leads to attenuation of LipA induced immune response. Overexpression of *OsWAKL21.2* in rice and ectopic expression in Arabidopsis induces plant defence response and confers enhanced tolerance to subsequent bacterial infection. However, we have observed that the mode of action of the receptor is dissimilar in rice and Arabidopsis. Our results suggest that OsWAKL21.2 requires its kinase activity to induce immune response in rice, whereas, in Arabidopsis, it requires GC activity.

## Results

### Expression of *OsWAKL21.2* was enhanced after treatment of rice leaves either with LipA or *Xoo*

In order to identify rice functions that are potentially involved in early stages of LipA induced immune responses, we performed transcriptome analysis of rice leaves after 30 minutes and 2hr of infiltration with LipA. After 30 minutes, no gene was significantly altered while 78 genes (74 unique set of genes) were differentially expressed (68up, 10 down) (FC>1.5 fold) after 2hr of LipA treatment (Supplemental Fig. S1A, Supplemental Table S1). This includes genes that might have roles in signaling, defence responses or in transcription/translation (Supplemental Fig. S1B). When compared with a previous microarray (Ranjan et al., 2015) performed after 12hr of LipA treatment, we observed 38 of these 78 genes are differentially expressed (37up, 1 down) at both time points (Fig. 1A, Supplemental Table S2). We compared with a publicly available microarray dataset that was performed 24hr after treatment of rice leaves with various *Xanthomonas oryzae* strains (GEO Acc. No. GSE36272) and observed that some of these 38 genes were commonly upregulated following *Xanthomonas* treatment (Supplemental Table S3). The upregulation of six of these commonly upregulated genes was validated by qRT-PCR after treatment of rice leaves with either *Xoo* or LipA (Supplemental Fig. S1C). Three of the 37 genes that were most commonly upregulated after *Xanthomonas* treatments include a putative wall-associated receptor kinase like gene (*OsWAKL21*, LOC_Os12g40419), a putative ubiquitin ligase (*OsPUB38*, LOC_Os04g35680) and a putative fructose-bisphosphate aldolase (LOC_Os08g02700) (Supplemental Table S3). Since the focus of this work was on the perception of cell wall damage in rice plants, we decided to explore the function of wall-associated receptor kinase OsWAKL21.

**Figure 1:**
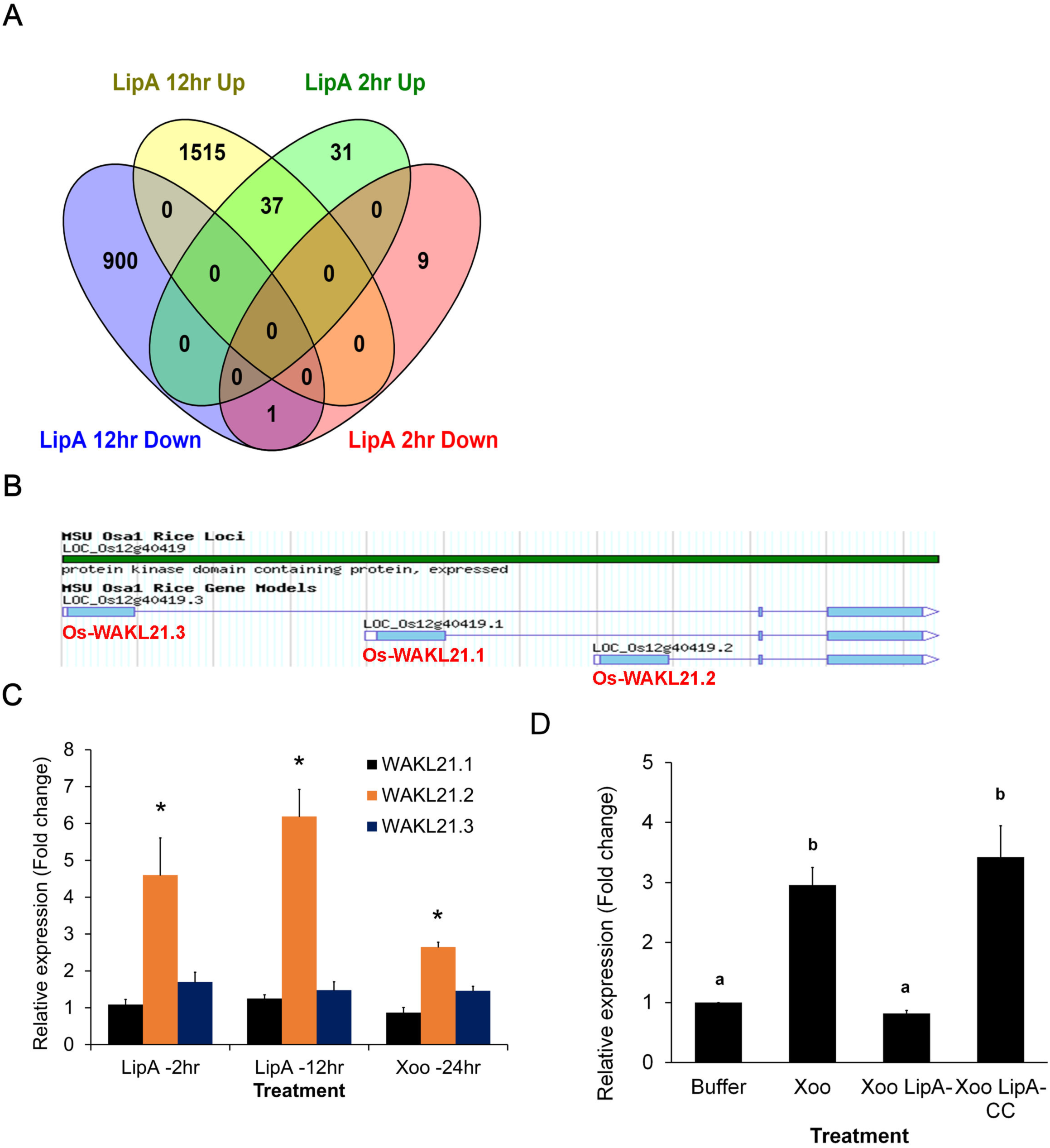
Expression of *OsWAKL21.2* is enhanced in rice leaves after treatment with either LipA or *Xoo*. A. Venn diagram indicating number of genes that are differentially expressed after 2hr and 12hr of LipA treatment.
B. Three splice variants of *OsWAKL21* as shown in Rice-MSU database.
C. qRT-PCR analysis of the expression of all three splice variants of *OsWAKL21* after 2hr and 12hr of LipA treatment, and after 24hr of *Xoo* treatment in rice leaves. Asterisk (*) represents significant difference in fold change with p<0.05 with respect to buffer treated leaves.
D. qRT-PCR analysis of expression of *OsWAKL21.2* in rice leaves after 24hr of treatment with either *Xoo*, LipA mutant of *Xoo* (*Xoo* LipA-) or LipA complementing clone of *Xoo* (*Xoo* LipA-CC). a and b above the bars indicate significant difference with p<0.05. In C and D, 12-14 days old leaves were infiltrated with either LipA (0.5mg/ml) or *Xoo* (O.D. 1.0). Each bar represents average value and error bar denotes standard error (SE) of at least three independent experiments. Relative expression was calculated in leaves treated with LipA or *Xoo* with respect to leaves treated with buffer. *OsActin1* was used as internal control for qRT-PCR. The relative fold change was calculated by using 2^−ΔΔ^ method.

*OsWAKL21* has three splice variants [*OsWAKL21.1* (LOC_Os12g40419.1)*, OsWAKL21.2* (LOC_Os12g40419.2) and *OsWAKL21.3* (LOC_Os12g40419.3)] (Fig. 1B). qRT-PCR analyses indicate that the second splice variant (*OsWAKL21.2*) is mainly upregulated in rice leaves after either LipA or *Xoo* treatment (Fig. 1C). Interestingly, treatment of rice leaves with LipA mutant of *Xoo* did not enhance expression of *OsWAKL21.2* while introduction of a LipA complementing clone into the LipA mutant restores the ability to enhance expression of *OsWAKL21.2* (Fig. 1D).

### Overexpression of *OsWAKL21.2* in rice mimics LipA induced immune responses

Treatment of rice tissue with LipA induces immune responses such as callose deposition, enhanced expression of defence related genes, activation of JA pathway and enhanced tolerance against subsequent *Xoo* infection (Jha et al., 2007, Ranjan et al., 2015). Agrobacterium mediated transient overexpression of *OsWAKL21.2* in young rice leaves significantly induces callose deposition which is comparable to callose deposition induced by LipA treatment (Fig. 2A,B). Transient overexpression of *OsWAKL21.2* in rice leaves also enhances tolerance against subsequent *Xoo* infection leading to reduced lesion length caused by *Xoo* which is also observed following treatment with LipA (Fig. 2C, Supplemental Fig. S2A). The overexpression of *OsWAKL21.2* was confirmed by qRT-PCR and Western blot analysis (Supplemental Fig. S2B,C).

**Figure 2:**
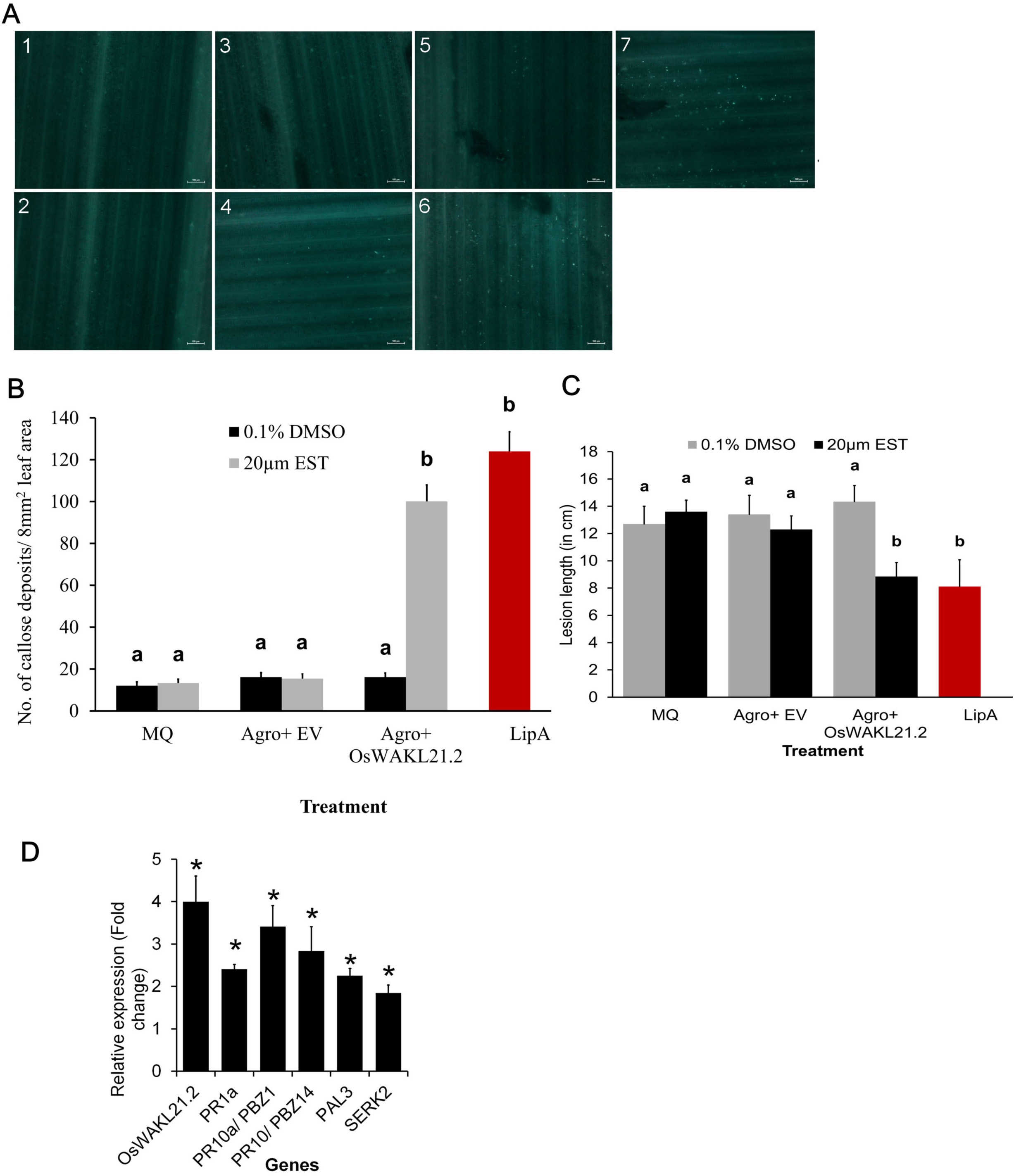
Overexpression of *OsWAKL21.2* in rice leaves induces plant immune responses. A. Callose deposition in rice leaves after treatment with various Agrobacterium constructs or controls. The image shown is representative image of one viewing area for each category. Scale bar represents 100µm. The numbers denotes: 1-0.1% DMSO, 2-20µM β-estradiol (Est), 3,4-Agrobacterium containing pMDC7 (Empty vector-EV) without (3) or with (4) inducer (Est), 5,6-Agrobacterium containing pMDC7::*OsWAKL21.2* without (5) or with (6) inducer (Est), 7-LipA.
B. Quantification of callose deposition in rice leaves after treatment with various Agrobacterium constructs or controls. Bar diagram showing the quantification of number of callose deposits per area in rice leaves. Number of callose deposits in 8 such viewing areas (as shown in Fig. 2A) per leaf were considered. Each bar represents the average and error bar represents SE of 10-15 leaves per treatment in one set of experiment. Similar results were obtained in three independent experiments.
C. Lesion length caused by *Xoo* in rice leaves when mid-vein of the leaves were previously treated with various Agrobacterium constructs or controls. Mid-veins of rice leaves of 60 day old plants were injected with either MQ, LipA or Agrobacterium carrying empty vector or *OsWAKL21.2* and also with (20µM β-estradiol) or without (0.1% DMSO) inducer. After 24hr, the leaves were pin prick inoculated with *Xoo*, 1cm above the point of Agrobacterium injection. Lesion length was measured after 10 days of infection of Xoo (Supplemental Fig. S2A). Each bar indicates average and error bar represents SE of >20 leaf per treatment in one set of experiment. Similar results were obtained in three independent experiments.
D. Relative expression of key defence related genes after transient overexpression of *OsWAKL21.2* in rice leaves. Each bar represents average fold change and the error bars indicate SE in three independent experiments (n=12 in each experiment). For each gene, transcript level of uninduced condition (treatment with Agrobacterium carrying *OsWAKL21.2* with 0.1% DMSO) was considered as 1 and was compared to induced condition (treatment with Agrobacterium carrying *OsWAKL21.2* with 20µM estradiol). *OsActin1* was used as internal control for qRT-PCR. The relative fold change was calculated by using 2^−ΔΔ^ method. Asterisk (*) represents significant difference in fold change with p<0.05 with respect to uninduced condition. In A and B, 12-14 days old rice leaves were infiltrated with either MQ, Agrobacterium carrying empty vector or vector containing *OsWAKL21.2* and also with (20µM β-estradiol) or without (0.1% DMSO) inducer. In B and C, a and b above the bars indicate significant difference with p<0.05. MQ (MilliQ or water) treatment indicate control without any Agrobacterium treatment. In A, B and C, Leaves treated with LipA were used as positive control.

Plant immune responses are known to be modulated via the expression of defence-related genes. Therefore, we tested the expression of some key defence-related genes of rice after the transient overexpression of *OsWAKL21.2* in mid-veinal regions of rice leaves. *OsWAKL21.2* overexpression in rice enhances expression of three pathogenesis-related genes (*OsPR1a*, *OsPR10*/*OsPBZ14* and *OsPR10a/OsPBZ1)*, a somatic embryogenesis receptor kinase (*OsSERK2*) and a phenylalanine ammonia lyase (*OsPAL3*) (Fig. 2D). We also tested expression of 10 genes that are upregulated following LipA/*Xoo* treatment (Supplemental Table S3) in microarray and observed seven of these ten genes are also significantly upregulated following overexpression of *OsWAKL21.2* in rice (Supplemental Fig. S2D). These results indicate that Agrobacterium-mediated transient overexpression of *OsWAKL21.2* in rice leaves mimics LipA treatment.

### Transient downregulation of *OsWAKL21.2* attenuates LipA induced immune responses in rice

We subsequently assessed the effect of transient knockdown of *OsWAKL21.2* by Virus-induced gene silencing (VIGS) on LipA induced immune responses. It was observed that the downregulation was not retained by all leaves for a long time which was also observed previously using this vector system (Kant and Dasgupta, 2017). So, an alternative approach was used for assessment of callose deposition in RNAi lines after LipA treatment (Supplemental Fig. S3). We categorized the leaf samples into three classes based on the amount of callose deposition as low (<30 deposits/leaf), medium (∼30-80 deposits/leaf) or high (>80 deposits/leaf) (Fig. 3A). Following LipA treatment, about 30-40% of the leaf samples showed high callose deposition, 10-15% showed low callose deposition while the rest of them (about 50%) showed a medium level of callose deposition (Fig. 3B). A similar ratio was observed if the seedlings were previously treated with VIGS-EV (Fig. 3C). The number of leaves showing low callose deposition significantly increased to more than 50% in WAKL-RNAi lines (WRI 1-300, WRI 450-600 and WRI 1-600 correspond to the fragment of *OsWAKL21.2* that was used for downregulation) while there was a reduction in the leaves that showed high or medium callose deposition (Fig. 3C). In RNAi lines, the leaves that show low callose deposition following LipA treatment also showed significantly lower transcript/protein level of OsWAKL21.2 which was not observed in the leaves that showed high callose deposition (Fig. 3D, Supplemental Fig. S4A).

**Figure 3:**
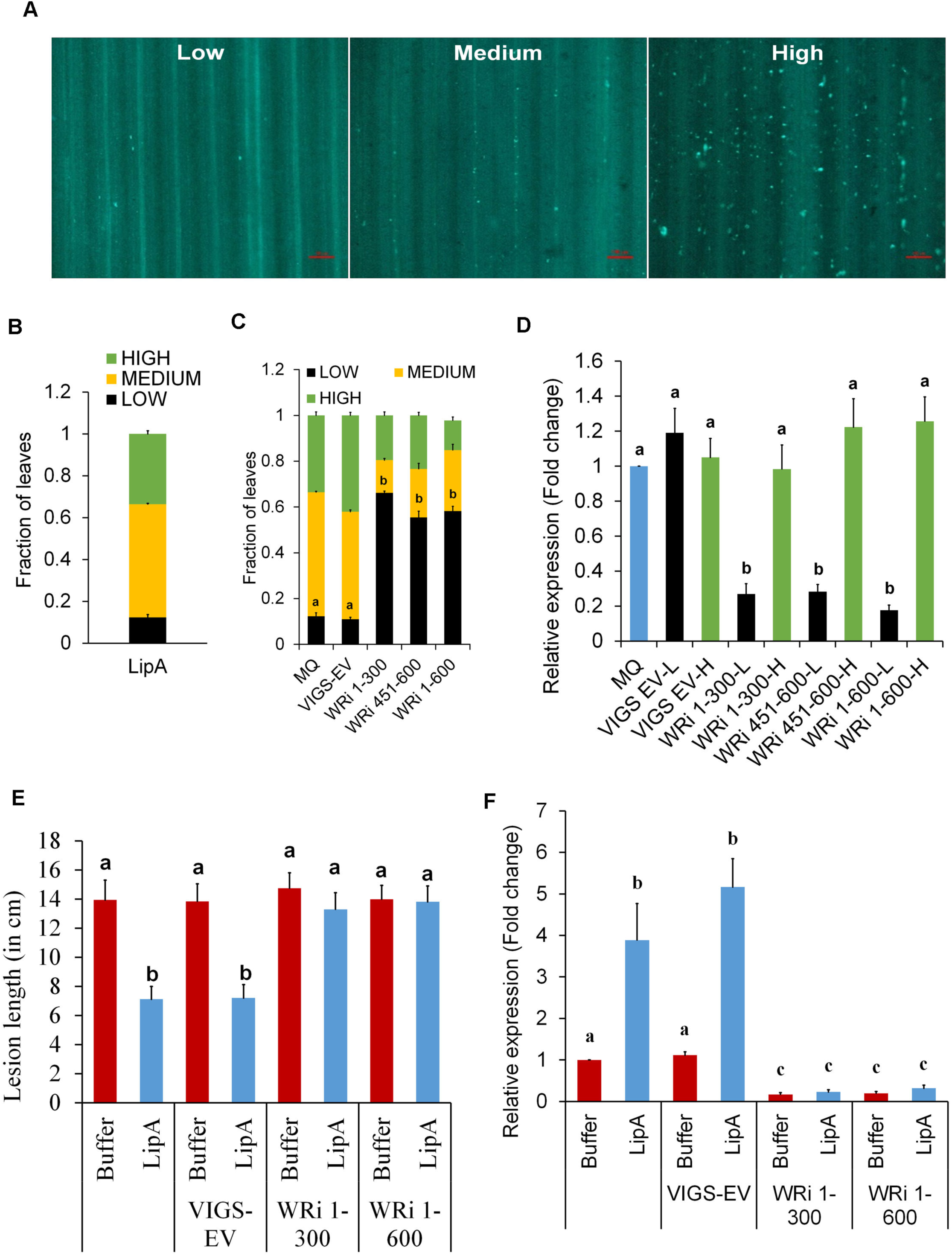
Downregulation of *OsWAKL21.2* attenuates LipA induced immune responses in rice. A. Categorization of number of callose deposits in three different groups: low, medium and high. The image shown is representative image of one viewing area for each group. 8 such areas per leaf were viewed for categorization.
B. Fraction of leaves showing low, medium or high callose deposition after LipA treatment.
C. Fraction of leaves showing callose deposits post LipA infiltration that were previously treated with either MQ (mock treatment), Agrobacterium containing VIGS-EV or WAK-RNAi constructs [WAKL-RNAi 1-300 (WRi 1-300), WAKL-RNAi 451-600 (WRi 451-600 or WAKL-RNAi 1-600 (WRi 1-600)] in 12-14 days old rice leaves.
D. qRT-PCR analysis of *OsWAKL21.2* transcript levels in leaves showing either low or high callose deposits (H: High callose, L: Low callose). Each bar represents average fold change and error bar indicates SE observed in three biological replicates. For each sample, 4-5 leaves showing respective callose phenotype were used for RNA isolation. Transcript level in mock (MQ) treated leaves was considered as 1 and fold change in Agrobacterium treated leaves was calculated with respect to it.
E. Lesion length caused by *Xoo* in mid-veins of 60 days old rice leaves that were pre-treated with either buffer and LipA alone or along with Agrobacterium strains [WAKL-RNAi 1-300 (WRi 1-300) or WALK-RNAi 1-600 (WRi 1-600)]. Each bar represents average lesion length and error bar show SE of at least 20 leaves in one experiment. Similar results were obtained in three independent experiments.
F. Expression level of *OsWAKL21.2* in rice leaves after 24hr of injection with either buffer and LipA alone or along with Agrobacterium strains [WAKL-RNAi 1-300 (WRi 1-300) or WAKL-RNAi 1-600 (WRi 1-600)]. Each bar represents average of three independent experiments, n>10 in each experiment. Transcript level of buffer injected leaves was considered as 1 and fold change in Agrobacterium with Buffer/LipA treated leaves was calculated with respect to it. In B, C, D and F, each bar represents the average and error bar denotes the SE of three different biological replicates. In B and C each bar denotes the ratio of leaves showing respective phenotype in at least 40 leaves. In C, D, E and F small letters (a, b and c) above the bars indicates significant difference with p<0.05. In D and F, *OsActin1* was used as internal control for qRT-PCR and the relative fold change was calculated by using 2^−ΔΔCt^ method.

Since downregulation of OsWAKL21.2 attenuated LipA induced callose deposition, we decided to test its effect on LipA induced tolerance towards *Xoo*. VIGS mediated downregulation of *OsWAKL21.2* in rice mid-vein attenuates LipA induced tolerance against subsequent *Xoo* infection (Fig. 3E, Supplemental Fig. S4B). qRT-PCR and Western blotting studies using anti-OsWAKL21 antibodies indicated the downregulation of OsWAKL21.2 in the mid vein following VIGS mediated *OsWAKL21.2* downregulation (Fig. 3F, Supplemental Fig. S4C). There was slight but usually non-significant reduction on transcript level of other splice variants and no significant difference was observed in transcript level of other predicted off-target genes (Supplemental Fig. S5). This suggests that optimal expression of *OsWAKL21.2* in rice leaves is required for LipA induced tolerance against *Xoo*.

### Ectopic expression of *OsWAKL21.2* in transgenic Arabidopsis lines induces plant immune responses

In order to determine whether expression of *OsWAKL21.2* would activate immune responses in other plants, we generated stable Arabidopsis transgenic lines expressing *OsWAKL21.2* under a 17-β-estradiol (Est) inducible promoter. Expression of *OsWAKL21.2* in transgenic β lines was examined after treatment with the inducer (Est) through qRT-PCR and Western blotting (Supplemental Fig. S6A,B). We observed that ectopic expression of *OsWAKL21.2* in Arabidopsis results in an enhanced callose deposition (Fig. 4A,B), enhanced tolerance against subsequent *Pseudomonas syringae* pv. *tomato* DC3000 (*Pst* DC3000) infection and reduction in *in planta* growth of *Pst* DC3000 (Fig. 4C). In Arabidopsis, the Salicylic acid (SA) and Jasmonic acid (JA) pathways are widely known to be involved in immune responses. We examined the expression of key genes linked to these two pathways in Arabidopsis transgenic lines. The ectopic expression of *OsWAKL21.2* in Arabidopsis resulted in a significant increase in the transcript levels of key SA pathway-related genes (*AtPR2, AtPR5*, and *AtWRKY33*) and *AtGSL5*, a major callose synthase of Arabidopsis (Fig. 4D) (Jacobs et al., 2003, Janda and Ruelland, 2015) while the transcript level of the key JA responsive gene *AtPDF1.2* was decreased. Overall, this data implies that ectopic expression of *OsWAKL21.2* in Arabidopsis, enhances callose deposition, enhances expression of SA pathway related genes, and in addition, enhances tolerance against subsequent *Pst* DC3000 infection.

**Figure 4:**
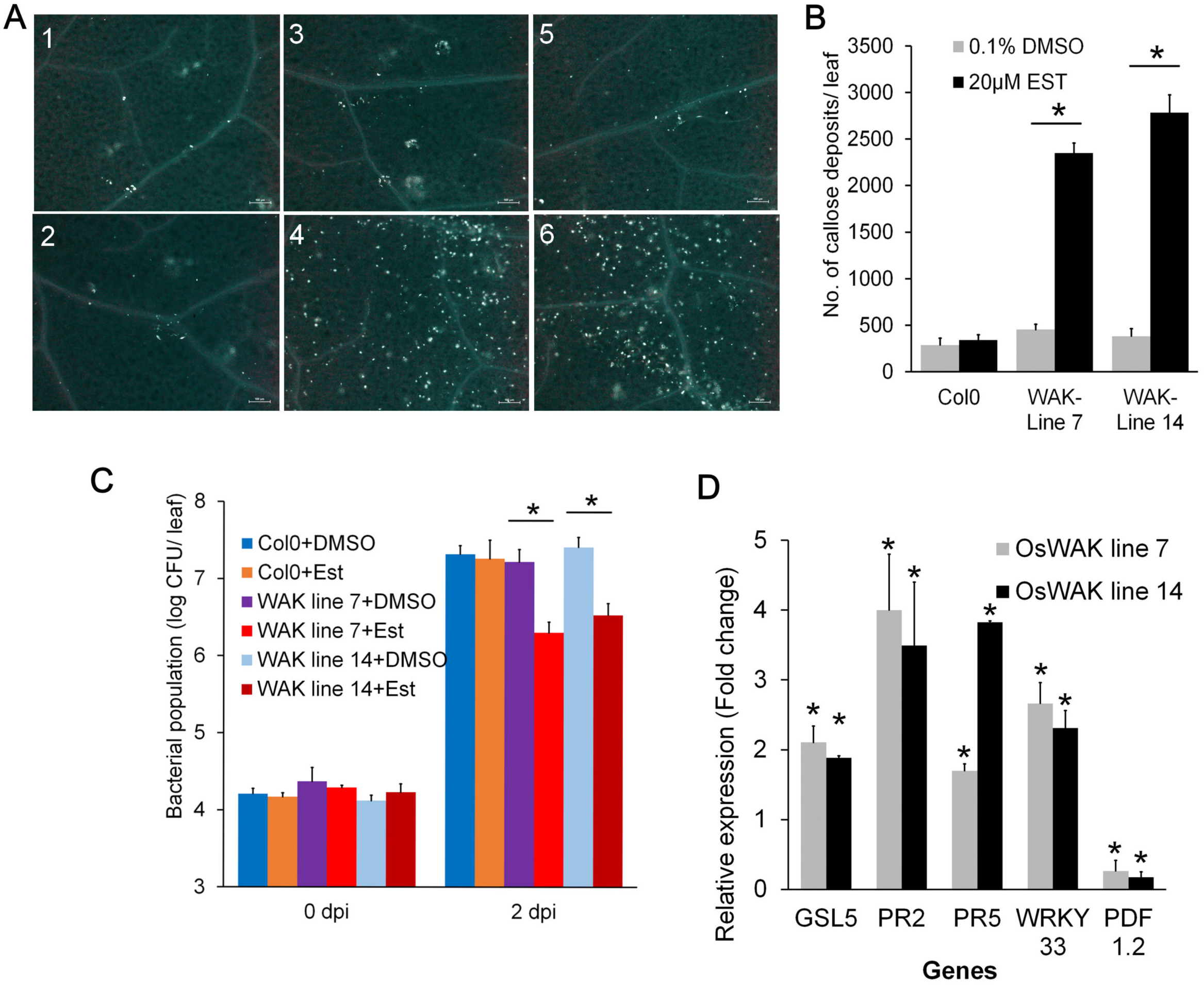
Ectopic expression of *OsWAKL21.2* in Arabidopsis induces immune responses. A. Callose deposition in leaves of wild type Columbia (Col-0) or *OsWAKL21.2* transgenic Arabidopsis lines following treatment with 20µM β-estradiol (inducer) or 0.1% DMSO (control). Numbers denote: 1,2-Col-0 treated with DMSO (1) or Est (2), 3,4-*OsWAKL21.2* transgenic line 7 treated with DMSO (3) or Est (4), 5,6-*OsWAKL21*.2 transgenic line 14 treated with DMSO (5) or Est (6).
B. Quantification of number of callose deposits in wild type Col-0 and two different Arabidopsis *OsWAKL21.2* transgenic lines after treatment with control or inducer. Leaves were treated with either 20µM β-estradiol (inducer) or 0.1% DMSO (control). Each bar represents the average and error bar represents SE of three different leaves for each treatment in an experiment.
C. Effect of ectopic expression of *OsWAKL21.2* on growth of *Pst* DC3000 after subsequent infection. Each bar represents average and error bar represents SE of five leaves for each treatment in an experiment.
D. Effect of ectopic expression of *OsWAKL21.2* in transgenic Arabidopsis lines on the expression of SA or JA pathway responsive genes. Expression in 0.1% DMSO treated leaves was considered as 1 and relative expression in 20µM estradiol treated leaves was calculated with respect to it. Each bar represents the average of three independent experiments for each line. For each sample, RNA was isolated from 3 leaves for every treatment. *AtActin2* was used as internal control for qRT-PCR. The relative fold change was calculated by using 2^−ΔΔCt^method Transgenic or wild type plant leaves were treated with 0.1% DMSO (Control) or 20 μM estradiol (inducer). 12hr later leaves were either collected for callose deposition or transcript/protein analysis or were infected with *Pst* DC3000. Similar results were obtained in three independent experiments for A, B and C. If the significant difference was observed, asterisk (*) represents significant difference with p<0.05 with respect to uninduced condition.

### OsWAKL21.2 is a membrane localizing moonlighting receptor kinase having *in vitro* kinase and guanylate cyclase (GC) activities

Sequence analyses of OsWAKL21.2 indicated that it is a receptor-like serine/threonine kinase that accommodates an N-terminal extracellular galacturonan binding domain (GBD), an epidermal growth factor (EGF) like repeat and an intracellular C-terminal kinase domain, resembling to other known wall-associated kinases (Fig. 5A). The analyses of OsWAKL21.2 also revealed the presence of a putative GC motif (from residue 569-585) inside the kinase domain (Supplemental Fig. S7A,B) (Xu et al., 2018). Enhanced GFP (EGFP) tagged recombinant OsWAKL21.2:EGFP localized to the cell membrane in onion epidermal cell indicating that it is a membrane bound receptor (Fig. 5B).

**Figure 5:**
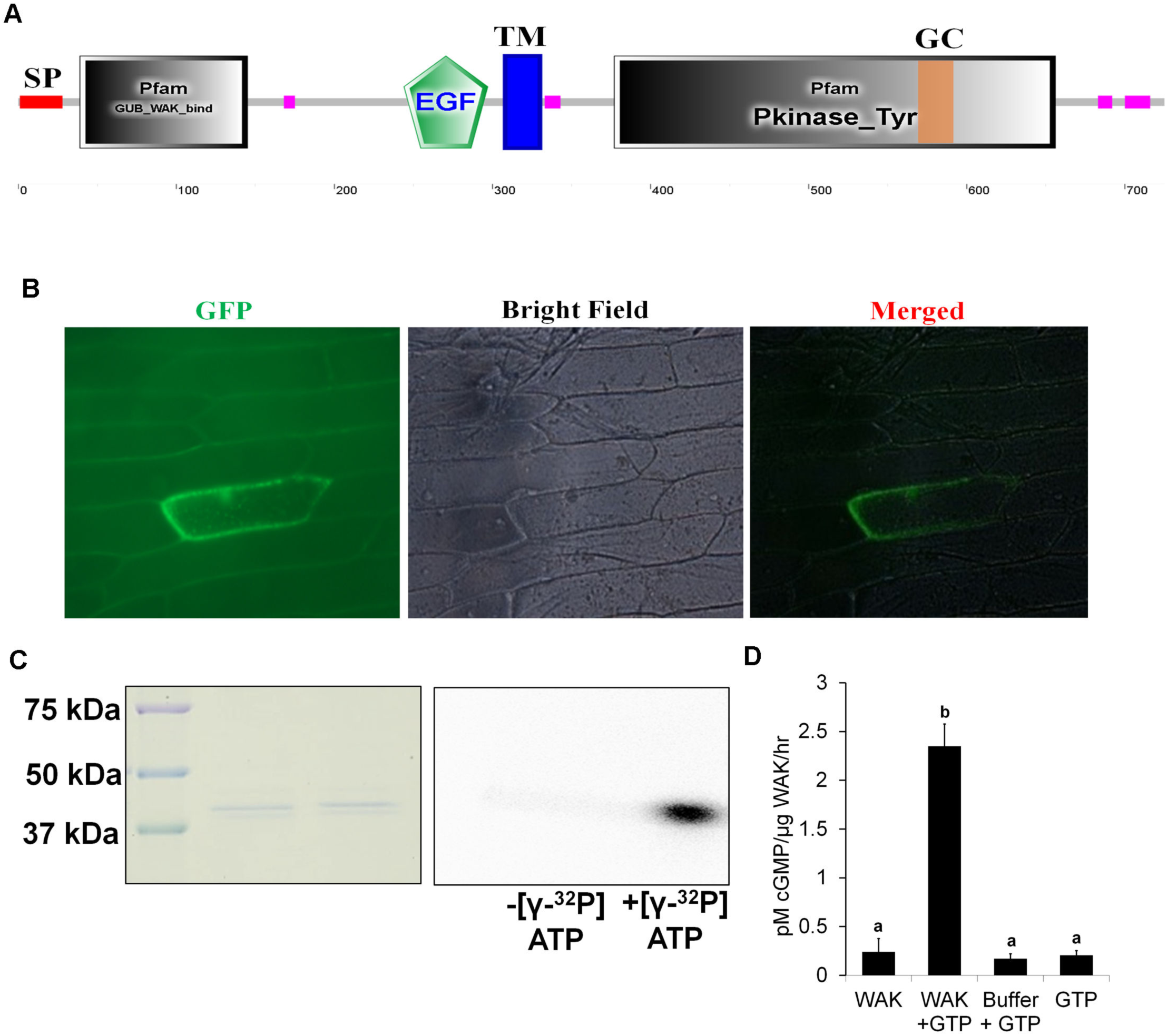
Biochemical characterization and localization of OsWAKL21.2. A. Domain architecture of OsWAKL21.2 using SMART tool (http://smart.embl-heidelberg.de/) (SP: signal peptide, GUB: galacturonan binding domain, EGF: epidermal growth factor like repeat, TM: transmembrane region, Pkinase_Tyr: kinase domain, GC: guanylate cyclase motif).
B. OsWAKL21.2-EGFP localize on the cell membrane in onion peel after transient expression. OsWAKL21.2-EGFP was transiently transformed to onion peel cells using Agrobacterium and peels were visualized after 2 days under epiflourescence microscope. The experiment was repeated three times and similar results were obtained.
C. Kinase assay: Kinase domain of OsWAKL21 cloned and purified from *E. coli* show autophosphorylation activity. 50μg of affinity purified recombinant protein was used for assay with or without radiolabelled ATP. After 1hr, denatured sample was loaded on 10% SDS-PAGE gel. The gel was further subjected to autoradiography and CBB staining. The experiment was repeated three times and similar results were obtained.
D. GC assay: 50μg (in 50 μl) of affinity purified recombinant protein was used for GC assay with or without GTP. After 1hr, 5 μl of the sample was directly used for cGMP quantification. Only GTP and GC buffer + GTP were used as controls. Each bar indicate average and error bar represents SE of three independent experiments. Small letters (a and b) above the bars indicate significant difference with p<0.05.

The biochemical characterization was performed by cloning the intracellular kinase domain of OsWAKL21.2 (OsWAKL21_376-725_) with an N-terminal 6x His tag and expressing it in *E. coli*. The purified cytoplasmic domain of OsWAKL21.2 showed an autophosphorylation activity when incubated with γ-^32^P-ATP indicating that it is an active kinase (Fig. 5C). For the guanylate cyclase activity, the same purified protein was incubated with GTP and cGMP was detected by qualitative and quantitative assays. cGMP was detected only when GTP was incubated with purified OsWAKL21_376-725_ (Fig. 5D, Supplemental Fig. S7C). The rate of cGMP synthesis was 2.1±0.75 pM/µg protein/hr (Fig. 5D) which is comparable to other known plant GCs such as AtPEPR1, AtPSKR1 and AtWAKL10 (Meier et al., 2010, Qi et al., 2010, Kwezi et al., 2011). The biochemical analyses strongly suggests that OsWAKL21.2 is a dual-function enzyme having kinase and GC activity.

### Kinase activity of OsWAKL21.2 is essential for induction of immune responses in rice but not in Arabidopsis

Considering that OsWAKL21.2 is a receptor kinase, we hypothesized that kinase activity of the protein would be required for the induction of immune responses. Based on homology with other plant receptor kinases, we mutated four active site residues (K407, D504, T542, T547) to alanine and generated a kinase-deficient mutant (OsWAKL21.2-kinase deficient or *OsWAKL21.2*-kd). Purified kinase domain of OsWAKL21.2-kd had almost lost kinase activity but it retains GC activity (Supplemental Fig. S8A,B,C). Furthermore, we observed that Agrobacterium-mediated transient overexpression of the full-length *OsWAKL21.2*-kd in rice leaves did not induce rice immune responses such as callose deposition, enhanced tolerance against *Xoo* or increased expression of key defence-related genes (Fig. 6A,B,C, and Supplemental Fig. S9A,B). This indicates that the kinase activity of OsWAKL21.2 is required for induction of immune responses in rice.

**Figure 6:**
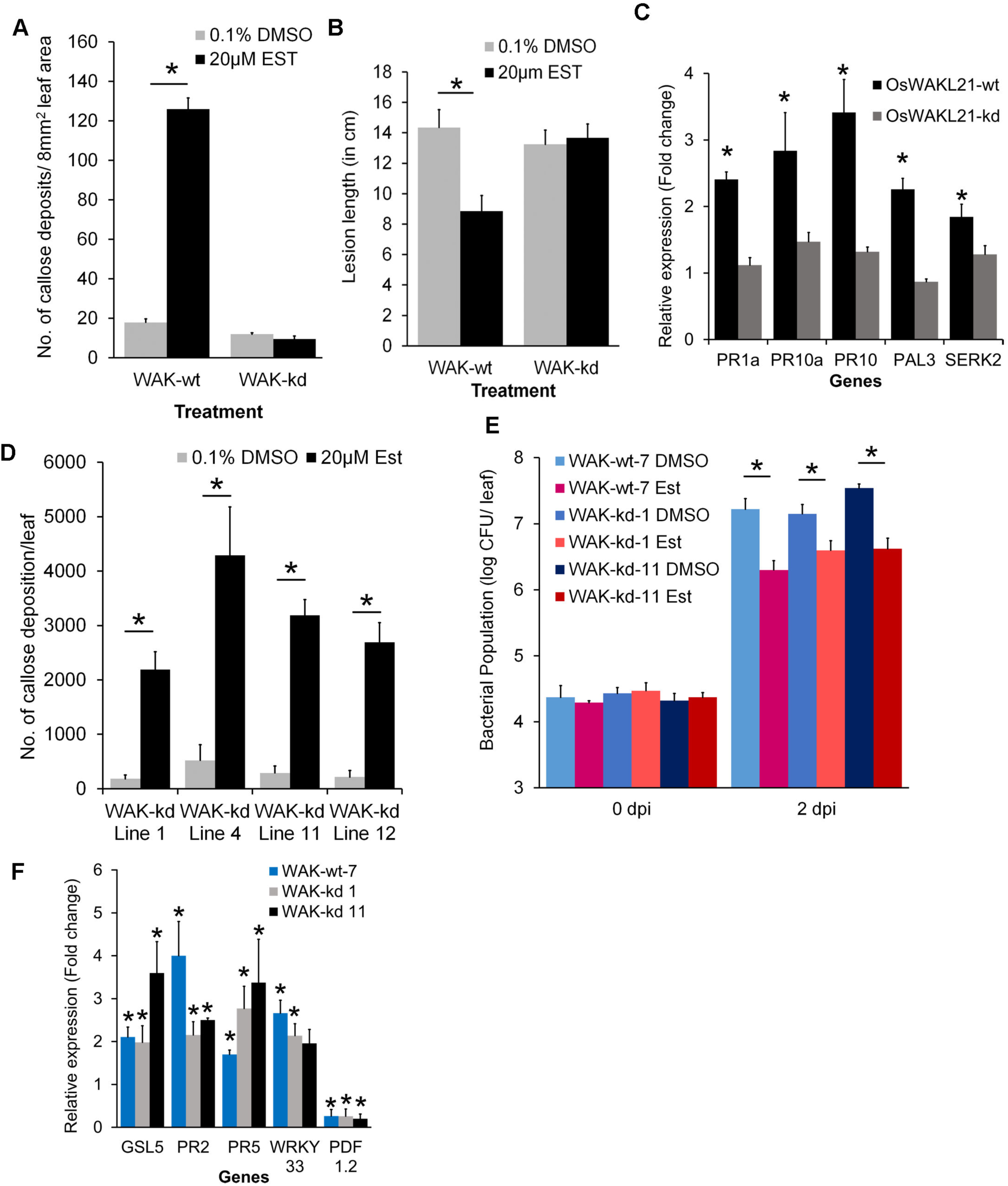
Kinase activity of OsWAKL21.2 is required for induction of immune responses in rice but not in Arabidopsis. A. Quantification of callose deposition after transient overexpression of either wild type *OsWAKL21.2* (WAK-wt) or kinase deficient mutant of *OsWAKL21.2* (*OsWAKL21.2*-kd or WAK-kd) in rice leaves. Each bar represents average and error bar represents SE of at least 12 leaves per treatment in an experiment
B. Lesion lengths after 10 days of *Xoo* pin prick inoculation when *OsWAKL21.2* or *OsWAKL21.2*-kd was transiently overexpressed prior to infection by *Xoo*. Each bar represents average and error bar represents SE of lesion length in 20-30 leaves in an experiment.
C. Relative expression of key defence related genes after transient overexpression of either *OsWAKL21.2* or *OsWAKL21.2*-kd in rice leaves. For each gene, transcript level of uninduced condition (treatment with Agrobacterium carrying WAK-wt or WAK-kd with 0.1% DMSO) was considered as 1 and was compared to induced condition (treatment with Agrobacterium carrying WAK-wt or WAK-kd with 20µM estradiol). Each bar represents average fold change and error bars indicate SE in three independent experiments (n=12 in each experiment).
D. Quantification of callose deposition in leaves of four different *OsWAKL21.2*-kd Arabidopsis transgenic lines (lines 1, 4, 11 and 12) treated with either 20µM β-estradiol (inducer) or 0.1% DMSO (control). Each bar represents average and error bar represents SE of three leaves in an experiment.
E. Effect of ectopic expression of *OsWAKL21.2*-kd on growth of *Pst* DC3000 after subsequent infection. Leaves of wild type *OsWAKL21.2* (WAK-wt) and two different *OsWAKL21.2*-kd Arabidopsis transgenic lines (lines 1 and 11) were infiltrated with either 20µM β-estradiol (inducer) or 0.1% DMSO (control) and were subsequently inoculated with *Pst* DC3000,12hr post infiltration. Each bar represents average and error bar represents SE of five leaves in each sample.
F. Effect of ectopic expression of *OsWAKL21.2*-kd on expression of key defence related *OsWAKL21.2* responsive genes in transgenic Arabidopsis lines. Expression in 0.1% DMSO treated leaves was considered as 1 and relative expression in 20µM estradiol treated leaves was calculated with respect to it. Each bar represents average fold change and error bars indicate SE in three independent experiments (n=3 in each experiment). In C and F, *OsActin1* and *AtActin2* were used respectively as internal control for qRT-PCR. The relative fold change was calculated by using 2^−ΔΔ^ method. Similar results were obtained in three different experiments in A, B, D and E. If the significant difference was observed, asterisk (*) represents significant difference with p<0.05 with respect to uninduced condition.

In order to further investigate the role of the kinase activity of OsWAKL21.2 in the induction of plant immune responses, we generated transgenic Arabidopsis lines expressing *OsWAKL21.2*-kd. Interestingly, we observed that the ectopic expression of *OsWAKL21.2*-kd in Arabidopsis caused an increase in callose deposition (Fig. 6D, Supplemental Fig. S9C,D). Similar results were observed in four different transgenic lines. In Arabidopsis, the ectopic expression of *OsWAKL21.2*-kd showed enhanced tolerance towards *Pst* DC3000 and also changed the expression of defence-related genes in a similar pattern as *OsWAKL21.2* (Fig. 6E,F). As mentioned above, this mutant did not induce immune responses in rice, indicating that the kinase activity of OsWAKL21.2 is vital for the induction of immune responses in rice but not in Arabidopsis.

### GC activity of *OsWAKL21.2* is required for induction of immune responses in Arabidopsis but not in rice

Owing to the fact that the kinase-deficient mutant of *OsWAKL21.2* induced immune responses in Arabidopsis, we decided to investigate whether the GC activity of OsWAKL21.2 might have role in induction of immune responses in Arabidopsis. In order to test this hypothesis, we initially induced the expression of *OsWAKL21.2* in Arabidopsis in the presence of a GC inhibitor LY83583 and observed that the GC inhibitor attenuates *OsWAKL21.2* and *OsWAKL21.2*-kd induced callose deposition in Arabidopsis (Supplemental Fig. S10). In order to confirm this, we generated a mutant of OsWAKL21.2 that lacked the GC activity (*OsWAKL21.2*-GC Deficient or *OsWAKL21.2*-gcd) but retained the kinase activity (Supplemental Fig. S8A,B,C) (Ma et al., 2012). Ectopic expression of *OsWAKL21.2*-gcd did not induce either callose deposition or enhanced tolerance towards *Pst* DC3000 (Fig. 7A,B, and Supplemental Fig. S9C,D). Furthermore, *OsWAKL21.2*-gcd failed to significantly alter the expression of most of the defence-related genes that are differentially regulated by *OsWAKL21.2* in Arabidopsis (Fig. 7C). Ectopic expression of *OsWAKL21.2* in Arabidopsis leaves also enhances *in planta* cGMP level which was not observed when *OsWAKL21.2-gcd* was expressed in transgenic Arabidopsis plants (Supplemental Fig. S11A,B,C). However, transient overexpression of *OsWAKL21.2*-gcd induces immune responses in rice that were similar to the ones induced by the wild-type OsWAKL21.2 (Fig. 7D,E,F, Supplemental Fig. S9A,B). These observations clearly indicated that the GC activity of OsWAKL21.2 is essential for induction of immune responses in Arabidopsis but not in rice.

**Figure 7:**
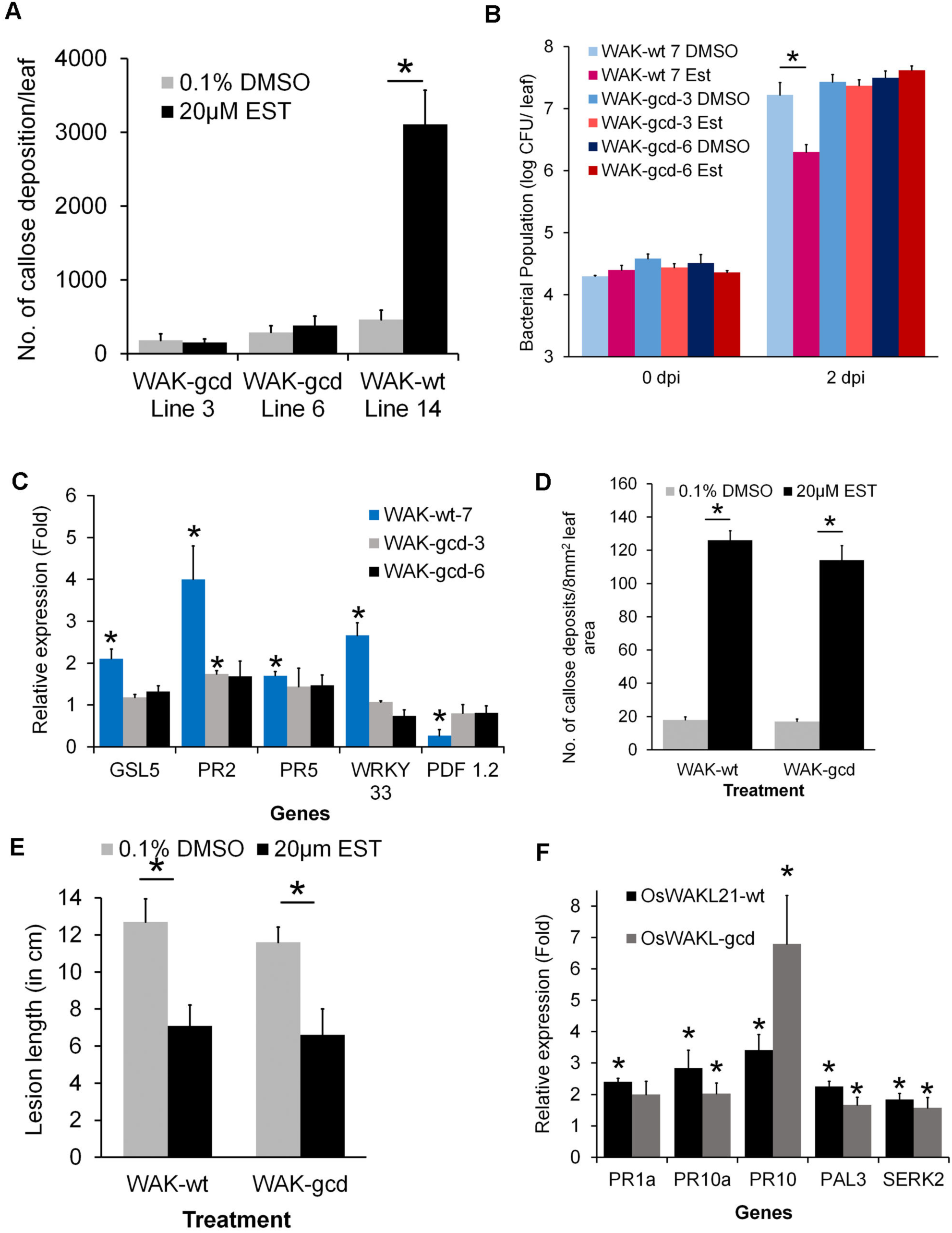
GC activity of OsWAKL21.2 is required for induction of immune responses in Arabidopsis but not in rice. A. Quantification of callose deposition in leaves of two different Arabidopsis transgenic lines (lines 3 and 6) expressing GC deficient *OsWAKL21.2* (*OsWAKL21.2*-gcd or WAK-gcd) that were treated with either 20µM β estradiol (inducer) or 0.1% DMSO (control). Each bar represents average and error bar represents SE of three leaves in an experiment.
B. Effect of ectopic expression of *OsWAKL21.2*-gcd on growth of *Pst* DC3000 after subsequent infection. Leaves of wild type *OsWAKL21.2* (WAK-wt) and two different *OsWAKL21.2*-gcd Arabidopsis transgenic lines (lines 3 and 6) were infiltrated with either 20µM β-estradiol (inducer) or 0.1% DMSO (control) and were subsequently inoculated with *Pst* DC3000, 12hr post infiltration. Each bar represents average and error bar represents SE of five leaves in each sample.
C. Effect of ectopic expression of *OsWAKL21.2*-gcd on expression of key defence related *OsWAKL21.2* induced genes in transgenic Arabidopsis lines. Expression in 0.1% DMSO treated leaves was considered as 1 and relative expression in 20µM estradiol treated leaves was calculated with respect to it. Each bar represents average fold change and error bars indicate SE in three independent experiments (n=3 in each experiment).
D. Quantification of callose deposition after transient overexpression of either wild type (WAK-wt) or WAK-gcd in rice leaves. Each bar represents average and error bar represents SE of at least 12 leaves per treatment in an experiment.
E. Lesion lengths after 10 days of *Xoo* pin prick inoculation when *OsWAKL21.2* or *OsWAKL21.2*-gcd was transiently overexpressed prior to infection by *Xoo*. Each bar represents average and error bar represents SE of lesion length in 20-30 leaves in an experiment.
F. Relative expression of key defence related genes after transient overexpression of either *OsWAKL21.2* or *OsWAKL21.2*-gcd in rice leaves. For each gene, transcript level of uninduced condition (treatment with Agrobacterium carrying WAK-wt or WAK-gcd with 0.1% DMSO) was considered as 1 and was compared to induced condition (treatment with Agrobacterium carrying WAK-wt or WAK-gcd with 20µM estradiol). Each bar represents average fold change and error bars indicate SE in three independent experiments (n=12 in each experiment). In C and F, *AtActin2* and *OsActin1* were used respectively as internal control for qRT-PCR. The relative fold change was calculated by using 2^−ΔΔ^ method. Similar results were obtained in three different experiments in A, B, D and E. If the significant difference was observed, asterisk (*) represents significant difference with p<0.05 with respect to uninduced condition.

### *OsWAKL21.2* possibly induces the JA pathway in rice while it activates SA pathway in Arabidopsis

The results in this study indicated that kinase activity of OsWAKL21.2 is required to induce rice immune responses and that the GC activity is required for induction of Arabidopsis immune responses. Our previous report indicated that the JA pathway is activated in rice leaves after treatment with LipA (Ranjan et al., 2015). We selected a subset of ten genes that were earlier predicted to be associated with the JA pathway in rice and were found to be upregulated after 12hr of LipA infiltration (Ranjan et al., 2015). We tested the expression of these 10 genes and observed that 8 out of 10 genes showed significant upregulation after *OsWAKL21.2* overexpression (Fig. 8A). This indicates that overexpression of *OsWAKL21.2* in rice enhances expression of JA pathway related genes.

**Figure 8:**
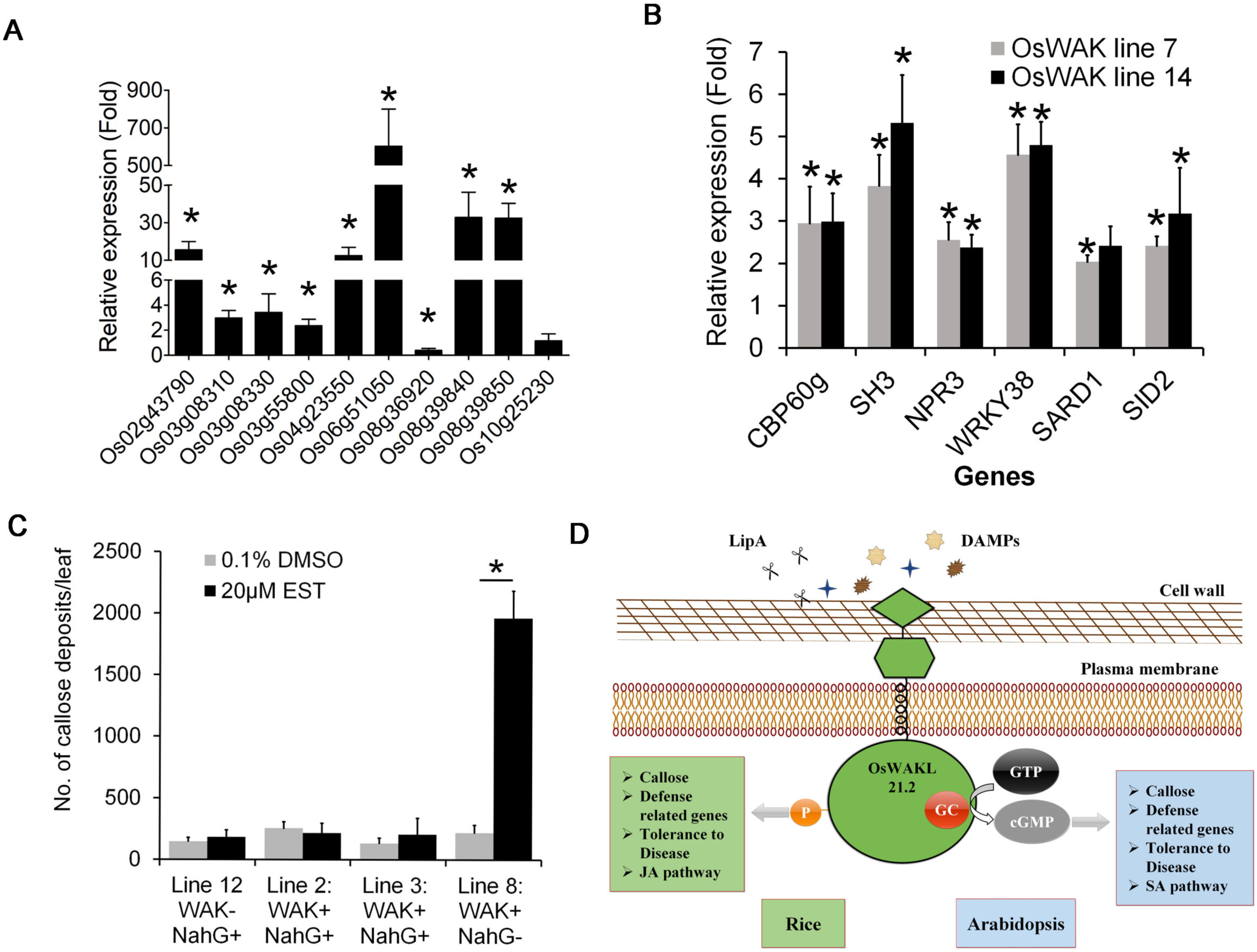
*OsWAKL21.2* induces expression of JA pathway related genes in rice while it activates SA pathway related genes in Arabidopsis. A. Relative expression of ten JA pathway related genes after transient overexpression of *OsWAKL21.2* in rice leaves. These genes include three ZIM domain-containing proteins (LOC_Os03g08310, LOC_Os03g08330 and LOC_Os10g25230), two lipoxygenases (LOC_Os08g39840 and LOC_Os08g39850), one allene oxide synthase (AOS, LOC_Os03g55800), one basic helix loop helix transcription factor (RERJ1, LOC_Os04g23550), one ethylene-responsive transcription factor (ERF, LOC_Os02g43790), one chitinase (PR3, LOC_Os06g51050) and an AP2 domain-containing transcription factor (LOC_Os08g36920). For each gene, transcript level of uninduced condition (treatment with Agrobacterium carrying WAK-wt with 0.1% DMSO) was considered as 1 and was compared to induced condition (treatment with Agrobacterium carrying WAK-wt with 20µM estradiol). Each bar represents average fold change and error bars indicate SE in three independent experiments (n=12 in each experiment). *OsActin1* was used as internal control. The relative fold change was calculated by using 2^−ΔΔCt^ method. Asterisk (*) represents significant difference with p<0.05 with respect to uninduced condition.
B. Effect of ectopic expression of *OsWAKL21.2* on expression of SA pathway related genes in transgenic Arabidopsis lines. Expression in 0.1% DMSO treated leaves was considered as 1 and relative expression in 20µM estradiol treated leaves was calculated with respect to it. Each bar represents average fold change and error bars indicate SE in three independent experiments (n=3 in each experiment). *AtActin2* was used as internal control for qRT-PCR. The relative fold change was calculated by using 2^−ΔΔ^ method. Asterisk (*) represents significant difference with p<0.05 with respect to uninduced condition.
C. Quantification of callose deposits in Arabidopsis crossing lines expressing *NahG* and *OsWAKL21.2* (line 2 and 3) or either one of those (*NahG* only: line 12, WAK-wt only: line 8). Leaves were treated with either 20µM β-estradiol (inducer) or 0.1% DMSO (control). Each bar represents average and error bar represents SE of three leaves in an experiment. Asterisk (*) represents significant difference with p<0.05 with respect to uninduced condition.
D. Modal depicting mechanistic role of OsWAKL21.2 in induction of immune responses in rice and Arabidopsis. OsWAKL21.2 likely perceive cell wall damage caused after LipA treatment in rice. Upon overexpression in rice, OsWAKL21.2 induces rice immune responses via its kinase activity. Upon ectopic expression in Arabidopsis transgenic lines, OsWAKL21.2 induce Arabidopsis immune responses by its GC activity.

The results above (Fig. 4D) suggested that expression of SA related genes was enhanced after ectopic expression of *OsWAKL21.2* in Arabidopsis. We further tested the expression of some more SA pathway related Arabidopsis genes (*AtSID2*, *AtCBP60g, AtSARD1, AtSH3, AtNPR3* and *AtWRKY38*) after ectopic expression of *OsWAKL21.2* and observed significantly enhanced expression of these genes (Fig. 8B). In order to validate the role of the SA pathway in *OsWAKL21.2* induced immune responses in Arabidopsis, we made the crosses between *OsWAKL21.2* transgenic lines with *NahG* transgenic lines that do not accumulate SA (Delaney et al., 1994). Transgenic offspring lines that express both *OsWAKL21.2* and *NahG* did not show enhanced callose deposition while sister lines that expressed only *OsWAKL21.2* showed enhanced callose deposition after treatment with estradiol (Fig. 8C, Supplemental Fig. S12). This observation indicated that *OsWAKL21.2* induces immune responses in Arabidopsis via activation of the SA pathway.

## Discussion

CWDEs are important virulence factors secreted by microbial plant pathogens. *Xoo* secretes numerous CWDEs to degrade the rice cell wall and treatment of rice with *Xoo* secreted purified CWDEs such as Cellulase A (ClsA), Cellobiosidase (CbsA) and Lipase/esterase (LipA) leads to activation of plant immune responses (Jha et al., 2007). Earlier we have shown that the biochemical activity of LipA is required for the induction of rice immune responses (Aparna et al., 2009). This indicates that the rice plant is capable of recognizing cell wall degradation products as DAMPs and further induce immune responses. The molecular players involved in the perception of cell wall damage caused by CWDEs in rice is yet to be deciphered. To discern the functions involved in LipA induced immune responses, we performed transcriptome analyses at various time points following LipA treatment. Comparison with online available microarray data indicates a handful of genes that are commonly upregulated following LipA or *Xoo* treatment. One such gene was the second splice variant of a rice wall-associated kinase-like gene 21 (*OsWAKL21.2*). The wall-associated kinase (WAK) is the only gene family known to recognize plant cell wall-derived DAMPs (Kohorn, 2015). Our study suggests that the expression of *OsWAKL21.2* is enhanced after treatment of rice leaves with either LipA or *Xoo* but not after treatment with a LipA mutant of *Xoo*. This indicates that the increase in *OsWAKL21.2* expression after *Xoo* treatment is specifically because of the presence of LipA in *Xoo*. We also observed that it is a membrane localized receptor kinase having *in vitro* kinase and GC activity.

Downregulation of some WAK gene family members in rice such as *OsWAK14, OsWAK91, OsWAK92* or *Xa4-WAK* have been reported to enhance the susceptibility of rice plants towards subsequent infection (Delteil et al., 2016, Hu et al., 2017). We downregulated the expression of *OsWAKL21.2* in rice leaves using VIGS. Although downregulation of *OsWAKL21.2* did not alter susceptibility against *Xoo*, it attenuated LipA induced tolerance to *Xoo* and callose deposition in rice indicating that it is a key intermediate of signaling activated after LipA treatment. Since optimal expression of *OsWAKL21.2* is essential for LipA induced immune responses, it might be an upstream component in signalling activated following LipA treatment.

Treatment of rice leaves with LipA leads to callose deposition, activation of JA pathway, enhanced expression of some defence related genes and enhanced tolerance against subsequent *Xoo* infection (Jha et al., 2007, Ranjan et al., 2015). Callose deposition is a hallmark of the immune response that is observed after treatment of the plant tissue with CWDEs (including LipA) or DAMPs (Jha et al., 2007, Galletti et al., 2008). We also observed that the overexpression of *OsWAKL21.2* in rice and ectopic expression in Arabidopsis leaves leads to the fortification of the cell wall in the form of callose deposition. Activation of the immune response leads to an increased tolerance towards subsequent infection in plants. We also observed that OsWAKL21.2 induced immune responses lead to enhanced tolerance against subsequent bacterial infection in rice and Arabidopsis. Overexpression of several other WAKs such as *OsWAK1* (Li et al., 2009), *OsWAK25* (Harkenrider et al., 2016), *OsWAK14*, *OsWAK91* or *OsWAK92* (Delteil et al., 2016), *AtWAK2* (Kohorn et al., 2009), *AtWAK1* (Brutus et al., 2010), and Ta-*WAKL4* (Saintenac et al., 2018) has been reported to enhance tolerance towards subsequent infections in different plant species. Immune responses are usually correlated with enhanced expression of defence-related genes. The overexpression of *OsWAKL21.2* in the mid-vein of rice leaves enhanced the expression of five defence-related and LipA responsive genes. The key defence-related genes upregulated by *OsWAKL21.2* overexpression include *OsPR1a* (Park et al., 2008), *OsPR10a* (Bai et al., 2011), *OsPR10* (Harkenrider et al., 2016), *OsSERK2* (Chen et al., 2014) and *OsPAL3* (Chen et al., 2018) which are well categorized as defence-related genes implicated in tolerance against *Xoo.* Interestingly, four of these five key defence genes (except *OsPR1a*) that are upregulated by *OsWAKL21.2* overexpression are also upregulated after 12hr of LipA treatment in a microarray that was earlier done in our lab (Ranjan et al., 2015). Overexpression of *OsWAKL21.2* also enhances the expression of most of the tested LipA responsive genes (7/10) and most of the tested JA pathway related LipA responsive genes (8/10) indicating that the overexpression of *OsWAKL21.2* partially mimics LipA treatment condition. These results establish that the overexpression of *OsWAKL21.2* in rice mimics the LipA treatment indicating *OsWAKL21.2* could be a major upstream component in the signaling process that is activated after cell wall damage caused by LipA.

Ectopic expression of *OsWAKL21.2* leads to enhanced expression of the SA responsive genes such as *AtPR2*, *AtPR5* and *AtWRKY33* and downregulation of the JA responsive gene, *AtPDF1.2* indicating that *OsWAKL21.2* likely activates the SA pathway in Arabidopsis. We observed enhanced expression of several other SA biosynthesis-, regulation- and response-related genes in Arabidopsis (*AtSID2, AtSARD1, AtCBP60G, AtNPR3, AtWRKY33, AtWRKY38* and *AtSH3*) following *OsWAKL21.2* ectopic expression (Janda and Ruelland, 2015). We also found that the transgenic plants expressing *OsWAKL21.2* and *NahG* together did not show callose deposition, demonstrating that SA accumulation is required for *OsWAKL21.2* induced immune response in Arabidopsis. These outcomes also explain the enhanced tolerance towards *Pst* DC3000, as an activation of the SA pathway in Arabidopsis leads to increased tolerance towards *Pst* DC3000 (Xin and He, 2013). Activation of SA pathway in Arabidopsis enhances expression of biotic stress-responsive callose synthase *AtGSL5* (Dong et al., 2008) which was also upregulated following ectopic expression of *OsWAKL21.2*. The results indicate that *OsWAKL21.2* when expressed ectopically in Arabidopsis acts as a defence gene and activates SA pathway-mediated immune responses. Some members of the WAK gene family in Arabidopsis such as *AtWAK1, AtWAK2, AtWAK3, AtWAK5* and *AtWAKL10* are known as SA responsive genes as treatment with SA leads to the enhanced expression of these genes indicating correlation of SA pathway and WAKs in Arabidopsis (He et al., 1998, He et al., 1999, Meier et al., 2010).

Ligand binding onto receptor kinases triggers phosphorylation that is further conveyed downstream via phosphorylation by/of kinases and their targets (Macho and Zipfel, 2014). Few receptor kinases such as AtBRI1, AtPSKR1, AtPEPR1, AtWAKL10 and HpPEPR1 are also known to possess dual enzymatic activity i.e. they possess GC activity along with kinase activity (Meier et al., 2010, Ma et al., 2012, Swiezawska et al., 2015, Gehring and Turek, 2017, Swiezawska et al., 2017). OsWAKL21.2 also possess such dual activity which is comparable with other plant GCs. Treatment with a GC inhibitor and mutations in active site residues of the GC motif showed that the GC activity of OsWAKL21.2 is required to induce immune responses in Arabidopsis but not in rice. GCs convert GTP to cGMP which acts as a secondary signaling molecule (Gehring and Turek, 2017). Overexpression of *AtBRI1*, *AtPSKR1* and *AtPEPR1* (having GC activity) in Arabidopsis leads to a partial increase in cytoplasmic cGMP concentrations (Gehring and Turek, 2017) which was also observed following ectopic expression of *OsWAKL21.2* in Arabidopsis. Some of the moonlighting kinases such as AtPEPR1, AtBRI1 and AtPSKR1 are already known for their direct or modulatory role in Arabidopsis immune responses (Igarashi et al., 2012, Lozano-Durán and Zipfel, 2015). AtPEPR1 is receptor of DAMP (Pep’s) and it’s GC activity is required for activation of immune responses (Ma et al., 2012). *AtWAKL10* has also been predicted as a defence-related gene that belong to similar gene family as *OsWAKL21.2*. These observations testify the possible involvement of GCs in Arabidopsis immune response. We have observed that in rice, OsWAKL21.2 requires the kinase activity to induce immunity, whereas, in Arabidopsis, it requires the GC activity. This does not rule out the possible role of GC activity of OsWAKL21 in rice as it might be involved in some other functions not studied here.

CWDEs secreted by *Xoo* cause degradation of the rice cell wall that leads to the release of cell wall derived DAMPs. These DAMPs, in turn, induce rice immune responses, but the mechanisms by which these DAMPs are perceived and recognized are obscure. Employing a variety of analyses, we have found that the rice receptor kinase *OsWAKL21.2* is required for the activation of plant immune responses post-LipA treatment. This suggests that *OsWAKL21.2* could be either a receptor or a co-receptor for cell wall damage and possibly the first DAMP receptor identified in rice. Overexpression of *OsWAKL21.2* in plants induces immune responses and enhances tolerance towards hemibiotrophic pathogens. We observed that this receptor kinase is a moonlighting kinase having *in vitro* GC activity along with kinase activity making it one of the few moonlighting kinases known in plants and the first one in rice. An interesting observation about *OsWAKL21.2* is that for the induction of immune responses in rice, the kinase activity is required, but in Arabidopsis, the GC activity is needed. Fig. 8D represents a mechanistic model of the role of OsWAKL21.2 in the induction of immune responses in rice and Arabidopsis. Future studies would be aimed at identifying interacting partners of OsWAKL21.2 that are involved in elaboration of LipA induced immune responses. Furthermore, the possibility of using this gene to provide enhanced tolerance to bacterial pathogens in a variety of crops including monocots and dicots can be explored.

## Materials and Methods

### Plant materials and growth conditions

Rice (*Oryza sativa* ssp. *indica*) variety TN1 (Taichung native 1) which is susceptible to *Xanthomonas oryzae* pv. *oryzae* (*Xoo*) was used for plant experiments. All the rice experiments were performed in either the growth chamber (12hr Day//Night) or greenhouse at 28°C. *Arabidopsis thaliana* ecotype Columbia (Col-0) and *NahG* lines were used for Arabidopsis experiments. Transgenic lines were generated using the floral dip method (Clough and Bent, 1998). Transgenic plants were selected by adding hygromycin and/or kanamycin (*NahG* lines) to the final concentration of 20µg/ml or 50µg/ml respectively. Plants were maintained in growth chamber at 22°C day and 18°C night temperature at about 70% humidity and with 12hr day/night cycle. Leaves of 4-5-week-old plants that are in rosette state were used for experiments.

### Bacterial cultures

*Xoo* wild type strain BXO43 (lab isolate) was used as a rice pathogen. The LipA mutant (BXO2001) of *Xoo* (BXO43) and its complemented strain (BXO2008) was also used in this study (Rajeshwari et al., 2005). *Pseudomonas syringae* pv. *tomato* DC3000 (*Pst* DC3000) was used as an Arabidopsis pathogen. Transient transformation in rice and floral dip of Arabidopsis was performed using *Agrobacterium tumefaciens* strain LBA4404. *E. coli* BL21-AI was used for recombinant protein expression for biochemical assays.

### LipA purification from *Xoo* culture supernatent

*Xoo* BXO2008, a LipA overproducing strain derived from BXO2001 was used for LipA overproduction and purification and LipA was purified by the protocol described previously (Aparna et al., 2007). The purity and activity of the enzyme was tested by running on a SDS-PAGE gel and activity on tributyrin containing plates respectively.

### Microarray analysis

The leaf treatment and microarray analysis was performed as described previously (Ranjan et al., 2015). RNA was isolated from 25-30 leaves after 30min or 2hr of treatment either with LipA (0.5mg/ml) or buffer. Processed data and ‘.cel’ files were also submitted to gene expression omnibus (GEO-NCBI, Acc. No. GSE53940). RMA and PLIER16 algorithms were used for analysis and probes showing significant differential expression (FC ≥ 1.5-fold and p<0.05) in both analyses were considered as differentially expressed genes.

### Vector construction and site-directed mutagenesis

Gateway^TM^ cloning technology was used for cloning. *OsWAKL21.2* was amplified using rice cDNA and cloned into pENTR-D-TOPO (Invitrogen™). The gene was subcloned using LR clonase reaction (Invitrogen^TM^) into pMDC7 plasmid (Curtis and Grossniklaus, 2003) for plant expression studies and in pH7FWG2 plasmid (Karimi et al., 2002) for localization experiments. In pMDC7, the target gene sequence is cloned downstream to XVE promoter, which is 17-β-estradiol inducible. 20 μM of 17-β-estradiol (Sigma Aldrich) was used in all overexpression studies as an inducer while 0.1% DMSO was used as a control (uninduced condition). Kinase domain OsWAKL21_376-725_ was cloned into bacterial expression vector pDEST17 (Invitrogen) and transformed into *E. coli* BL21-AI for recombinant protein expression. The constructs in pENTR-D-TOPO were used for site-directed mutagenesis (Zheng et al., 2004). The mutant versions were then transferred into desired destination vectors using LR clonase reaction. All the clones and mutations were confirmed using Sanger sequencing. All the plant expression constructs were introduced into *Agrobacterium tumefaciens* strain LBA4404. LBA4404:XVE_pro_:*OsWAKL21.2*, LBA4404:XVE_pro_:*OsWAKL21.2*-kd and LBA4404:XVE_pro_:*OsWAKL21.2*-gcd were used for transient transformation in rice and for generation of Arabidopsis transgenic lines.

### Callose deposition assay in rice and Arabidopsis

For callose deposition assay in rice, 12-14 days old leaves were used for Agrobacterium-mediated transient transformation (Jha et al., 2010, Pillai et al., 2018). The suspension was infiltrated in third rice leaf using a needleless 1ml syringe with inducer [20µM 17-β-estradiol; (Est), Sigma-Aldrich] or control (0.1% DMSO). Leaves collected for callose deposition were stained with aniline blue according to Millet et al. (2010) (Millet et al., 2010). Callose deposition was visualized under blue light (excitation wavelength 365nm) in ECLIPSE Ni-E, epifluorescence microscope (Nikon, Japan) with 10X magnification. Eight images (∼1mm^2^ each) were captured from each leaf from the zone of infiltration and proximal region. The number of callose deposits in all eight images for a leaf was added to get callose deposition per leaf (per 8mm^2^). Average was calculated for 10-12 leaves for each treatment.

For callose deposition in Arabidopsis transgenic plants, similar size of rosette stage leaves were infiltrated either with 100µl of 0.1%DMSO or 20µM estradiol (inducer) using the needleless 1.0 ml syringe. After 12hr, leaves were collected and stained for callose deposition and observed under the microscope as mentioned above for rice. Nearly 40-50 images per leaf were captured and the number of callose deposits in each image was added to get number of callose deposits in one leaf. For each sample average was calculated for 3 such leaves obtained from three separate plants.

### Virulence assay in rice and Arabidopsis

About 60 days old TN1 rice plants were used for infection of *Xoo*. For transient overexpression in rice mid-vein, 200µl actively growing Agrobacterium (LBA4404) resuspended in 10mM MES + 10mM MgCl_2_ + 200µM acetosyringone (final OD 0.8) [either with (20µM 17-β-estradiol) or without (0.1% DMSO) inducer] was injected using a 1.0 ml syringe. After 24hr, about 1cm above Agrobacterium injection site, the mid-veins of leaves were pin-pricked with needle touched to fresh *Xoo* colony. Lesion length caused by *Xoo* was measured after 10 days of *Xoo* infection.

*Pseudomonas syringae* pv. *tomato* (*Pst* DC3000) was used for infection in Arabidopsis leaves. Similar size leaves from five different rosette stage plants were infiltrated with either 0.1% DMSO or 20µM estradiol. After 12hr, leaves were infected with actively growing culture of *Pst* DC3000 (Diluted to OD 0.02) by infiltration using a needleless 1.0 ml syringe. Colony forming unit (CFU) at 0dpi (days post infection) and 2dpi was calculated.

### Downregulation of *OsWAKL21.2* using virus-induced gene silencing (VIGS)

Virus-induced gene silencing was used for Agrobacterium-mediated transient downregulation of *OsWAKL21.2* in rice. Three RNAi constructs of different length from unique 5’-end of *OsWAKL21.2* were cloned in pRTBV-MVIGS (Purkayastha et al., 2010). Downregulation was performed with a modified protocol mentioned previously (Purkayastha et al., 2010, Kant and Dasgupta, 2017). For callose deposition studies, just germinated rice seedlings (1 day old) were dipped in activated Agrobacterium culture (in 10mM MES+10mM MgCl_2_+200µM acetosyringone) for 24hr (Supplemental Fig. S3). 10 days after Agrobacterium treatment, the third leaf of each plant was infiltrated with LipA using a needleless syringe (0.5mg/ml) (at least 40 leaves for each Agrobacterial strain). After 16hr, a small piece (∼1.5cm) of each leaf around the zone of infiltration was collected for callose deposition while the rest of the leaf piece was stored for transcript/protein quantification. Each leaf was collected separately for callose and transcript/protein quantification and labelled. Callose deposition was observed as mentioned above for callose deposition assay. Rest of the part of 4-5 leaves that showed either low or high callose deposition were pooled and RNA/protein was isolated from those pooled leaves for qRT-PCR or Western blotting.

For virulence assay after downregulation of *OsWAKL21.2*, mid-veins of 60 days old rice plants were injected with 200µl Agrobacterium containing the VIGS construct along with either buffer or LipA (0.5mg/ml) (n>40). After 24hr, mid-veins of 10 leaves were collected (3cm each) for *OsWAKL21.2* transcript/protein quantification while remaining 20-30 leaves were infected with a freshly growing colony of *Xoo* as mentioned earlier. Lesion length caused by *Xoo* was measured after 10 days of infection.

### Purification of recombinant protein and *in vitro* biochemical assays

The recombinant kinase domain of OsWAKL21.2, OsWAKL21*.2*_376-725_ with 6X-His tag was cloned, expressed and purified from *E. coli* BL21-AI. 50µg of purified recombinant protein was used for kinase or GC assay in a 50µl reaction. The purified protein was incubated with 10µCi of [γ-^32^P] ATP in kinase assay buffer (50mM Tris (pH 7.5), 10mM MgCl_2_, 2mM MnCl_2_, 0.5mM CaCl_2_, 1mM DTT and 20mM ATP) for 1hr at room temperature (Li et al., 2009), run on 10% SDS-PAGE gel and gel was subsequently exposed to phosphoimager screen which was later scanned in phosphoimager (Personal molecular imager, Biorad) instrument.

GC assay was also performed from the same purified recombinant protein in GC assay buffer [50mM Tris (pH 7.5), 2mM MgCl_2_, 1mM MnCl_2_, 0.5mM CaCl_2_, 0.2mM NONOate (Sigma)] modified from the protocol described previously (Meier et al., 2010). The reaction was incubated at 37°C for either 1hr or 12hr. The 1hr reaction was used for quantitative analysis while 12hr reactions were used for qualitative analysis. cGMP produced after 1hr was quantified using cGMP enzyme immunoassay kit (Sigma-Aldrich, Cat. No-CG201) according to manufacturer’s protocol and the data was analyzed using online tool ‘Elisaanalysis’ (https://elisaanalysis.com/app). For qualitative analysis, the resultant product was blotted on nitrocellulose membrane (Amersham, Cat No. RPN203E) and dried in the laminar hood with UV on for 1hr. The nucleotides were further crosslinked to the membrane by keeping in UV transilluminator for 30min. The membrane was blocked, washed and further incubated with anti cGMP antibody (1:1000, Sigma-Aldrich, Cat. No-G4899) and processed as mentioned in Western blot section.

### RNA isolation and gene expression analysis

For qRT-PCR, RNA was isolated by the protocol of Sánchez et al. (2008) with some modifications (Oñate-Sánchez and Vicente-Carbajosa, 2008, Couto et al., 2015). For rice, 10-12 leaf pieces (or mid-vein pieces) were crushed together for each treatment unless mentioned otherwise. For Arabidopsis, three leaf pieces from separate plants were crushed together for each treatment. cDNA was made from 5µg of total RNA [RNA to cDNA EcoDry™ Premix (Oligo dT), (Clontech)] according to the manufacturer’s protocol. qRT-PCR was performed with diluted cDNA using Power SYBR^TM^ Green PCR Master Mix (Thermo Fisher Scientific) in ViiA 7 Real-Time PCR System (Applied Biosystems). Relative expression was calculated in enzyme or 17-β-estradiol treated leaves with respect to mock/control (buffer or 0.1% DMSO) treated leaves. The fold change was calculated using 2^−ΔΔCt^ method (Livak and Schmittgen, 2001). *OsActin1* and *AtActin2* were used as internal control for rice and Arabidopsis respectively. All the primers for qRT-PCR were designed using QuantPrime (Arvidsson et al., 2008).

### Protein isolation and Western blotting

For Western blot the protein was isolated from 10-12 leaf pieces of rice or three leaves of Arabidopsis using the protocol described previously with minor modifications (Rohila et al., 2006). 20µg of total protein was loaded in 10% SDS-PAGE gel for Western blot/Coomassie brilliant blue staining. The protein was transferred to PVDF membrane (Millipore) and processed for blotting. Anti OsWAKL21_376-725_ antibodies were generated in the rabbit in our institute animal house facility and used in dilution of 1:100. HRP tagged anti-Rabbit IgG secondary antibody (Abcam) (dilution 1:50000) was used and the blot was visualized in chemidoc (Vilber Lourmat).

### Localization of OsWAKL21.2

The localization of OsWAKL21.2 was observed by transient transformation of onion peel cell as described previously (Sun et al., 2007). Os*WAKL21.2* was cloned into Gateway compatible vector pH7FWG2 (Karimi et al., 2002) and transformed in onion peel using Agrobacterium-mediated transient transformation. The GFP signal was visualized under GFP filter in ECLIPSE Ni-E, epifluorescence microscope (Nikon, Japan).

### cGMP quantification

cGMP was quantified in leaves of rosette stage transgenic Arabidopsis plants by the method used by, Dubovskaya et al. (2011), Nan et al. (2014) and Chen et al. (2018) with minor modifications (Dubovskaya et al., 2011, Nan et al., 2014, Chen et al., 2018). Six similar sized leaves (total approximate 200mg) were collected from different plants for untreated control (UT). 3-3 similar size leaves from three different plants were infiltrated either with 0.1% DMSO or 20µM estradiol. Two leaves from each plant (total 6 leaves, ∼200mg) were collected for cGMP quantification while the third leaf was used for testing of expression of *OsWAKL21.2*. After 3hr of infiltration, leaves were collected and crushed in a fine powder using liquid nitrogen. The powder was resuspended in 2ml ice cold 6% (v/v) trichloroacetic acid (TCA) and was collected in the 5ml tube. After brief vortexing (10s), tubes were centrifuged twice at 1000g for 15min at 4°C and supernatant was collected each time in the 5ml tube. The aqueous supernatant was washed 7-8 times with water-saturated diethyl ether.

The solvent was evaporated in cold vacuum centrifuge at 4°C (SCANVAC, CoolSafe). cGMP was quantified in the extract using cGMP enzyme immunoassay kit (Sigma-Aldrich, Cat. No-CG201) according to the manufacturer’s protocol. Data were analyzed using the online tool Elisaanalysis (https://elisaanalysis.com/app).

### Analyses of publicly available transcriptome data

Rice microarray data performed after *Xanthomonas oryzae* treatment was obtained from GEO, NCBI (Acc. No. GSE36272). ‘.cel’ files were downloaded, analyzed and processed using expression console (Affymetrix) using RMA based normalization. ‘.chp’ files obtained after analysis were used in TAC software (Transcriptome analysis console v3.0, Affymetrix) for relative expression analysis. Genes that show FC ≥ 1.5-fold with p<0.05 were considered as differentially expressed.

### Statistical analysis

All experiments were independently performed at least thrice. All data represented here indicate mean ± SE (standard error). The results of lesion length, callose deposition and bacterial growth in CFU were analysed by one-way ANOVA (p<0.05) followed by the Tukey-Kramer test. The results of qRT-PCR were analyzed by Student’s *t*-test and the genes that show significanty altered expression (p<0.05) between control and treated were considered as differentially expressed.

## Accession numbers

The PLIER16 and RMA processed microarray data files generated and used in this experiment are submitted to gene expression omnibus (GEO) (https://www.ncbi.nlm.nih.gov/geo/) under the accession number GSE53940. Other publicly available microarray data used in our analysis was harvested from GEO under the accession numbers GSE49242 and GSE36272. Accession numbers of genes referred in this study are provided in supplemental table S5.

## Acknowledgement

We thank Mr. Ramesh P. (CSIR-CCMB) for helping in analysing the microarray data. We thank Dr. Alok K. Sinha (DBT-NIPGR), Dr. Gopaljee Jha (DBT-NIPGR) and Dr. Puran Singh Sijwali (CSIR-CCMB) for their key suggestions in experiments. We also thank Dr. Subhadeep Chatterjee (DBT-CDFD) for providing *NahG* transgenic lines and *Pseudomonas syringae* DC3000 strain.

## Supplemental data

**Supplemental Table S1:** List of probe sets that show differential expression after 2hr of LipA treatment.

**Supplemental Table S2:** List of differentially expressed genes after 2hr and 12hr of LipA treatment.

**Supplemental Table S3:** Frequency of differentially expressed genes after LipA treatment in the microarray data performed after 24hr of *Xanthomonas oryzae* treatment in GEO submission GSE36272.

**Supplemental Table S4:** List of primers used in this study.

**Supplemental Table S5:** Accession numbers of the genes mentioned in this study.

**Supplemental Fig. S1:** Transcriptome profiling of rice leaves after treatment with LipA.

**Supplemental Fig. S2:** Overexpression of *OsWAKL21.2* induces rice immune responses.

**Supplemental Fig. S3:** Methodology for downregulation of *OsWAKL21.2* in rice seedlings using Virus Induced Gene Silencing (VIGS).

**Supplemental Fig. S4:** Transient downregulation of *OsWAKL21.2* in rice.

**Supplemental Fig. S5:** VIGS mediated transient downregulation of *OsWAKL21.2* does not have significant effect on expression of predicted off-targets genes.

**Supplemental Fig. S6:** qRT-PCR and Western blot validation for ectopically expressing *OsWAKL21.2* transgenic Arabidopsis plants.

**Supplemental Fig. S7:** Biochemical characterization of OsWAKL21.2.

**Supplemental Fig. S8:** Biochemical activities of purified kinase domain of mutant versions of OsWAKL21.2.

**Supplemental Fig. S9:** qRT-PCR and Western blot validation of expression of mutant versions of *OsWAKL21.2* by transient transformation in rice and ectopic expression in Arabidopsis transgenic lines.

**Supplemental Fig. S10:** Treatment with GC inhibitor attenuates *OsWAKL21.2* induced callose deposition in transgenic Arabidopsis leaves.

**Supplemental Fig. S11:** Ectopic expression of *OsWAKL21.2* in Arabidopsis enhances *in planta* cGMP level by its GC activity.

**Supplemental Figure S12:** Western blot validation of ectopic expression of *OsWAKL21.2* in Arabidopsis transgenic lines generated after crossing with *NahG* lines.

## References

Albersheim, P. & Anderson-Prouty, A. J. 1975. Carbohydrates, proteins, cell surfaces, and the biochemistry of pathogenesis. Annual Review of Plant Physiology, 26, 31–52.

Aparna, G., Chatterjee, A., Jha, G., Sonti, R. V. & Sankaranarayanan, R. 2007. Crystallization and preliminary crystallographic studies of LipA, a secretory lipase/esterase from Xanthomonas oryzae pv. oryzae. Acta Crystallogr Sect F Struct Biol Cryst Commun, 63, 708–10.

Aparna, G., Chatterjee, A., Sonti, R. V. & Sankaranarayanan, R. 2009. A cell wall-degrading esterase of Xanthomonas oryzae requires a unique substrate recognition module for pathogenesis on rice. Plant Cell, 21, 1860–73.

Arvidsson, S., Kwasniewski, M., Riaño-Pachón, D. M. & Mueller-Roeber, B. 2008. QuantPrime – a flexible tool for reliable high-throughput primer design for quantitative PCR. BMC Bioinformatics, 9, 465.

Bai, W., Chern, M., Ruan, D., Canlas, P. E., Sze-To, W. H. & Ronald, P. C. 2011. Enhanced disease resistance and hypersensitivity to BTH by introduction of an NH1/OsNPR1 paralog. Plant Biotechnology Journal, 9, 205–215.

Brutus, A., Sicilia, F., Macone, A., Cervone, F. & Lorenzo, G. 2010. A domain swap approach reveals a role of the plant wall-associated kinase 1 (WAK1) as a receptor of oligogalacturonides. Proc Natl Acad Sci U S A, 107.

Chen, X., Zuo, S., Schwessinger, B., Chern, M., Canlas, P. E., Ruan, D., Zhou, X., Wang, J., Daudi, A., Petzold, C. J., Heazlewood, J. L. & Ronald, P. C. 2014. An XA21-associated kinase (OsSERK2) regulates immunity mediated by the XA21 and XA3 immune receptors. Mol Plant, 7, 874–92.

Chen, Z., Chen, T., Sathe, A., He, Y., Zhang, X.-B. & Wu, J.-L. 2018. Identification of a Novel Semi-Dominant Spotted-Leaf Mutant with Enhanced Resistance to Xanthomonas oryzae pv. oryzae in Rice. International journal of molecular sciences, 19, 3766.

Claverie, J., Balacey, S., Lemaître-Guillier, C., Brulé, D., Chiltz, A., Granet, L., Noirot, E., Daire, X., Darblade, B., Héloir, M.-C. & Poinssot, B. 2018. The Cell Wall-Derived Xyloglucan Is a New DAMP Triggering Plant Immunity in Vitis vinifera and Arabidopsis thaliana. Frontiers in Plant Science, 9, 1725.

Clough, S. J. & Bent, A. F. 1998. Floral dip: a simplified method for Agrobacterium-mediated transformation of Arabidopsis thaliana. Plant J, 16, 735–43.

Couto, D., Stransfeld, L., Arruabarrena, A., Zipfel, C. & Lozano-Durán, R. 2015. Broad application of a simple and affordable protocol for isolating plant RNA. BMC Research Notes, 8, 154.

Curtis, M. D. & Grossniklaus, U. 2003. A Gateway Cloning Vector Set for High-Throughput Functional Analysis of Genes in Planta. Plant Physiology, 133, 462–469.

De Azevedo Souza, C., Li, S., Lin, A. Z., Boutrot, F., Grossmann, G., Zipfel, C. & Somerville, S. 2017. Cellulose-derived oligomers act as damage-associated molecular patterns and trigger defense-like responses. Plant Physiology.

Delaney, T. P., Uknes, S., Vernooij, B., Friedrich, L., Weymann, K., Negrotto, D., Gaffney, T., Gut-Rella, M., Kessmann, H., Ward, E. & Ryals, J. 1994. A central role of salicylic Acid in plant disease resistance. Science, 266, 1247–50.

Delteil, A., Gobbato, E., Cayrol, B., Estevan, J., Michel-Romiti, C., Dievart, A., Kroj, T. & Morel, J.-B. 2016. Several wall-associated kinases participate positively and negatively in basal defense against rice blast fungus. BMC Plant Biology, 16, 17.

Dong, X., Hong, Z., Chatterjee, J., Kim, S. & Verma, D. P. S. 2008. Expression of callose synthase genes and its connection with Npr1 signaling pathway during pathogen infection. Planta, 229, 87–98.

Dubovskaya, L. V., Bakakina, Y. S., Kolesneva, E. V., Sodel, D. L., Mcainsh, M. R., Hetherington, A. M. & Volotovski, I. D. 2011. cGMP-dependent ABA-induced stomatal closure in the ABA-insensitive Arabidopsis mutant abi1-1. New Phytol, 191, 57–69.

Galletti, R., Denoux, C., Gambetta, S., Dewdney, J., Ausubel, F. M., De Lorenzo, G. & Ferrari, S. 2008. The AtrbohD-Mediated Oxidative Burst Elicited by Oligogalacturonides in Arabidopsis Is Dispensable for the Activation of Defense Responses Effective against <em>Botrytis cinerea</em>. Plant Physiology, 148, 1695–1706.

Gehring, C. & Turek, I. S. 2017. Cyclic nucleotide monophosphates and their cyclases in plant signaling. Frontiers in plant science, 8, 1704.

Gust, A. A., Pruitt, R. & Nürnberger, T. 2017. Sensing Danger: Key to Activating Plant Immunity. Trends in plant science, 22, 779–791.

Harkenrider, M., Sharma, R., De Vleesschauwer, D., Tsao, L., Zhang, X., Chern, M., Canlas, P., Zuo, S. & Ronald, P. C. 2016. Overexpression of Rice Wall-Associated Kinase 25 (OsWAK25) Alters Resistance to Bacterial and Fungal Pathogens. PLoS ONE, 11, e0147310.

He, Z. H., Cheeseman, I., He, D. & Kohorn, B. D. 1999. A cluster of five cell wall-associated receptor kinase genes, Wak1-5, are expressed in specific organs of Arabidopsis. Plant Mol Biol, 39, 1189–96.

He, Z. H., He, D. & Kohorn, B. D. 1998. Requirement for the induced expression of a cell wall associated receptor kinase for survival during the pathogen response. Plant J, 14, 55–63.

Hématy, K., Cherk, C. & Somerville, S. 2009. Host–pathogen warfare at the plant cell wall. Current Opinion in Plant Biology, 12, 406–413.

Hu, K., Cao, J., Zhang, J., Xia, F., Ke, Y., Zhang, H., Xie, W., Liu, H., Cui, Y., Cao, Y., Sun, X., Xiao, J., Li, X., Zhang, Q. & Wang, S. 2017. Improvement of multiple agronomic traits by a disease resistance gene via cell wall reinforcement. 3, 17009.

Hurni, S., Scheuermann, D., Krattinger, S. G., Kessel, B., Wicker, T., Herren, G., Fitze, M. N., Breen, J., Presterl, T. & Ouzunova, M. 2015. The maize disease resistance gene Htn1 against northern corn leaf blight encodes a wall-associated receptor-like kinase. Proceedings of the National Academy of Sciences, 112, 8780–8785.

Igarashi, D., Tsuda, K. & Katagiri, F. 2012. The peptide growth factor, phytosulfokine, attenuates pattern-triggered immunity. Plant J, 71, 194–204.

Jacobs, A. K., Lipka, V., Burton, R. A., Panstruga, R., Strizhov, N., Schulze-Lefert, P. & Fincher, G. B. 2003. An Arabidopsis Callose Synthase, GSL5, Is Required for Wound and Papillary Callose Formation. Plant Cell, 15, 2503–13.

Janda, M. & Ruelland, E. 2015. Magical mystery tour: salicylic acid signalling. Environmental and Experimental Botany, 114, 117–128.

Jha, G., Patel, H. K., Dasgupta, M., Palaparthi, R. & Sonti, R. V. 2010. Transcriptional Profiling of Rice Leaves Undergoing a Hypersensitive Response Like Reaction Induced by Xanthomonas oryzae pv. oryzae Cellulase. Rice, 3, 1–21.

Jha, G., Rajeshwari, R. & Sonti, R. V. 2007. Functional interplay between two Xanthomonas oryzae pv. oryzae secretion systems in modulating virulence on rice. Molecular Plant-Microbe Interactions, 20, 31–40.

Kant, R. & Dasgupta, I. 2017. Phenotyping of VIGS-mediated gene silencing in rice using a vector derived from a DNA virus. Plant cell reports, 36, 1159–1170.

Karimi, M., Inze, D. & Depicker, A. 2002. GATEWAY vectors for Agrobacterium-mediated plant transformation. Trends Plant Sci, 7, 193–5.

Kohorn, B. D. 2015. Cell wall-associated kinases and pectin perception. Journal of experimental botany, 67, 489–494.

Kohorn, B. D., Johansen, S., Shishido, A., Todorova, T., Martinez, R., Defeo, E. & Obregon, P. 2009. Pectin activation of MAP kinase and gene expression is WAK2 dependent. Plant J, 60, 974–82.

Kohorn, B. D., Kobayashi, M., Johansen, S., Friedman, H. P., Fischer, A. & Byers, N. 2006. Wall-associated kinase 1 (WAK1) is crosslinked in endomembranes, and transport to the cell surface requires correct cell-wall synthesis. Journal of cell science, 119, 2282–2290.

Kwezi, L., Ruzvidzo, O., Wheeler, J. I., Govender, K., Iacuone, S., Thompson, P. E., Gehring, C. & Irving, H. R. 2011. The phytosulfokine (PSK) receptor is capable of guanylate cyclase activity and enabling cyclic GMP-dependent signaling in plants. J Biol Chem, 286, 22580–8.

Li, H., Zhou, S. Y., Zhao, W. S., Su, S. C. & Peng, Y. L. 2009. A novel wall-associated receptor-like protein kinase gene, OsWAK1, plays important roles in rice blast disease resistance. Plant Mol Biol, 69.

Livak, K. J. & Schmittgen, T. D. 2001. Analysis of relative gene expression data using real-time quantitative PCR and the 2(-Delta Delta C(T)) Method. Methods, 25, 402–8.

Lozano-Durán, R. & Zipfel, C. 2015. Trade-off between growth and immunity: role of brassinosteroids. Trends in plant science, 20, 12–19.

Ma, Y., Walker, R. K., Zhao, Y. & Berkowitz, G. A. 2012. Linking ligand perception by PEPR pattern recognition receptors to cytosolic Ca2+ elevation and downstream immune signaling in plants. Proc Natl Acad Sci U S A, 109, 19852–7.

Macho, A. P. & Zipfel, C. 2014. Plant PRRs and the activation of innate immune signaling. Mol Cell, 54, 263–72.

Meier, S., Ruzvidzo, O., Morse, M., Donaldson, L., Kwezi, L. & Gehring, C. 2010. The Arabidopsis Wall Associated Kinase-Like 10 Gene Encodes a Functional Guanylyl Cyclase and Is Co-Expressed with Pathogen Defense Related Genes. PLoS ONE, 5, e8904.

Meng, X. & Zhang, S. 2013. MAPK cascades in plant disease resistance signaling. Annu Rev Phytopathol, 51, 245–66.

Millet, Y. A., Danna, C. H., Clay, N. K., Songnuan, W., Simon, M. D., Werck-Reichhart, D. & Ausubel, F. M. 2010. Innate immune responses activated in Arabidopsis roots by microbe-associated molecular patterns. Plant Cell, 22, 973–90.

Nan, W., Wang, X., Yang, L., Hu, Y., Wei, Y., Liang, X., Mao, L. & Bi, Y. 2014. Cyclic GMP is involved in auxin signalling during Arabidopsis root growth and development. J Exp Bot, 65, 1571–83.

Oñate-Sánchez, L. & Vicente-Carbajosa, J. 2008. DNA-free RNA isolation protocols for Arabidopsis thaliana, including seeds and siliques. BMC Research Notes, 1, 93.

Park, C. J., Peng, Y., Chen, X., Dardick, C., Ruan, D., Bart, R., Canlas, P. E. & Ronald, P. C. 2008. Rice XB15, a protein phosphatase 2c, negatively regulates cell death and XA21-mediated innate immunity. PLoS Biol, 6, e231.

Pillai, S. E., Kumar, C., Patel, H. K. & Sonti, R. V. 2018. Overexpression of a cell wall damage induced transcription factor, OsWRKY42, leads to enhanced callose deposition and tolerance to salt stress but does not enhance tolerance to bacterial infection. BMC Plant Biology, 18, 177.

Purkayastha, A., Mathur, S., Verma, V., Sharma, S. & Dasgupta, I. 2010. Virus-induced gene silencing in rice using a vector derived from a DNA virus. Planta, 232, 1531–1540.

Qi, Z., Verma, R., Gehring, C., Yamaguchi, Y., Zhao, Y., Ryan, C. A. & Berkowitz, G. A. 2010. Ca2+ signaling by plant Arabidopsis thaliana Pep peptides depends on AtPepR1, a receptor with guanylyl cyclase activity, and cGMP-activated Ca2+ channels. Proc Natl Acad Sci U S A, 107, 21193–8.

Rajeshwari, R., Jha, G. & Sonti, R. V. 2005. Role of an In Planta-Expressed Xylanase of Xanthomonas oryzae pv. oryzae in Promoting Virulence on Rice. Molecular Plant-Microbe Interactions, 18, 830–837.

Ranjan, A., Vadassery, J., Patel, H. K., Pandey, A., Palaparthi, R., Mithofer, A. & Sonti, R. V. 2015. Upregulation of jasmonate biosynthesis and jasmonate-responsive genes in rice leaves in response to a bacterial pathogen mimic. Funct Integr Genomics, 15, 363–73.

Rohila, J. S., Chen, M., Chen, S., Chen, J., Cerny, R., Dardick, C., Canlas, P., Xu, X., Gribskov, M., Kanrar, S., Zhu, J. K., Ronald, P. & Fromm, M. E. 2006. Protein-protein interactions of tandem affinity purification-tagged protein kinases in rice. Plant J, 46, 1–13.

Saintenac, C., Lee, W.-S., Cambon, F., Rudd, J. J., King, R. C., Marande, W., Powers, S. J., Berges, H., Phillips, A. L. & Uauy, C. 2018. Wheat receptor-kinase-like protein Stb6 controls gene-for-gene resistance to fungal pathogen Zymoseptoria tritici. Nature genetics, 50, 368.

Sun, W., Cao, Z., Li, Y., Zhao, Y. & Zhang, H. 2007. A simple and effective method for protein subcellular localization using Agrobacterium-mediated transformation of onion epidermal cells. Biologia, 62, 529–532.

Swiezawska, B., Jaworski, K., Duszyn, M., Pawelek, A. & Szmidt-Jaworska, A. 2017. The Hippeastrum hybridum PepR1 gene (HpPepR1) encodes a functional guanylyl cyclase and is involved in early response to fungal infection. J Plant Physiol, 216, 100–107.

Swiezawska, B., Jaworski, K., Szewczuk, P., Pawelek, A. & Szmidt-Jaworska, A. 2015. Identification of a Hippeastrum hybridum guanylyl cyclase responsive to wounding and pathogen infection. J Plant Physiol, 189, 77–86.

Verica, J. A. & He, Z. 2002. The Cell Wall-Associated Kinase (WAK) and WAK-Like Kinase Gene Family. Plant Physiology, 129, 455–459.

Wang, L., Einig, E., Almeida-Trapp, M., Albert, M., Fliegmann, J., Mithöfer, A., Kalbacher, H. & Felix, G. 2018. The systemin receptor SYR1 enhances resistance of tomato against herbivorous insects. Nature Plants, 4, 152–156.

Wong, A., Gehring, C. & Irving, H. R. 2015. Conserved Functional Motifs and Homology Modeling to Predict Hidden Moonlighting Functional Sites. Frontiers in Bioengineering and Biotechnology, 3, 82.

Xin, X. F. & He, S. Y. 2013. Pseudomonas syringae pv. tomato DC3000: a model pathogen for probing disease susceptibility and hormone signaling in plants. Annu Rev Phytopathol, 51, 473–98.

Xu, N., Fu, D., Li, S., Wang, Y. & Wong, A. 2018. GCPred: a web tool for guanylyl cyclase functional centre prediction from amino acid sequence. Bioinformatics, 34, 2134–2135.

Zhang, N., Zhang, B., Zuo, W., Xing, Y., Konlasuk, S., Tan, G., Zhang, Q., Ye, J. & Xu, M. 2017. Cytological and Molecular Characterization of ZmWAK-Mediated Head-Smut Resistance in Maize. Mol Plant Microbe Interact, 30, 455–465.

Zheng, L., Baumann, U. & Reymond, J.-L. 2004. An efficient one-step site-directed and site-saturation mutagenesis protocol. Nucleic Acids Research, 32, e115–e115.

Zuo, W., Chao, Q., Zhang, N., Ye, J., Tan, G., Li, B., Xing, Y., Zhang, B., Liu, H., Fengler, K. A., Zhao, J., Zhao, X., Chen, Y., Lai, J., Yan, J. & Xu, M. 2015. A maize wall-associated kinase confers quantitative resistance to head smut. Nat Genet, 47, 151–7.

