## Supplemental data for "The dual function receptor kinase, OsWAKL21.2, is involved in elaboration of lipaseA/esterase induced immune responses in rice"

| Supplemental Table 1: Differentially expressed genes after two hour of LipA treatment | | | | |
|---|---|---|---|---|
| **Affymetrix Probe ID** | **MSU ID[a]** | **RAP ID[b]** | **FC[c]** | **MSU name[d]** |
| Os.50533.1.S1_at | LOC_Os04g35680.1 | Os04g0437300 | 3.00 | U-box domain containing protein, expressed |
| OsAffx.31223.1.S1_at | LOC_Os11g30290.1 | Os11g0495400 | 2.83 | PHF5-like protein domain containing protein, expressed |
| Os.30000.1.S1_at | LOC_Os07g36560.1 | Os07g0550600 | 2.63 | transferase family protein, putative, expressed |
| Os.22312.3.A1_a_at | LOC_Os05g28740.1 | Os05g0355400 | 2.45 | universal stress protein domain containing protein, putative, expressed |
| Os.6244.1.S1_x_at | LOC_Os03g13740.1 | Os03g0240600 | 2.43 | immediate-early fungal elicitor protein CMPG1, putative, expressed |
| | LOC_Os01g57735.1 | Os01g0787300 | 2.43 | expressed protein |
| Os.54936.1.S1_at | LOC_Os03g57640.1 | Os03g0790500 | 2.36 | gibberellin receptor GID1L2, putative, expressed |
| Os.6288.1.S1_at | LOC_Os08g31850.1 | Os08g0412700 | 2.31 | expressed protein |
| Os.6244.1.S1_at | LOC_Os03g13740.1 | Os03g0240600 | 2.29 | immediate-early fungal elicitor protein CMPG1, putative, expressed |
| Os.27507.1.S1_at | LOC_Os06g35700.1 | Os06g0549900 | 2.18 | reticuline oxidase-like protein precursor, putative, expressed |
| Os.29859.1.S1_at | LOC_Os07g14080.1 | Os07g0244200 | 2.14 | transferase family protein, putative, expressed |
| Os.38447.1.S1_s_at | LOC_Os06g07030.1 | Os06g0166400 | 2.10 | AP2 domain containing protein, expressed |
| Os.26626.1.S1_at | LOC_Os05g33400.1 | Os05g0402900 | 2.01 | basic 7S globulin precursor, putative, expressed |
| Os.1043.1.S1_at | LOC_Os01g42860.1 | Os01g0615100 | 1.99 | inhibitor I family protein, putative, expressed |
| Os.36283.1.S1_at | LOC_Os12g25090.1 | Os12g0437800 | 1.97 | expressed protein |
| | LOC_Os12g25090.2 | | 1.97 | expressed protein |
| Os.10579.1.S1_at | LOC_Os08g02700.1 | Os08g0120600 | 1.97 | fructose-bisphospate aldolase isozyme, putative, expressed |
| Os.4804.1.S1_at | LOC_Os06g07030.1 | Os06g0166400 | 1.94 | AP2 domain containing protein, expressed |
| Os.53936.1.S1_at | LOC_Os03g15270.1 | Os03g0258200 | 1.91 | gibberellin receptor GID1L2, putative, expressed |
| Os.19861.1.S1_at | LOC_Os07g03730.1 | Os07g0129300 | 1.89 | SCP-like extracellular protein, expressed |
| Os.53744.2.S1_x_at | LOC_Os12g40419.1 | Os12g0595800 | 1.88 | WAKL21 wall associated kinase like receptor kinase |
| | LOC_Os12g40419.2 | | 1.88 | |
| | LOC_Os12g40419.3 | | 1.88 | |

| | | | | |
|---|---|---|---|---|
| Os.9206.1.S1_at | LOC_Os10g35630.1 | Os10g0499400 | 1.85 | cystathionin beta synthase protein, putative, expressed |
| | LOC_Os10g35630.2 | | 1.85 | |
| Os.38299.1.S1_at | LOC_Os07g02330.1 | Os07g0114000 | 1.85 | protein phosphatase 2C, putative, expressed |
| Os.27043.1.A1_at | LOC_Os05g03920.1 | Os05g0130100 | 1.81 | TKL_IRAK_DUF26-lf.3 - DUF26 kinases have homology to DUF26 containing loci, expressed |
| Os.34962.1.S1_at | LOC_Os01g38110.1 | Os01g0561600 | 1.80 | cytochrome P450, putative, expressed |
| OsAffx.30533.1.S1_s_at | LOC_Os10g25830.1 | Os10g0397800 | 1.78 | mitochondrial carrier protein, putative, expressed |
| Os.40428.1.S1_at | LOC_Os01g73770.1 | Os01g0968800 | 1.78 | dehydration-responsive element-binding protein, putative, expressed |
| Os.1665.2.S1_x_at | LOC_Os01g07300.1 | Os01g0167400 | 1.77 | uncharacterized 50.6 kDa protein in the 5region of gyrA and gyrB, putative, expressed |
| | LOC_Os01g07300.2 | | 1.77 | |
| Os.24003.2.S1_x_at | LOC_Os11g40570.1 | Os11g0621000 | 1.76 | plant viral response family protein, putative, expressed |
| | LOC_Os11g40570.2 | | 1.76 | |
| | LOC_Os11g40570.3 | | 1.76 | |
| Os.1665.1.S1_a_at | LOC_Os01g07300.1 | Os01g0167400 | 1.75 | uncharacterized 50.6 kDa protein in the 5region of gyrA and gyrB, putative, expressed |
| | LOC_Os01g07300.2 | | 1.75 | |
| Os.6321.1.S1_at | LOC_Os01g63690.1 | Os01g0855600 | 1.74 | hs1, putative, expressed |
| Os.50013.1.S1_at | LOC_Os10g28210.1 | Os10g0417800 | 1.73 | plant-specific domain TIGR01615 family protein, expressed |
| Os.31344.1.S1_at | LOC_Os01g63970.1 | Os01g0858900 | 1.73 | sialyltransferase family domain containing protein, expressed |
| Os.54944.1.S1_at | LOC_Os02g52670.1 | Os02g0764700 | 1.73 | AP2 domain containing protein, expressed |
| | LOC_Os01g54340.1 | Os01g0747300 | 1.73 | plant-specific domain TIGR01615 family protein, expressed |
| OsAffx.19579.1.S1_at | LOC_Os12g09640.1 | Os12g0198200 | 1.72 | protein phosphatase 2C, putative, expressed |
| OsAffx.26677.1.S1_x_at | LOC_Os05g01444.1 | Os05g0104700 | 1.70 | polygalacturonase inhibitor 2 precursor, putative, expressed |
| | LOC_Os05g01430.1 | Os05g0104600 | 1.70 | polygalacturonase inhibitor 2 precursor, putative, expressed |
| Os.55944.1.S1_at | LOC_Os11g06150.1 | Os11g0160400 | 1.70 | basic proline-rich protein precursor, putative, expressed |

| | | | | |
|---|---|---|---|---|
| Os.57041.1.S1_at | LOC_Os12g18560.1 | Os12g0283400 | 1.69 | invertase/pectin methylesterase inhibitor family protein, putative, expressed |
| OsAffx.23706.1.S1_at | LOC_Os01g48120.1 | | 1.69 | expressed protein |
| OsAffx.26845.1.S1_at | LOC_Os05g12090.1 | Os05g0211700 | 1.69 | VQ domain containing protein, putative |
| Os.17036.1.S1_x_at | LOC_Os05g03620.1 | Os05g0127300 | 1.68 | TKL_IRAK_CR4L.4 - The CR4L subfamily has homology with Crinkly4, expressed |
| Os.18229.1.S1_at | LOC_Os01g72610.1 | Os01g0956200 | 1.66 | glycosyltransferase, putative, expressed |
| Os.52414.1.S1_at | LOC_Os02g19650.1 | Os02g0299300 | 1.66 | hydrolase, alpha/beta fold family domain containing protein, expressed |
| | LOC_Os02g19650.3 | | 1.66 | |
| Os.27395.1.S1_a_at | LOC_Os04g57760.1 | Os04g0674000 | 1.65 | expressed protein |
| OsAffx.14458.1.S1_x_at | LOC_Os04g57760.1 | Os04g0674000 | 1.64 | expressed protein |
| Os.52004.1.S1_at | LOC_Os09g31940.1 | | 1.64 | retrotransposon protein, putative, unclassified |
| Os.30376.1.S1_at | LOC_Os01g02130.1 | Os01g0111700 | 1.62 | expressed protein |
| Os.48131.1.S1_s_at | LOC_Os04g58890.1 | Os04g0685700 | 1.61 | expressed protein |
| Os.18717.2.S1_at | LOC_Os09g28180.1 | Os09g0454900 | 1.60 | D-mannose binding lectin family protein, expressed |
| Os.12030.1.S1_at | LOC_Os01g67480.1 | Os01g0900800 | 1.60 | helix-loop-helix DNA-binding domain containing protein, expressed |
| Os.17181.1.S1_at | LOC_Os07g32010.1 | Os07g0503300 | 1.60 | UDP-glucoronosyl and UDP-glucosyl transferase domain containing protein, expressed |
| Os.3581.1.A1_at | LOC_Os06g05070.1 | Os06g0142650 | 1.60 | protein kinase domain containing protein, expressed |
| Os.10872.1.S1_at | LOC_Os07g09420.1 | Os07g0192000 | 1.59 | ATPase, putative, expressed |
| Os.46956.1.S1_at | LOC_Os01g50940.1 | Os01g0705700 | 1.58 | helix-loop-helix DNA-binding domain containing protein, expressed |
| Os.37865.1.S1_at | LOC_Os02g40700.1 | Os02g0620400 | 1.58 | enzyme of the cupin superfamily protein, putative, expressed |
| Os.27138.1.S1_at | LOC_Os10g21590.1 | Os10g0360100 | 1.57 | transporter family protein, putative, expressed |
| | LOC_Os10g21590.2 | | 1.57 | |

| | | | | |
|---|---|---|---|---|
| Os.10709.1.S1_at | LOC_Os03g22720.1 | Os03g0349600 | 1.57 | expressed protein |
| | LOC_Os03g22720.2 | | 1.57 | |
| | LOC_Os03g22720.3 | | 1.57 | |
| Os.50310.1.S1_at | LOC_Os01g66544.1 | Os01g0888900 | 1.57 | expressed protein |
| OsAffx.7301.1.S1_at | LOC_Os11g35410.1 | Os11g0558400 | 1.56 | expressed protein |
| Os.12921.1.S1_at | LOC_Os04g57770.1 | Os04g0674050 | 1.55 | expressed protein |
| Os.49855.1.S1_at | LOC_Os02g57560.1 | Os02g0821400 | 1.54 | tyrosine protein kinase domain containing protein, putative, expressed |
| Os.55582.1.S1_at | LOC_Os09g28650.1 | Os09g0460700 | 1.54 | gibberellin receptor, putative, expressed |
| Os.32022.1.S1_x_at | LOC_Os07g37620.1 | Os07g0563400 | 1.52 | fiber expressed protein, putative, expressed |
| OsAffx.4103.1.S1_s_at | LOC_Os04g45940.1 | Os04g0543500 | 1.52 | transcription factor like protein, putative, expressed |
| Os.14823.1.S1_s_at | LOC_Os03g20090.1 | Os03g0315400 | 1.52 | MYB family transcription factor, putative, expressed |
| Os.55671.1.S1_at | LOC_Os06g44250.1 | Os06g0652200 | 1.51 | haemolysin-III, putative, expressed |
| Os.6671.2.S1_x_at | LOC_Os05g39930.1 | Os05g0476700 | 1.51 | spotted leaf 11, putative, expressed |
| Os.19326.1.S1_at | LOC_Os07g34940.1 | Os07g0533800 | 1.51 | aspartic proteinase nepenthesin-1 precursor, putative, expressed |
| OsAffx.17942.1.S1_at | LOC_Os09g28160.1 | Os09g0454600 | 1.51 | phosphate carrier protein, mitochondrial precursor, putative, expressed |
| Os.49607.1.S1_at | LOC_Os03g09880.1 | Os03g0194600 | 1.50 | AIR12, putative, expressed |
| Os.50292.1.S1_at | LOC_Os02g13560.1 | Os02g0229400 | -1.52 | transporter family protein, putative, expressed |
| | LOC_Os02g13560.2 | | -1.52 | |
| | LOC_Os02g13560.3 | | -1.52 | |
| | LOC_Os02g13560.4 | | -1.52 | |
| Os.46223.1.S1_s_at | LOC_Os09g33690.1 | Os09g0511700 | -1.55 | Os9bglu32 - beta-glucosidase homologue, similar to G. max hydroxyisourate hydrolase, expressed |
| | LOC_Os09g33690.2 | | -1.55 | |
| | LOC_Os09g33690.3 | | -1.55 | |
| | LOC_Os09g33690.4 | | -1.55 | |
| Os.27705.1.S1_a_at | N.A | N.A | -1.60 | N.A |

| | | | | |
|---|---|---|---|---|
| Os.49564.1.S1_at | LOC_Os04g33610.1 | Os04g0412100 | -1.64 | expressed protein |
| Os.5318.1.S1_a_at | LOC_Os10g42610.1 | Os10g0576600 | -1.68 | expressed protein |
| | LOC_Os10g42610.2 | | -1.68 | |
| Os.7370.1.S1_at | LOC_Os03g40540.1 | Os03g0602300 | -1.68 | cytochrome P450, putative, expressed |
| Os.338.1.S1_x_at | LOC_Os03g17980.2 | Os03g0289100 | -1.68 | CAMK_KIN1/SNF1/Nim1_like_AMPKh.2 - CAMK includes calcium/calmodulin depedent protein kinases, expressed |
| | LOC_Os03g17980.1 | | -1.68 | |
| | LOC_Os08g37800.1 | Os08g0484600 | -1.68 | CAMK_KIN1/SNF1/Nim1_like_AMPKh.4 - CAMK includes calcium/calmodulin depedent protein kinases, expressed |
| Os.27967.1.A1_at | LOC_Os01g63620.1 | Os01g0855200 | -1.69 | expressed protein |
| Os.54940.1.S1_at | LOC_Os04g44150.1 | Os04g0522500 | -1.71 | gibberellin 2-beta-dioxygenase 7, putative, expressed |
| Os.3386.1.S1_x_at | LOC_Os06g10350.1 | Os06g0205100 | -2.08 | MYB family transcription factor, putative, expressed |

[a]: Gene ID on Michigan State University database

[b]: Gene ID on Rice Annotation Project Database

[c]: FC: Fold change

[d]: Name assigned in MSU database

N.A.: Not annotated

| Supplemental Table 2: List of differentially expressed genes after 2hr and 12hr of LipA treatment | | | | | |
|---|---|---|---|---|---|
| **Common probes IDs** | **MSU ID[a]** | **RAP ID[b]** | **FC-2hr** | **FC-12hr** | **MSU name[c]** |
| Os.50533.1.S1_at | LOC_Os04g35680.1 | Os04g0437300 | 3 | 6.75 | U-box domain containing protein, expressed |
| OsAffx.31223.1.S1_at | LOC_Os11g30290.1 | Os11g0495400 | 2.83 | 2.35 | PHF5-like protein domain containing protein, expressed |
| Os.30000.1.S1_at | LOC_Os07g36560.1 | Os07g0550600 | 2.63 | 11.25 | transferase family protein, putative, expressed |
| Os.6244.1.S1_x_at | LOC_Os03g13740.1 | Os03g0240600 | 2.43 | 5.77 | immediate-early fungal elicitor protein CMPG1, putative, expressed |
| Os.54936.1.S1_at | LOC_Os03g57640.1 | Os03g0790500 | 2.36 | 3.16 | gibberellin receptor GID1L2, putative, expressed |
| Os.6288.1.S1_at | LOC_Os08g31850.1 | Os08g0412700 | 2.31 | 2.69 | expressed protein |
| Os.6244.1.S1_at | LOC_Os03g13740.1 | Os03g0240600 | 2.29 | 5.94 | immediate-early fungal elicitor protein CMPG1, putative, expressed |
| Os.27507.1.S1_at | LOC_Os06g35700.1 | Os06g0549900 | 2.18 | 4.00 | reticuline oxidase-like protein precursor, putative, expressed |
| Os.29859.1.S1_at | LOC_Os07g14080.1 | Os07g0244200 | 2.14 | 1.72 | transferase family protein, putative, expressed |
| Os.26626.1.S1_at | LOC_Os05g33400.1 | Os05g0402900 | 2.01 | 2.60 | basic 7S globulin precursor, putative, expressed |
| Os.1043.1.S1_at | LOC_Os01g42860.1 | Os01g0615100 | 1.99 | 2.61 | inhibitor I family protein, putative, expressed |
| Os.10579.1.S1_at | LOC_Os08g02700.1 | Os08g0120600 | 1.97 | 1.92 | fructose-bisphospate aldolase isozyme, putative, expressed |
| Os.53744.2.S1_x_at | LOC_Os12g40419 | Os12g0595800 | 1.88 | 2.43 | WAKL21 wall associated kinase like receptor kinase |
| Os.9206.1.S1_at | LOC_Os10g35630 | Os10g0499400 | 1.85 | 1.77 | cystathionin beta synthase protein, putative, expressed |
| Os.38299.1.S1_at | LOC_Os07g02330.1 | Os07g0114000 | 1.85 | 2.24 | protein phosphatase 2C, putative, expressed |
| Os.27043.1.A1_at | LOC_Os05g03920.1 | Os05g0130100 | 1.81 | 1.78 | TKL_IRAK_DUF26-lf.3 - DUF26 kinases have homology to DUF26 containing loci, expressed |
| Os.34962.1.S1_at | LOC_Os01g38110.1 | Os01g0561600 | 1.8 | 3.13 | cytochrome P450, putative, expressed |
| Os.24003.2.S1_x_at | LOC_Os11g40570 | Os11g0621000 | 1.76 | 1.70 | plant viral response family protein, putative, expressed |
| Os.6321.1.S1_at | LOC_Os01g63690.1 | Os01g0855600 | 1.74 | 2.85 | hs1, putative, expressed |

| | | | | | |
|---|---|---|---|---|---|
| Os.31344.1.S1_at | LOC_Os01g63970.1 | Os01g0858900 | 1.73 | 1.85 | sialyltransferase family domain containing protein, expressed |
| OsAffx.26677.1.S1_x_at | LOC_Os05g01444.1 | Os05g0104700 | 1.7 | 1.59 | polygalacturonase inhibitor 2 precursor, putative, expressed |
| Os.55944.1.S1_at | LOC_Os11g06150.1 | Os11g0160400 | 1.7 | 1.84 | basic proline-rich protein precursor, putative, expressed |
| Os.57041.1.S1_at | LOC_Os12g18560.1 | Os12g0283400 | 1.69 | 2.23 | invertase/pectin methylesterase inhibitor family protein, putative, expressed |
| OsAffx.26845.1.S1_at | LOC_Os05g12090.1 | Os05g0211700 | 1.69 | 1.89 | VQ domain containing protein, putative |
| Os.17036.1.S1_x_at | LOC_Os05g03620.1 | Os05g0127300 | 1.68 | 2.56 | TKL_IRAK_CR4L.4 - The CR4L subfamily has homology with Crinkly4, expressed |
| Os.30376.1.S1_at | LOC_Os01g02130.1 | Os01g0111700 | 1.62 | 4.22 | expressed protein |
| Os.17181.1.S1_at | LOC_Os07g32010.1 | Os07g0503300 | 1.6 | 2.52 | UDP-glucoronosyl and UDP-glucosyl transferase domain containing protein, expressed |
| Os.10872.1.S1_at | LOC_Os07g09420.1 | Os07g0192000 | 1.59 | 2.48 | ATPase, putative, expressed |
| Os.46956.1.S1_at | LOC_Os01g50940.1 | Os01g0705700 | 1.58 | 1.83 | helix-loop-helix DNA-binding domain containing protein, expressed |
| Os.37865.1.S1_at | LOC_Os02g40700.1 | Os02g0620400 | 1.58 | 2.21 | enzyme of the cupin superfamily protein, putative, expressed |
| Os.27138.1.S1_at | LOC_Os10g21590.1 | Os10g0360100 | 1.57 | 2.20 | transporter family protein, putative, expressed |
| Os.12921.1.S1_at | LOC_Os04g57770.1 | Os04g0674050 | 1.55 | 1.60 | expressed protein |
| Os.14823.1.S1_s_at | LOC_Os03g20090.1 | Os03g0315400 | 1.52 | 1.96 | MYB family transcription factor, putative, expressed |
| Os.55671.1.S1_at | LOC_Os06g44250.1 | Os06g0652200 | 1.51 | 3.16 | haemolysin-III, putative, expressed |
| Os.6671.2.S1_x_at | LOC_Os05g39930.1 | Os05g0476700 | 1.51 | 2.54 | spotted leaf 11, putative, expressed |
| Os.19326.1.S1_at | LOC_Os07g34940.1 | Os07g0533800 | 1.51 | 1.87 | aspartic proteinase nepenthesin-1 precursor, putative, expressed |
| OsAffx.17942.1.S1_at | LOC_Os09g28160.1 | Os09g0454600 | 1.51 | 3.40 | phosphate carrier protein, mitochondrial precursor, putative, expressed |
| Os.27705.1.S1_a_at | N.A | N.A | -1.6 | -1.68 | N.A |

[a]: Gene ID in Michigan State University database

[b]: Gene ID in Rice Annotation Project database

[c]: Name assigned in MSU database

FC: Fold change

N.A.: Not annotated

### Supplemental Table 3: Frequency of differentially expressed genes after LipA treatment in the microarray data performed after 24hr of *Xanthomonas oryzae* treatment in GEO submission GSE36272

| Affy. Probe ID | MSU ID[a] | RAP ID[b] | MSU annotation[c] | Data points[d] | Ave FC | SD | Min FC | Max FC | LipA 2hr FC | LipA 12hr FC |
|---|---|---|---|---|---|---|---|---|---|---|
| Os.50533.1.S1_at | LOC_Os04g35680.1 | Os04g0437300 | U-box domain containing protein, expressed | 16 | 3.52 | 1.73 | 1.65 | 8.91 | 3.00 | 6.75 |
| Os.10579.1.S1_at | LOC_Os08g02700.1 | Os08g0120600 | fructose-bisphospate aldolase isozyme, putative, expressed | 16 | 4.62 | 2.30 | 1.63 | 9.66 | 1.97 | 1.92 |
| Os.53744.2.S1_x_at | LOC_Os12g40419.1 | Os12g0595800 | WAKL21 wall associated kinase like receptor kinase | 16 | 3.91 | 2.08 | 1.56 | 8.77 | 1.88 | 2.43 |
| Os.30000.1.S1_at | LOC_Os07g36560.1 | Os07g0550600 | transferase family protein, putative, expressed | 15 | 53.28 | 71.96 | 1.94 | 248.34 | 2.63 | 11.25 |
| Os.17181.1.S1_at | LOC_Os07g32010.1 | Os07g0503300 | UDP-glucoronosyl and UDP-glucosyl transferase domain containing protein, expressed | 15 | 2.08 | 0.43 | 1.57 | 2.92 | 1.60 | 2.52 |
| Os.55671.1.S1_at | LOC_Os06g44250.1 | Os06g0652200 | haemolysin-III, putative, expressed | 15 | 10.37 | 11.76 | 2.12 | 38.59 | 1.51 | 3.16 |
| Os.46956.1.S1_at | LOC_Os01g50940.1 | Os01g0705700 | helix-loop-helix DNA-binding domain containing protein, expressed | 14 | 6.70 | 4.50 | 2.48 | 18.17 | 1.58 | 1.83 |
| OsAffx.17942.1.S1_at | LOC_Os09g28160.1 | Os09g0454600 | phosphate carrier protein, mitochondrial precursor, putative, expressed | 14 | 4.54 | 2.04 | 1.82 | 8.13 | 1.51 | 3.40 |
| Os.27043.1.A1_at | LOC_Os05g03920.1 | Os05g0130100 | TKL_IRAK_DUF26-lf.3 - DUF26 kinases, expressed | 13 | 2.99 | 0.97 | 1.56 | 4.46 | 1.81 | 1.78 |
| Os.24003.2.S1_x_at | LOC_Os11g40570.1 | Os11g0621000 | plant viral response family protein, putative, expressed | 12 | 2.31 | 0.68 | 1.51 | 3.44 | 1.76 | 1.70 |
| Os.6321.1.S1_at | LOC_Os01g63690.1 | Os01g0855600 | hs1, putative, expressed | 12 | 1.51 | 3.14 | -3.89 | 5.68 | 1.74 | 2.85 |

| | | | | | | | | | | |
|---|---|---|---|---|---|---|---|---|---|---|
| Os.17036.1.S1_x_at | LOC_Os05g03620.1 | Os05g0127300 | TKL_IRAK_CR4L.4 - The CR4L subfamily has homology with Crinkly4, expressed | 12 | 2.70 | 0.49 | 1.91 | 3.32 | 1.68 | 2.56 |
| Os.10872.1.S1_at | LOC_Os07g09420.1 | Os07g0192000 | ATPase, putative, expressed | 12 | 2.74 | 0.58 | 1.66 | 3.73 | 1.59 | 2.48 |
| Os.6244.1.S1_x_at | LOC_Os03g13740.1 | Os03g0240600 | immediate-early fungal elicitor protein CMPG1, putative, expressed | 11 | 7.34 | 5.49 | 2.02 | 18.92 | 2.43 | 5.77 |
| Os.37865.1.S1_at | LOC_Os02g40700.1 | Os02g0620400 | enzyme of the cupin superfamily protein, putative, expressed | 10 | 2.59 | 0.52 | 1.81 | 3.21 | 1.58 | 2.21 |
| Os.6244.1.S1_at | LOC_Os03g13740.1 | Os03g0240600 | immediate-early fungal elicitor protein CMPG1, putative, expressed | 9 | 9.86 | 6.66 | 3.43 | 21.33 | 2.29 | 5.94 |
| Os.31344.1.S1_at | LOC_Os01g63970.1 | Os01g0858900 | sialyltransferase family domain containing protein, expressed | 9 | 1.78 | 0.20 | 1.56 | 2.19 | 1.73 | 1.85 |
| Os.55944.1.S1_at | LOC_Os11g06150.1 | Os11g0160400 | basic proline-rich protein precursor, putative, expressed | 9 | 3.17 | 2.28 | 1.51 | 8.92 | 1.70 | 1.84 |
| Os.19326.1.S1_at | LOC_Os07g34940.1 | Os07g0533800 | aspartic proteinase nepenthesin-1 precursor, putative, expressed | 9 | 3.26 | 1.07 | 1.91 | 5.55 | 1.51 | 1.87 |
| Os.30376.1.S1_at | LOC_Os01g02130.1 | Os01g0111700 | expressed protein | 8 | 2.65 | 2.95 | -4.45 | 5.64 | 1.62 | 4.22 |
| Os.57041.1.S1_at | LOC_Os12g18560.1 | Os12g0283400 | invertase/pectin methylesterase inhibitor family protein, putative, expressed | 6 | -13.94 | 7.99 | -31.27 | -6.42 | 1.69 | 2.23 |
| Os.6671.2.S1_x_at | LOC_Os05g39930.1 | Os05g0476700 | spotted leaf 11, putative, expressed | 6 | 2.38 | 0.74 | 1.7 | 3.79 | 1.51 | 2.54 |
| Os.54936.1.S1_at | LOC_Os03g57640.1 | Os03g0790500 | gibberellin receptor GID1L2, putative, expressed | 5 | 5.42 | 0.32 | 5.07 | 5.86 | 2.36 | 3.16 |
| Os.29859.1.S1_at | LOC_Os07g14080.1 | Os07g0244200 | transferase family protein, putative, expressed | 5 | 2.15 | 0.37 | 1.74 | 2.77 | 2.14 | 1.72 |
| Os.27138.1.S1_at | LOC_Os10g21590.1 | Os10g0360100 | transporter family protein, putative, expressed | 4 | -0.58 | 2.13 | -1.89 | 3.11 | 1.57 | 2.20 |
| Os.27705.1.S1_a_at | N.A. | N.A. | N.A. | 4 | 1.97 | 0.30 | 1.52 | 2.35 | -1.60 | -1.68 |
| OsAffx.31223.1.S1_at | LOC_Os11g30290.1 | Os11g0495400 | PHF5-like protein domain containing protein, expressed | 3 | 4.65 | 1.45 | 2.81 | 6.34 | 2.83 | 2.35 |
| Os.26626.1.S1_at | LOC_Os05g33400.1 | Os05g0402900 | basic 7S globulin precursor, putative, expressed | 3 | 3.67 | 0.62 | 2.82 | 4.28 | 2.01 | 2.60 |
| Os.9206.1.S1_at | LOC_Os10g35630.1 | Os10g0499400 | cystathionin beta synthase protein, putative, expressed | 3 | -2.36 | 0.18 | -2.52 | -2.11 | 1.85 | 1.77 |

| | | | | | | | | | | |
|---|---|---|---|---|---|---|---|---|---|---|
| OsAffx.26845.1.S1_at | LOC_Os05g12090.1 | Os05g0211700 | VQ domain containing protein, putative | 3 | 2.04 | 0.19 | 1.77 | 2.19 | 1.69 | 1.89 |
| Os.14823.1.S1_s_at | LOC_Os03g20090.1 | Os03g0315400 | MYB family transcription factor, putative, expressed | 2 | -0.21 | 1.78 | -1.99 | 1.57 | 1.52 | 1.96 |
| Os.38299.1.S1_at | LOC_Os07g02330.1 | Os07g0114000 | protein phosphatase 2C, putative, expressed | 1 | 1.52 | 0 | 1.52 | 1.52 | 1.85 | 2.24 |
| Os.34962.1.S1_at | LOC_Os01g38110.1 | Os01g0561600 | cytochrome P450, putative, expressed | 1 | 3.03 | 0 | 3.03 | 3.03 | 1.80 | 3.13 |
| Os.6288.1.S1_at | LOC_Os08g31850.1 | Os08g0412700 | expressed protein | 0 | | | | | 2.31 | 2.69 |
| Os.27507.1.S1_at | LOC_Os06g35700.1 | Os06g0549900 | reticuline oxidase-like protein precursor, putative, expressed | 0 | | | | | 2.18 | 4.00 |
| Os.1043.1.S1_at | LOC_Os01g42860.1 | Os01g0615100 | inhibitor I family protein, putative, expressed | 0 | | | | | 1.99 | 2.61 |
| OsAffx.26677.1.S1_x_at | LOC_Os05g01444.1 | Os05g0104700 | polygalacturonase inhibitor 2 precursor, putative, expressed | 0 | | | | | 1.70 | 1.59 |
| Os.12921.1.S1_at | LOC_Os04g57770.1 | Os04g0674050 | expressed protein | 0 | | | | | 1.55 | 1.60 |

[a]: Gene ID in Michigan State University database

[b]: Gene ID in Rice Annotation Project database

[c]: Name assigned in MSU database

[d]: Number of times gene was differentially expressed out of 18 data points

Ave FC: Average fold change
SD: Standard deviation
N.A.: Not annotated
FC: Fold change
Min FC: Minimun fold change
Max FC: Maximum fold change

| Supplemental Table 4: Primers used in this study | |
|---|---|
| **Primers used for cloning** | |
| **Primer** | **Sequence (5'-3')** |
| OsWAKL21.2 F | CACCATGCACCTCGCCGGCGGCC |
| OsWAKL21 R | CTAAGCAAATCGCGGCATGGAGCC |
| OsWAKL21-Ter R | AGCAAATCGCGGCATGGAGCC |
| $OsWAKL21_{376}$ -F | CACCTCCTTCTCCCACACGCACCG |
| M13 F | GTAAAACGACGGCCAGT |
| M13 R | GGAAACAGCTATGACCATG |
| pMDC7 F | CAGCAGTCGAGGTAAGAT |
| pMDC7 R | GGTGTGTGGGCAATGAAA |
| pH7FWG2 F | CCGCACTAGTGATATCACAAGTTT |
| pH7FWG2 R | TTACTTGTACAGCTCGTCCATG |
| T7 F | TAATACGACTCACTATAGGG |
| T7 R | GCTAGTTATTGCTCAGCGG |
| PACI+WAK 1-20 F | TTAATTAAATGCACCTCGCCGGCGGCCG |
| PACI+WAK 451-470 F | TTAATTAATGCAGCGTCCCCGCCGAAGC |
| MLUI +WAK 300-282 R | ACGCGTTTGAACGACTTGCCGACGA |
| MLUI+WAK 600-582 R | ACGCGTAACAGGCCCACACCCTTCG |
| pRTBV F | GGATCCGGGCCCTTAATTA |
| pRTBV R | AGGCTGGAGGCATAAACGC |
| **Primers used for site directed mutagenesis** | |
| **Primer Name** | **Sequence (5'-3')** |
| OsWAKL21.2 D504A F | GCCCATCCTCCACCGCGCGGTCAAGTCCAGCAACAT |
| OsWAKL21.2 D504A R | ATGTTGCTGGACTTGACCGCGCGGTGGAGGATGGGC |
| OsWAKL21.2 K407A F | CGCTGGTGGCGATCGCGCGGATGCGGCGGC |
| OsWAKL21.2 K407A R | GCCGCCGCATCCGCGCGATCGCCACCAGCG |
| OsWAKL21.2 T542AT547A F | TGTCGCACGTCTCGGCGGCGCCGCAGGGCGC GCCGGGGTACCTC |
| OsWAKL21.2 T542AT547A R | GAGGTACCCCGGCGCGCCCTGCGGCGCCGC CGAGACGTGCGACA |
| WAKL21.2-S569A, G571A F | AAGAGCGACGTCTACGCTTTCGCTGTCGTCCTCCTCG |
| WAKL21.2-S569A, G571A R | CGAGGAGGACGACAGCGAAAGCGTAGACGTCGCTCTT |
| WAKL21.2 K582Q F | TCACCGCCATGCAAGTCGTCGACTT |
| WAKL21.2 K582Q R | AAGTCGACGACTTGCATGGCGGTGA |
| **Primers used for qRT-PCR in rice** | |
| **Primer name** | **Sequence (5'-3')** |
| OsActin1 F | TGGATTGGAGGATCCATCTTGGC |
| OsActin1 R | CCTTGGCAATCCACATCTGCTG |
| OsGAPDH F | ACGGGAATGTCCTTCCGTGTTC |
| OsGAPDH R | AGCTTTCCCTCTGATGCAGACTTG |
| OsWAKL21.1 F | CCTTGGCAATCCACATCTGCTG |

| | |
|---|---|
| OsWAKL21.1 R | AACTCGGCAAGCGAGCCTAATG |
| OsWAKL21.2 F | GCCACTTTCCCGCTAAGAAGAG |
| OsWAKL21.2 R | CGCCAAGACACCTCCAACTATG |
| OsWAKL21.3 F | GGATGCAACCTTTCCGCTAAGAAG |
| OsWAKL21.3 R | CGCCAAGACACCTCCAACTATG |
| OsPR1a F | AGCTGTACTGTCAGCCGTATTTGC |
| OsPR1a R | ACCATGCATGTAACCACGAAGGAC |
| OsPR10a/PBZ1 F | ACGCCGCAAGTCATGTCCTAAAG |
| OsPR10a/PBZ1 R | TCGAGTGTGACTTGAGCTTCCC |
| OsPR10/PBZ14 F | ATGAAGCTCAACCCTGCTGTGG |
| OsPR10/PBZ14 R | TAATGTGAGCTGCGTTGTCACG |
| OsSERK2 F | ACTCTGGTCAATCCGTGCACTTG |
| OsSERK2 R | AGTGCAGCATTCCCAAGATCAAC |
| OsPAL3 F | TCATGTCCTCCACGTTCTTGGTC |
| OsPAL3 R | GCTCTTGACGTTCTCCTCGATTTG |
| LOC_Os01g50940 F | TGCGTACGGTATAGCTGCCAAC |
| LOC_Os01g50940 R | TGCTTGAAGTCACAAGGAGTTGC |
| LOC_Os02g43790 F | TGGTGAGCTAAGTGGCGATGTG |
| LOC_Os02g43790 R | AGCAGCAATCGATCACGCACAG |
| LOC_Os03g08310 F | CGGTCGAGTTGGAAGATGGTTC |
| LOC_Os03g08310 R | TCAGGCTCGGCGAAATCAACTC |
| LOC_Os03g08330 F | AACTCACCAAGCAAAGCACCAG |
| LOC_Os03g08330 R | AACGCGGCTTCTCTTCACCTTC |
| LOC_Os03g13740 F | GCTCAACAAGCACAAAGGGTTGG |
| LOC_Os03g13740 R | TTGAGCCCTCTGAAATCCACAGC |
| LOC_Os03g55800 F | TCGACGAATTGACACCATCTGCAC |
| LOC_Os03g55800 R | AGCTCCAACTGCAAGTCCACAG |
| LOC_Os04g23550 F | TGGCGCGAACAAGAACATCCTC |
| LOC_Os04g23550 R | GACGCCTTGTCCATCTTGGTGATG |
| LOC_Os04g35680 F | TGAGGAGCTCTTGATTCGATTCGG |
| LOC_Os04g35680 R | AGGTGCGGAATGCTCATCTCTTC |
| LOC_Os05g03620 F | ACAACAGCTCGTGCAAATGCG |
| LOC_Os05g03620 R | TCACAGAACCTTCTGCAGATGACG |
| LOC_Os06g44250 F | TGTGTATGTATGTGCGTGCCATTG |
| LOC_Os06g44250 R | CCAAAGAATCACCTGTGCTACGTC |
| LOC_Os06g51050 F | AGCTATGGCGATAACCTGGATTGC |
| LOC_Os06g51050 R | TATCAACTAGGAAGGCGGGTAGGG |
| LOC_Os07g09420 F | TTGGTCAAGGAGCTCGAGAAGG |
| LOC_Os07g09420 R | AGTCTACTCCTCGTCGTCATCG |
| LOC_Os07g32010 F | TCCCGAAGAACTGAGCATACGTG |
| LOC_Os07g32010 R | ACAAGCGGCATCATCTCTTGTTAC |
| LOC_Os07g36560 F | GCTCCTCTTCATTCAGGTGACG |

| | |
|---|---|
| LOC_Os07g36560 R | GTCGGCGATGTTGTGGCATATC |
| LOC_Os08g02700 F | TTCCCGCCATCTCTGCATCTTC |
| LOC_Os08g02700 R | GCGTTCCTGATCAACTCATCCTTG |
| LOC_Os08g36920 F | ACTCACATGACCAACCGGATCTC |
| LOC_Os08g36920 R | GCCGTCGAATCGGATCATGTACTC |
| LOC_Os08g39840 F | GATCGACATCAGGGATCTCATCGG |
| LOC_Os08g39840 R | GTTGCTTTCTCCTTCCCGGTCTTC |
| LOC_Os08g39850 F | AAGGGCTTCTCAACAGCCTGAG |
| LOC_Os08g39850 R | CTTCTTCTTCCCTGTCTTCGCTTC |
| LOC_Os09g28160 F | CAGGAACGAAAGCTATGCAGGTC |
| LOC_Os09g28160 R | AGATGCTTGTATGCCATCTCCAC |
| LOC_Os10g25230 F | AAGGACCGGTGATCATCTTGGC |
| LOC_Os10g25230 R | GACGGACGGTCAAACATTGGATAC |
| LOC_Os11g40570 F | TTAGCGAGAGAGGTTGGGCAGTAG |
| LOC_Os11g40570 R | GCGTTGACGTGAGTGATGAGATTC |
| **Primers used for qRT-PCR in Arabidopsis** | |
| **Primer name** | **Sequence (5'-3')** |
| AtActin2 F | TCTTCCGCTCTTTCTTTCCAAGC |
| AtActin2 R | ACCATTGTCACACACGATTGGTTG |
| AtUBQ5 F | AAGAAGACTTACACCAAGCCGAAG |
| AtUBQ5 R | ACAGCGAGCTTAACCTTCTTATGC |
| AtPR2 F | TCTTGAACCCACTTGTCGGC |
| AtPR2 R | GGCTCTGACATCGAGCTCATC |
| AtPR5 F | TCCTTGACCGGCGAGAGTT |
| AtPR5 R | AGGAACAATTGCCCTACCACC |
| AtGSL5 F | CCACCACGAGTACATTCAGGTC |
| AtGSL5 R | GTACACATCTCGGCTGAGAACC |
| AtPDF1.2 F | CTTGTTCTCTTTGCTGCTTTCGAC |
| AtPDF1.2 R | TTGGCTCCTTCAAGGTTAATGCAC |
| AtWRKY33 F | CTTCCACTTGTTTCAGTCCCTCTC |
| AtWRKY33 R | CTGTGGTTGGAGAAGCTAGAACG |
| AtSARD1 F | AGAATCCCTCAACCAGCCCTAC |
| AtSARD1 R | GTGGCTCGCAGCATATTGTTGG |
| AtCBP60G F | CGATAGGACCTTTGTGGGTCATCC |
| AtCBP60G R | ACTTCCTTGAAAGTCGATGTGCTG |
| AtNPR3 F | TCAGCGGCGGCTTTGTAACTTTG |
| AtNPR3 R | CCTCGCCACTCTCTCAATACACTG |
| AtWRKY38 F | ACTGCGAAGCAAGAAAGCATGAAC |
| AtWRKY38 R | TGGTGGCCAAAGTAAGTGGTTCG |
| AtSH3 F | TATCGGCGACCAAATGCAGGTC |
| AtSH3 R | ACTACGGCTCTATGGAGCACAC |
| AtSID2 F | GCTTGGCTAGCACAGTTACAGC |

| | |
|---|---|
| AtSID2 R | CACTGCAGACACCTAATTGAGTCC |

| Supplemental Table 5: Accession numbers of genes used in this study | | | |
|---|---|---|---|
| **Rice** | | | |
| **Gene name** | **MSU ID[a]** | **RAP ID[b]** | **Gene Family** |
| WAKL21 | LOC_Os12g40419 | Os12g0595800 | Wall associated kinase |
| PR10/PBZ14 | LOC_Os12g36830 | Os12g0555000 | Pathogenesis related |
| PAL3 | LOC_Os02g41670 | Os02g0626600 | Phenylalanine ammonia lyase |
| SERK2 | LOC_Os04g38480 | Os04g0457800 | LRR-Receptor like kinase |
| PR1a | LOC_Os07g03710 | Os07g0129200 | Pathogenesis related |
| PR10a/PBZ1 | LOC_Os12g36880 | Os12g0555500 | Pathogenesis related |
| Actin1 | LOC_Os03g50885 | Os03g0718100 | Actin |
| GAPDH | LOC_Os04g40950 | Os04g0486600 | Glyceraldehyde 3-phosphate dehydrogenase |
| bHLH116 | LOC_Os01g50940 | Os01g0705700 | Basic helix-loop-helix transcription factor |
| N.D. | LOC_Os02g43790 | Os02g0654700 | ethylene-responsive transcription factor |
| N.D. | LOC_Os03g08310 | Os03g0180800 | ZIM domain-containing protein |
| N.D. | LOC_Os03g08330 | Os03g0181100 | ZIM domain-containing protein |
| PUB41 | LOC_Os03g13740 | Os03g0240600 | ubiquitin ligase |
| N.D. | LOC_Os03g55800 | Os03g0767000 | Allene oxide synthase |
| OsRERJ1 | LOC_Os04g23550 | Os04g0301500 | Basic helix loop helix transcription factor |
| PUB38 | LOC_Os04g35680 | Os04g0437300 | ubiquitin ligase |
| CRR3 | LOC_Os05g03620 | Os05g0127300 | Crinkly4 subfamily protein |
| N.D. | LOC_Os06g44250 | Os06g0652200 | haemolysin-III |
| CHIT7/PR3 | LOC_Os06g51050 | Os06g0726100 | Chitinase |
| N.D. | LOC_Os07g09420 | Os07g0192000 | AAA-type ATPase |
| N.D. | LOC_Os07g32010 | Os07g0503300 | UDP-glucoronosyl domain containing |
| N.D. | LOC_Os07g36560 | Os07g0550600 | transferase |
| N.D. | LOC_Os08g02700 | Os08g0120600 | fructose-bisphosphate aldolase |
| N.D. | LOC_Os08g36920 | Os08g0474000 | AP2 transcription factor |
| N.D. | LOC_Os08g39840 | Os08g0508800 | Lipoxygenase |
| N.D. | LOC_Os08g39850 | Os08g0509100 | Lipoxygenase |
| N.D. | LOC_Os09g28160 | Os09g0454600 | Mitochondrial phosphate carrier |
| N.D. | LOC_Os10g25230 | Os10g0391400 | ZIM domain-containing protein |
| N.D. | LOC_Os11g40570 | Os11g0621000 | viral response family protein |
| **Arabidopsis** | | | |
| **Gene name** | **TAIR ID[c]** | | |
| Actin2 | AT3G18780 | | |
| PR2 | AT3G57260 | | |
| PR5 | AT1G75040 | | |
| GSL5 | AT4G03550 | | |

| | |
|---|---|
| WRKY33 | AT2G38470 |
| PDF1.2a | AT5G44420 |
| CBP60G | AT5G26920.1 |
| SH3 | AT4G10500 |
| NPR3 | AT5G45110.1 |
| WRKY38 | AT5G22570 |
| SARD1 | AT1G73805 |
| SID2 | AT1G74710 |
| UBQ5 | AT3G62250 |
| BRI1 | AT4G39400 |
| PSKR1 | AT2G02220 |
| WAKL10 | AT1G79680 |
| PEPR1 | AT1G73080 |
| PEPR2 | AT1G17750 |

[a]: Gene ID in Michigan State University database
[b]: Gene ID in Rice Annotation Project database

[c]: Gene ID in The Arabidopsis Information Resource database

N.D.: Name not defined

### Supplemental Figure S1

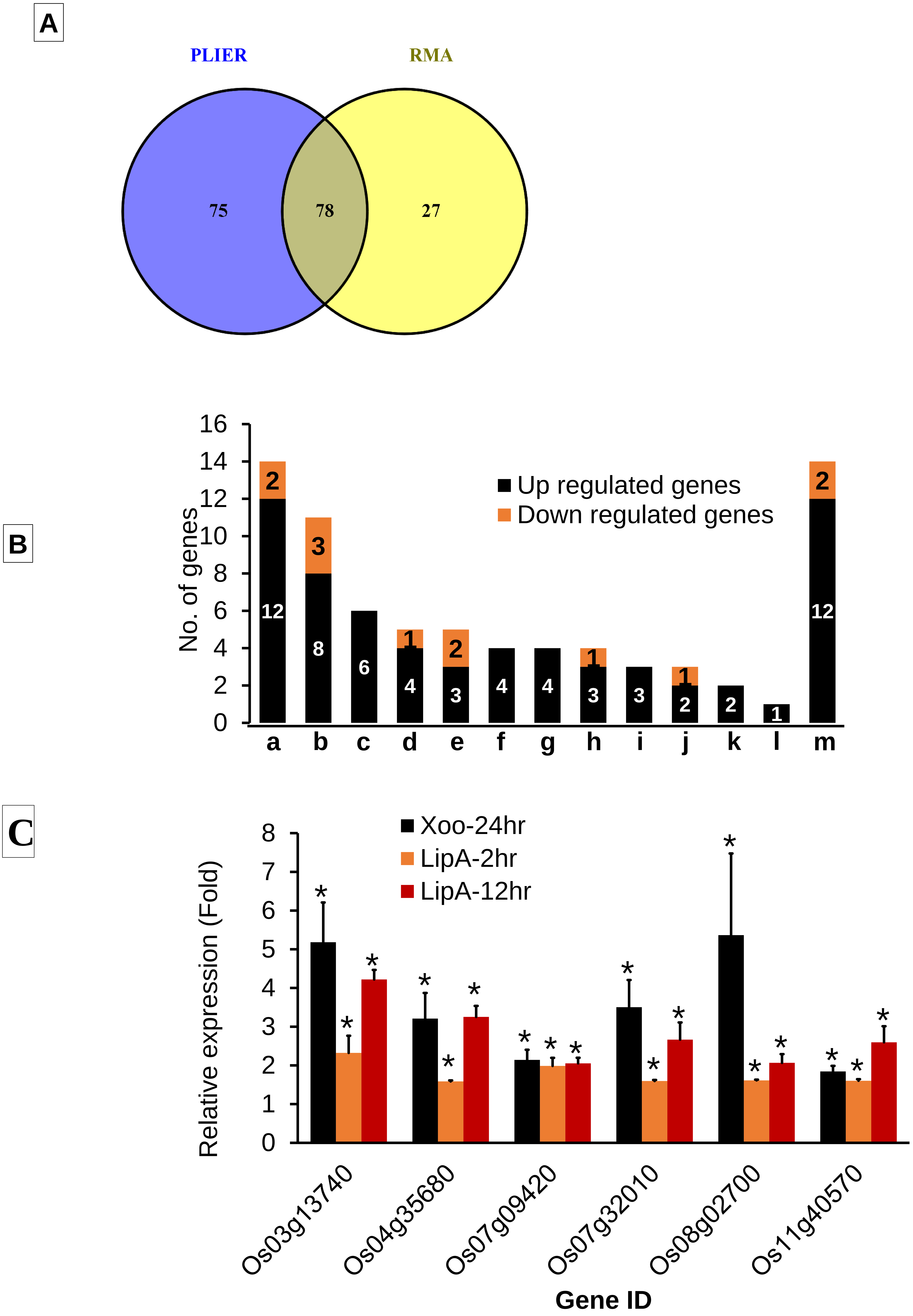

**Supplemental Figure S1: Transcriptome profiling of rice leaves after treatment with LipA.**

(A) Venn diagram indicating number of unique and common genes differentially expressed after 2hr of LipA treatment using RMA and PLIER16 analysis.

(B) Functional categorization using MapMan tool showing number of upregulated and downregulated genes in various functional categories. a: Signal transduction, b: Transcription/transcription factors, c: Defense, d: Hormone metabolism, e: Transport, f: Stress, g: Protein synthesis/turnover, h: Metabolism, i: Secondary metabolites, j: Redox, k: Cell wall structure/synthesis, l: Transposons, m: Others.

(C) qRT-PCR validation of six differentially expressed genes after 2hr and 12hr of LipA treatment and also after 24hr of *Xoo* treatment. 12-14 days old rice leaves were infiltrated either with LipA (0.5mg/ml) or *Xoo* (O.D. 1.0). Each bar represents average value and error bar denotes standard error (SE) of at least three different experiments. Relative expression was calculated in leaves treated with LipA or *Xoo* with respect to leaves treated with buffer. Asterisk (*) represents significant difference in fold change with $p<0.05$. *OsActin1* was used as internal control for qRT-PCR. The relative fold change was calculated by using $2^{-\Delta\Delta Ct}$ method.

### Supplemental Figure S2

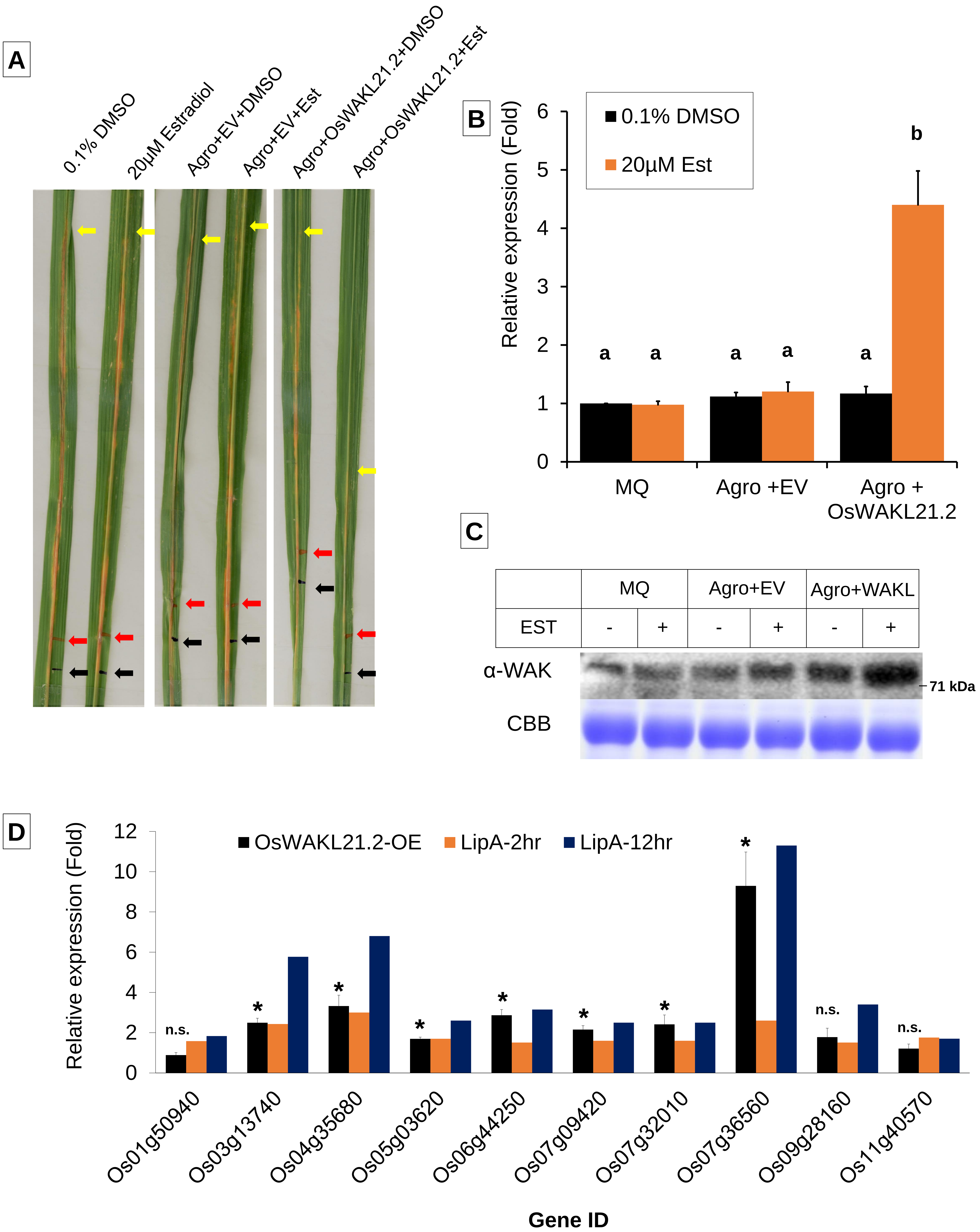

**Supplemental Figure S2: Overexpression of *OsWAKL21.2* induces rice immune responses:**

(A) Representative image of lesion length caused by *Xoo*. The black mark indicate point of Agrobacterium injection while red mark indicate point of pin prick inoculation of *Xoo*. The yellow mark indicate the last point of disease progression at 10dpi. The quantification of lesion length in an experiment is mentioned in Figure 2C.

(B) qRT-PCR indicating relative expression of *OsWAKL21.2* in rice leaves after 24hr of injection with either MQ (Mock), Agrobacterium containing empty vector or Agrobacterium containing wild type *OsWAKL21.2* under uninduced (0.1% DMSO) and induced (20μM estradiol) condition. Expression level in leaves treated with mock (0.1% DMSO) was considered as 1 and relative expression was calculated with respect to it. small letters (a, b and c) above the bars indicates significant difference with $p<0.05$.

(C) Western blot performed from protein isolated from rice mid-veins samples as mentioned in the Supplemental figure S2B. Anti OsWAKL21.2$_{376\text{-}725}$ antibody (α-WAK) was used for Western blotting. CBB indicate Coomassie brilliant blue staining of gel ran parallelly for loading control.

(D) Relative expression of ten LipA responsive genes either after *OsWAKL21.2* overexpression or 2hr and 12hr post LipA treatment. These genes are LOC_Os03g13740 (a ubiquitin ligase, *OsPUB41*), LOC_Os04g35680 (a ubiquitin ligase, Os*PUB38*), LOC_Os05g03620 (a Crinkly4 subfamily protein, *OsCRR3*), LOC_Os06g44250 (a haemolysin-III), LOC_Os07g09420 (An AAA-type ATPase), LOC_Os07g32010 (a UDP-glucoronosyl and UDP-glucosyl transferase domain-containing protein), LOC_Os07g36560 (a transferase family protein), LOC_Os01g50940 (a Myc-type, basic helix-loop-helix (bHLH) domain-containing protein), LOC_Os09g28160 (a mitochondrial phosphate carrier protein) and LOC_Os11g40570 (a plant viral response family protein). Expression change after *OsWAKL21.2* overexpression was analyzed by qRT-PCR while 2hr and 12hr post LipA treatment data represent the microarray data. For each gene, transcript level of uninduced condition (treatment with Agrobacterium carrying *OsWAKL21.2* with 0.1% DMSO) was considered as 1 and was compared to induced condition (treatment with Agrobacterium carrying *OsWAKL21.2* with 20μM estradiol). Asterisk (*) represents significant difference with respect to control with $p<0.05$. n.s. indicate not significant difference in relative expression.

In B and D, each bar represents average value and error bar denotes standard error (SE) of at least three different experiments. *OsActin1* was used as internal control for qRT-PCR. The relative fold change was calculated by using $2^{-\Delta\Delta Ct}$ method.

### Supplemental Figure S3

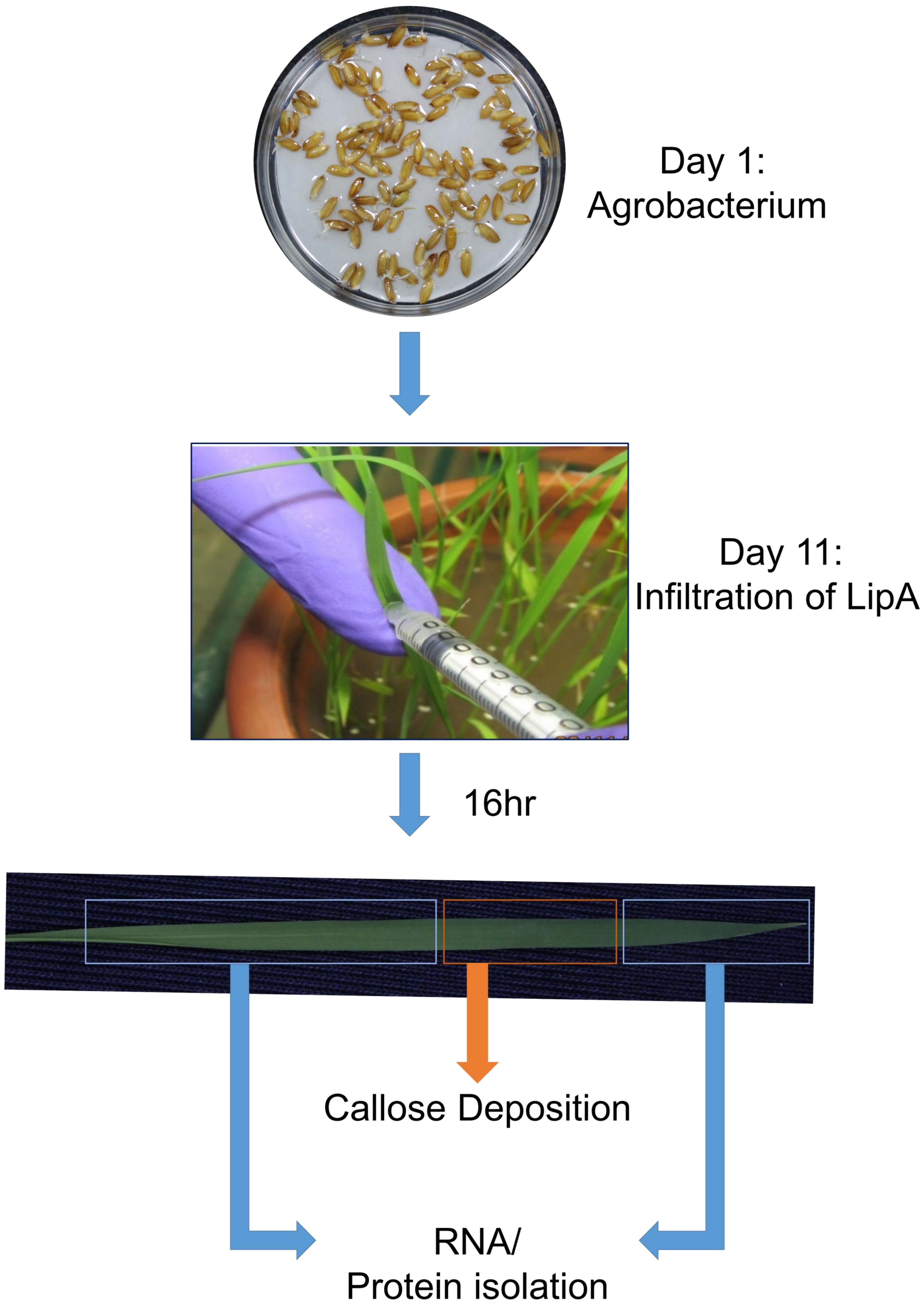

**Supplemental Figure S3: Methodology for downregulation of *OsWAKL21.2* in rice seedlings using Virus Induced Gene Silencing (VIGS).**
One day old seedlings were submerged in Agrobacterium suspension carrying VIGS vector for 24hr and washed subsequently. Seedlings were grown on water and third leaf was infiltrated with LipA (0.5mg/ml) on day 11. After 16hr, infiltrated zone (in orange box) was collected for visualization of callose deposition while the rest of the leaf (in blue box) was collected for Western blot or qRT-PCR analysis.

### Supplemental Figure S4

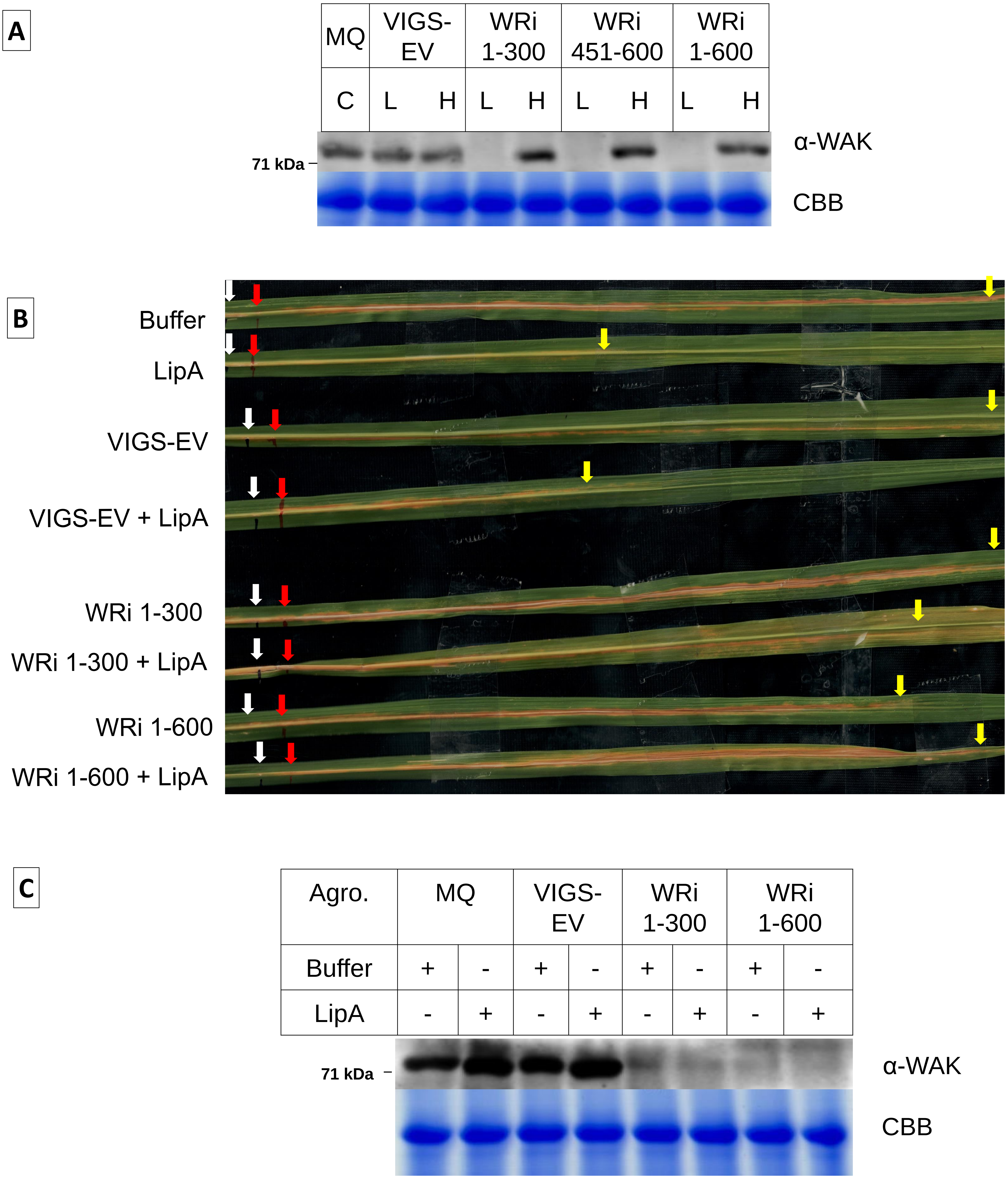

**Supplemental Figure S4: Transient downregulation of *OsWAKL21.2* in rice**

(A) Western blot indicating protein level in representative leaves whose transcript level was shown in Figure 3D. 'L' indicates protein isolated from leaves showing low callose deposition while 'H' indicates protein isolated from leaves showing high callose deposition. C indicates protein from leaves that were never treated with Agrobacterium or LipA.

(B) Lesion length caused by *Xoo* after 10 days of pin prick inoculation on rice leaves when leaves were pre-injected with respective treatments (Buffer, LipA, or Agrobacterium along with buffer/LipA). The white mark indicate point of Agrobacterium injection while red mark indicate point of pin prick inoculation of *Xoo*. The yellow mark indicate the last point of disease progression at 10dpi. The quantification of lesion length is shown in Figure 3E.

(C) Western blot indicating protein level in representative leaves whose transcript level was shown in Figure 3F. Protein was isolated from 10-12 leaves after 24hr of respective treatment.

In A and C anti-OsWAKL21.2$_{376\text{-}725}$ antibody (α-WAK) was used for Western blotting. CBB indicate Coomassie brilliant blue staining of gel ran parallelly for loading control.

### Supplemental Figure S5

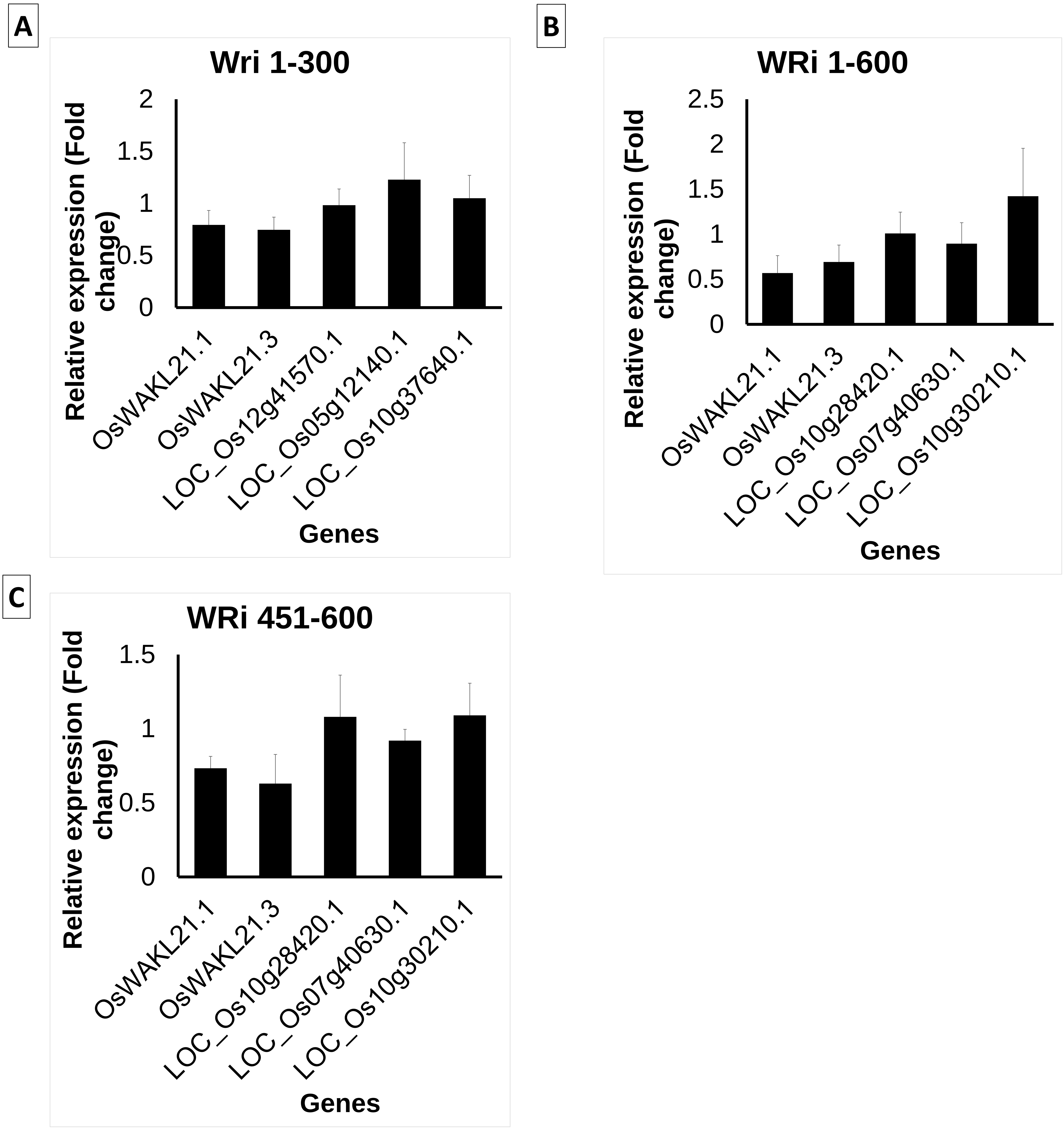

**Supplemental Figure S5: VIGS mediated transient downregulation of *OsWAKL21.2* does not have significant effect on expression of predicted off-targets genes**
Expression of genes showing highest identity with respective *OsWAKL21.2* sequence used for VIGS was also tested following treatment with OsWAKL21.2-RNAi constructs. RNA that was used to test expression of OsWAKL21.2 in Fig. 3 F was used for qRT-PCR. Each bar represents average of three independent experiments, n>10 in each experiment. Transcript level of buffer injected leaves was considered as 1 and fold change in Agrobacterium containing respective RNAi vector treated leaves was calculated with respect to it.

### Supplemental Figure S6

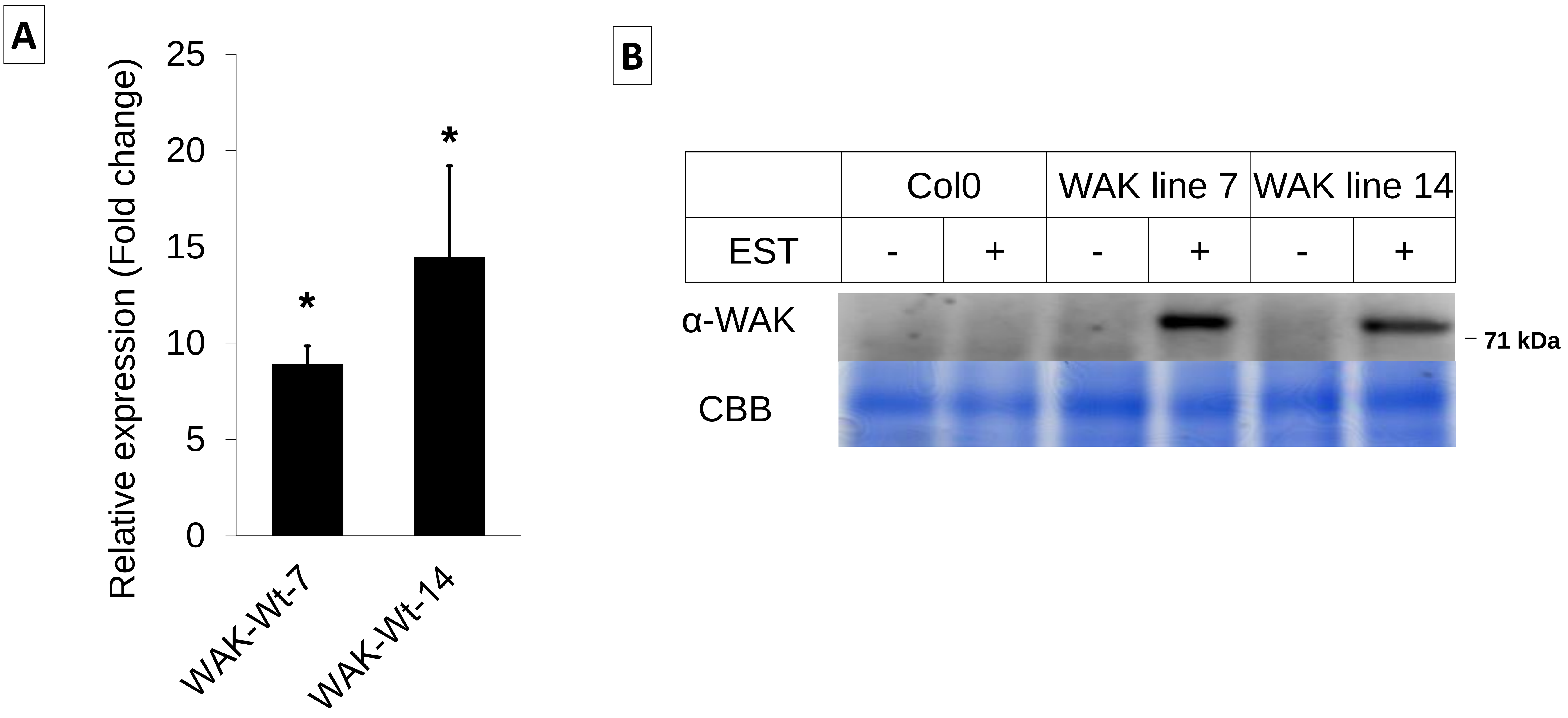

**Supplemental Figure S6: qRT-PCR and Western blot validation for ectopically expressing *OsWAKL21.2* transgenic Arabidopsis plants.**

(A) qRT-PCR indicate induction of expression of *OsWAKL21.2* in transgenic Arabidopsis lines (Line 7 and line 14). Expression in 0.1% DMSO treated leaves was considered as 1 and relative expression in 20µM estradiol treated leaves was calculated with respect to it. Each bar represents the average of three independent experiments for each line. *AtActin2* was used as internal control for qRT-PCR. The relative fold change was calculated by using $2^{-\Delta\Delta Ct}$ method.

(B) Estradiol treatment induces protein level of OsWAKL21.2 in Arabidopsis transgenic lines expressing *OsWAKL21.2* under estradiol inducible promoters. Est+ indicate infiltration with 20µM estradiol while Est- indicate control that is infiltrated with 0.1% DMSO. Anti $OsWAKL21.2_{376\text{-}725}$ antibody (α-WAK) was used for Western blotting. CBB indicate Coomassie brilliant blue staining of gel ran parallelly for loading control. Asterisk (*) represents significant difference in expression with $p<0.05$.

In an experiment, for each treatment, RNA/protein was isolated from 3 leaves of different plants. Leaves were collected after 12hr of infiltration with DMSO/Est.

### Supplemental Figure S7

A

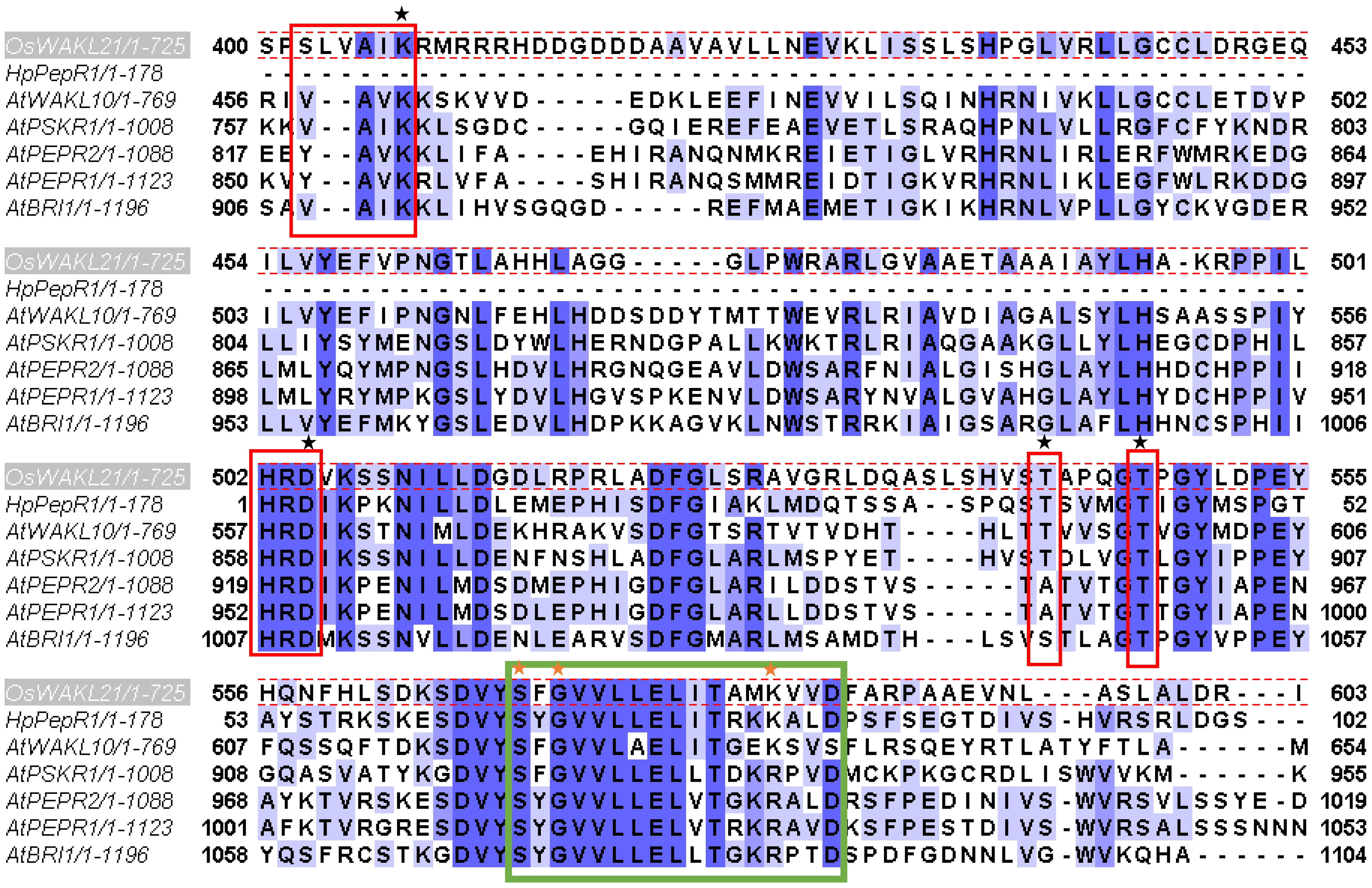

B

Hits found:

| Hit # | Position | GCC sequence | GCC Hydrophobicity | GCC Molecular Weight | GCC Isoelectric point | GCC Mean |
|---|---|---|---|---|---|---|
| 1 | 506-522 | KSSNILLDGDLRPRLAD | 0.563 | 0.796 | 0.8 | 0.72 |
| 2 | 569-585 | SFGVVLLELITAMKVVD | 0.706 | 0.856 | 0.918 | 0.827 |

C

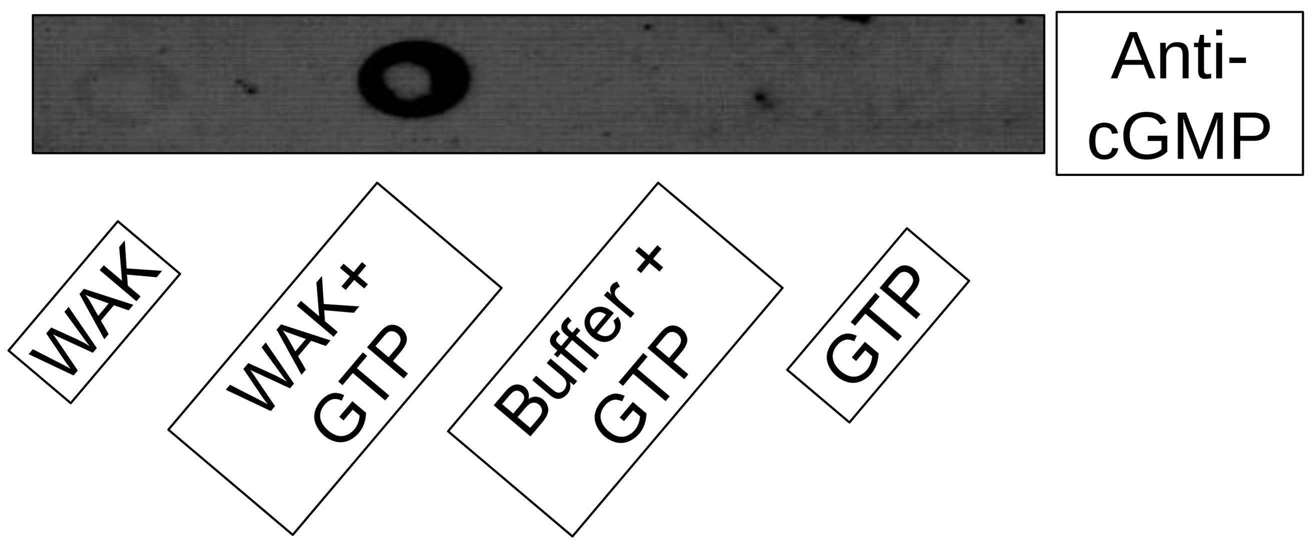

**Supplemental Figure S7: Biochemical characterization of OsWAKL21.2**

(A) Alignment of kinase domain of OsWAKL21.2 with other known plant moonlighting receptor kinases that have GC activity. The residues highlighted in red box indicate key residues for kinase activity. The residues highlighted in green box indicate GC motif. Black star above the residues indicate residues substituted with Alanine (A) in kinase deficient mutant. The orange star above the residues indicate residues mutated in GC deficient mutant. S569 and F571 were substituted with Alanine while K582 was substituted with Glutamine as previously done by Ma et al., 2012.

(B) Score obtained after submission of amino acid sequence of OsWAKL21.2 for prediction of GC motif using Gcpred tool (http://gcpred.com/). The second predicted GC motif (569-685) have high probability of having GC activity as score in all the parameters is above required cut-off score (so indicated by green color by tool).

(C) Dot blot performed using anti cGMP antibody after GC assay. 50μg of purified protein (WAK) was incubated with/without GTP for 12hr and subsequently used for dot blot. GTP alone and GC buffer + GTP were used as controls. The blot was probed using anti cGMP antibody 9Sigma).

### Supplemental Figure S8

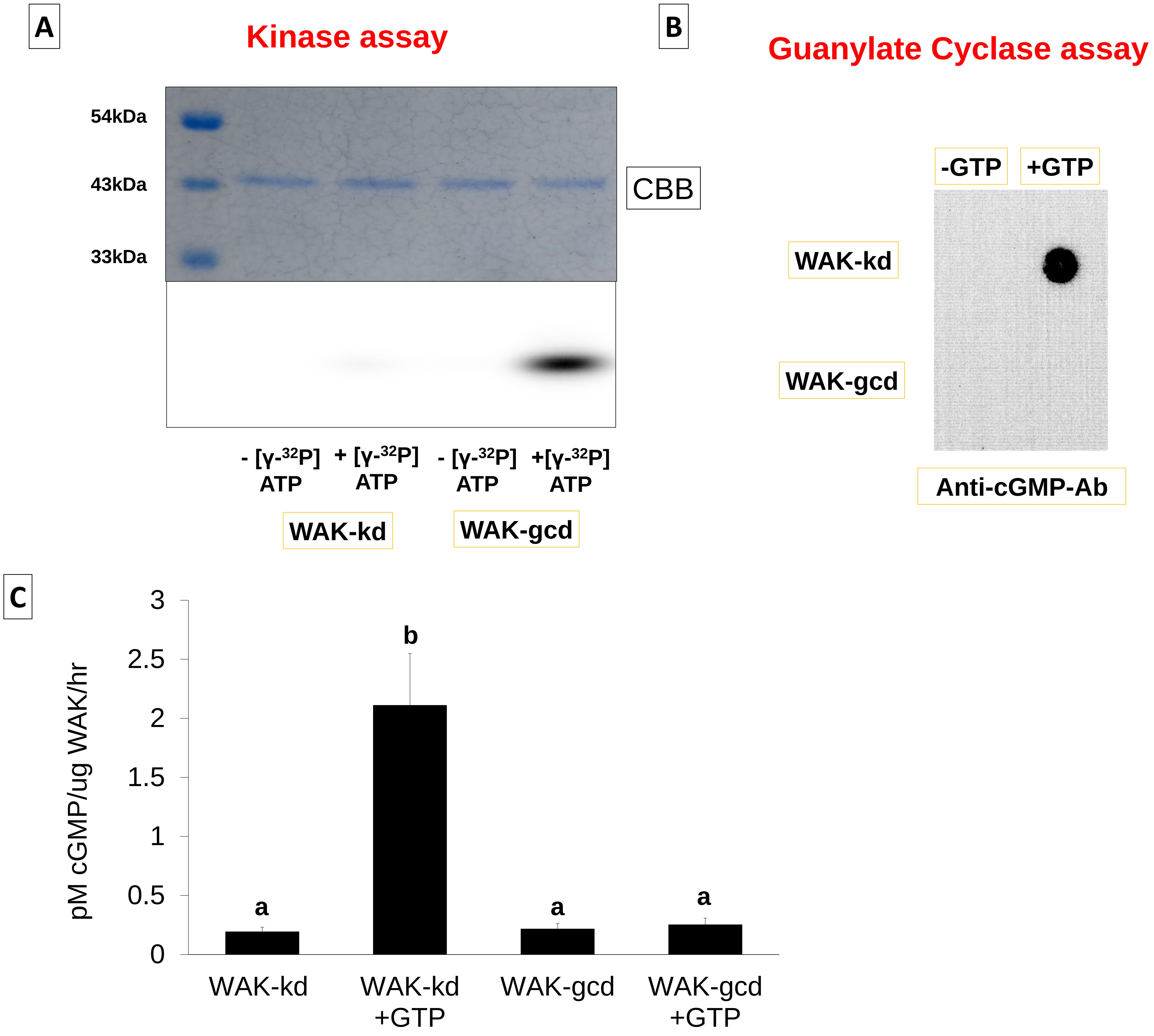

**Supplemental Figure S8: Biochemical activities of purified kinase domain of mutant versions of OsWAKL21.2**

(A) Kinase assay using kinase deficient (WAK-kd) or guanylate cyclase deficient (WAK-gcd) versions of OsWAKL21.2 kinase domain. 50µg of purified protein was incubated with radiolabeled ATP ($\gamma$-$^{32}$P] ATP) for 1hr and used for autoradiography. CBB indicate Coomassie brilliant blue staining of gel ran parallelly for loading control.

(B) Qualitative GC assay performed using WAK-kd or WAK-gcd mutants of OsWAKL21.2 kinase domain. 50µg of purified protein was incubated with/without GTP for 12hr and subsequently used for dot blot. The blot was probed using anti cGMP antibody.

(C) Quantification of cGMP produced by kinase domain of WAK-kd or WAK-gcd after 1hr of incubation with cGMP. Each bar represents average and error bar indicate standard error of three individual experiments. Small letters (a and b) above the bars indicate significant difference with $p<0.05$. All experiments were repeated three times and similar results were obtained.

### Supplemental Figure S9

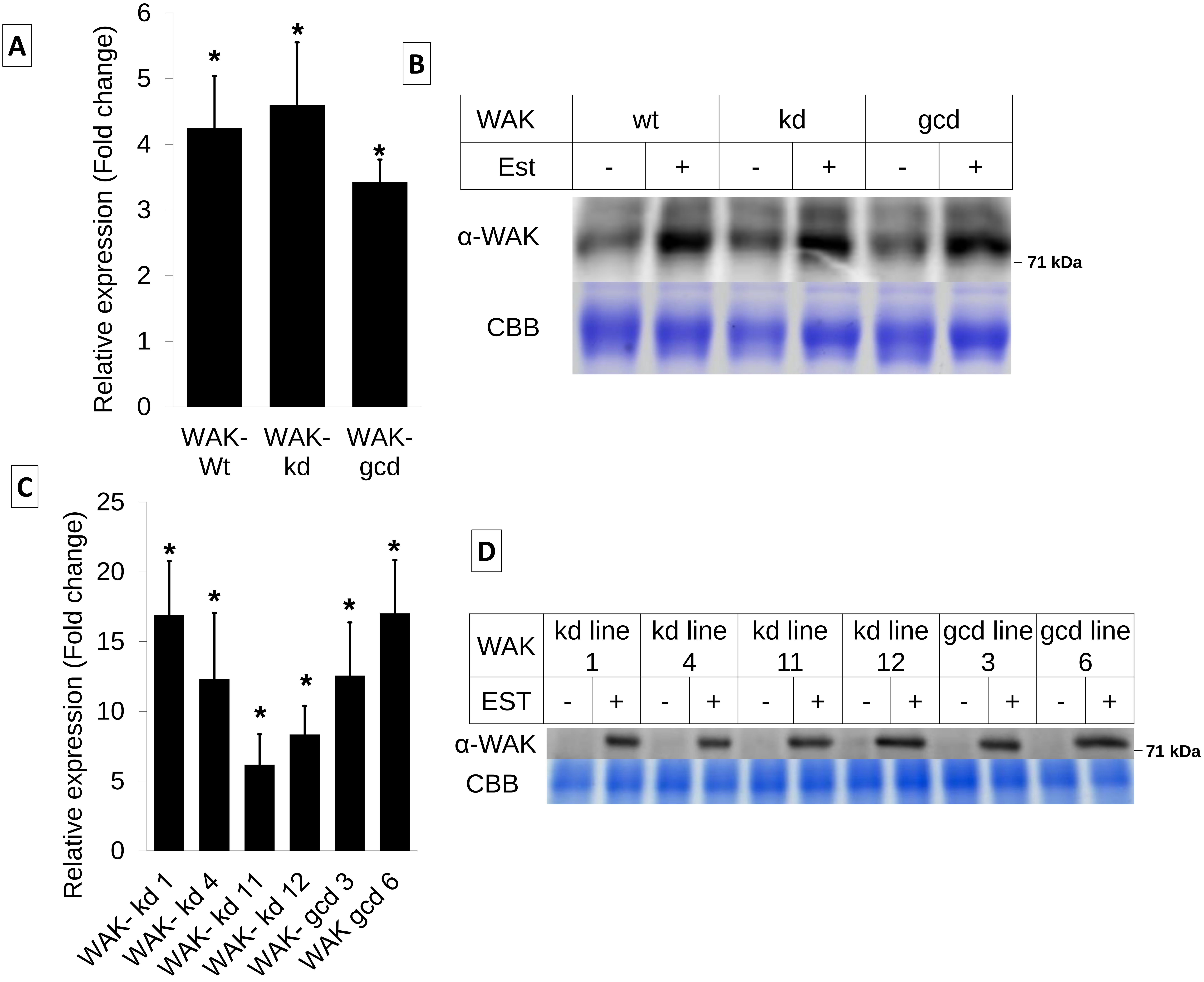

**Supplemental Figure S9: qRT-PCR and Western blot validation of expression of mutant versions of *OsWAKL21.2* by transient transformation in rice and ectopic expression in Arabidopsis transgenic lines.**

(A) qRT-PCR indicate overexpression of *OsWAKL21.2* and mutants (WAK-kd and WAK-gcd) in rice following Agrobacterium mediated transient transformation. *OsActin1* was used as the internal control.

(B) Western blot indicating protein level of OsWAKL21.2 or its mutants after Agrobacterium mediated transient transformation in rice leaves.

(C) qRT-PCR indicating induced expression of ectopically expressing *OsWAKL21.2* mutants [*OsWAKL21.2-kd* (WAK-kd) or *OsWAKL21.2-gcd* (WAK-gcd)] in respective transgenic Arabidopsis lines. Four different lines (line 1, 4, 11 and 12) were used for WAK-kd while two different lines (line 3 and 6) were used for WAK-gcd. *AtActin2* was used as the internal control.

(D) Western blot indicating induced expression of ectopically expressing OsWAKL21.2 mutants [Os*WAKL21.2*-kd (WAK-kd) or Os*WAKL21.2*-gcd (WAK-gcd)] in respective transgenic Arabidopsis lines. Four different lines were used for WAK-kd while two different lines were used for WAK-gcd as mentioned in Supplemental Figure S9C.

In B and D Est+ indicate infiltration with 20µM estradiol while Est- indicate control that is infiltrated with 0.1% DMSO. In A and C Expression level in leaves treated with control (0.1% DMSO) was considered as 1 and expression level in leaves treated with 20µM estradiol was calculated with respect to it. Asterisk (*) represents significant difference in expression with $p<0.05$. In A and B, Samples were collected after 24hr of treatment with Agrobacterium constructs. In C and D, samples were collected after 12hr of infiltration of DMSO/Est. In B and D anti-OsWAKL21.2$_{376\text{-}725}$ antibody (α-WAK) was used for Western blotting. CBB indicate Coomassie brilliant blue staining of gel ran parallelly for loading control.

### Supplemental Figure S10

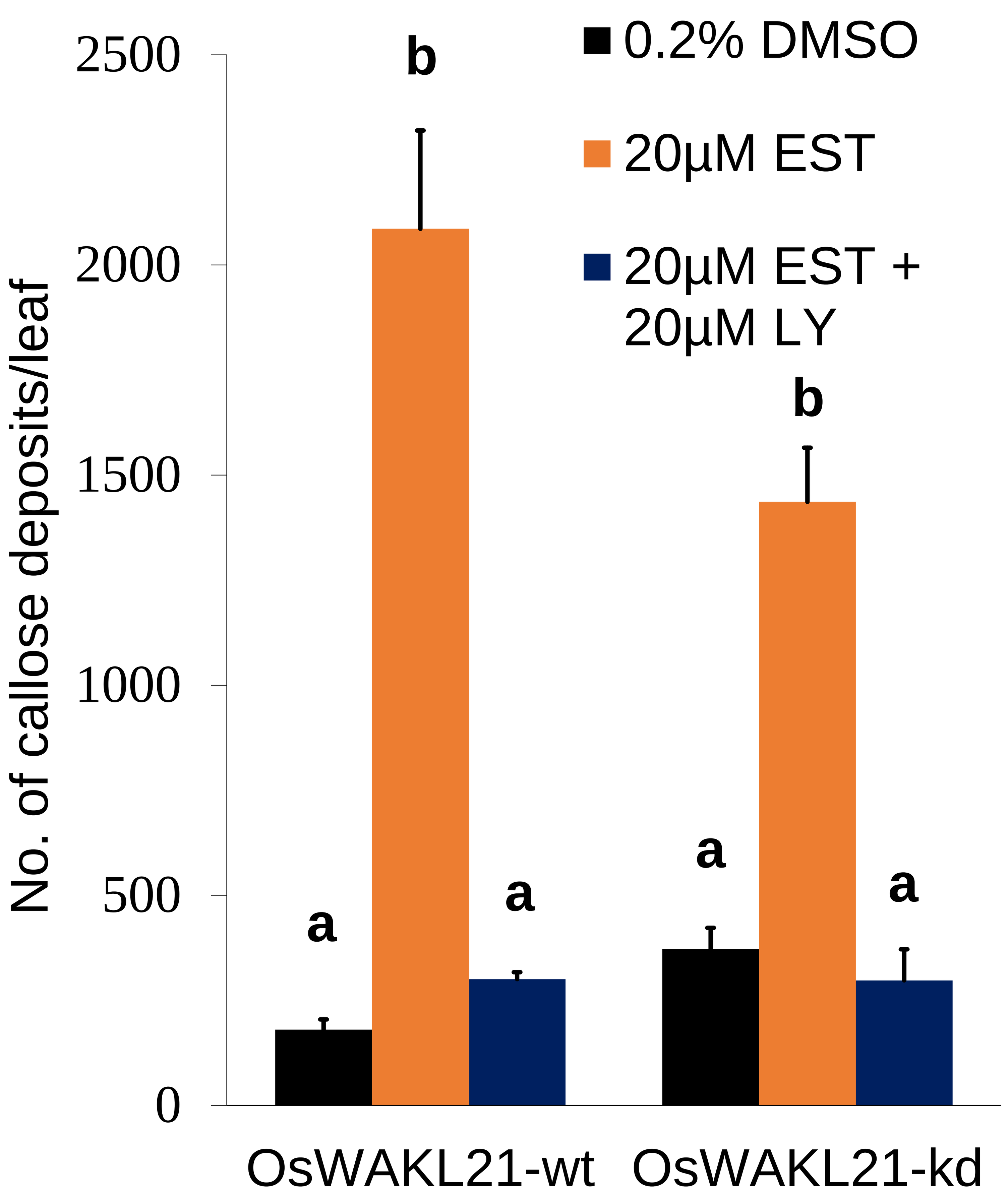

**Supplemental Figure S10: Treatment with GC inhibitor attenuates *OsWAKL21.2* induced callose deposition in transgenic Arabidopsis leaves.**

Leaves of *OsWAKL21.2* or Os*WAKL21.2*-kd transgenic plants were treated with either 0.2%DMSO, 20µM estradiol or 20µM estradiol + 20µM GC inhibitor (LY83583 or LY). Each bar represents the average and error bar represents SE of three different leaves for each treatment in an experiment. Small letters (a and b) above the bars indicate significant difference with $p<0.05$. The experiment was repeated three times and similar results were obtained.

### Supplemental Figure S11

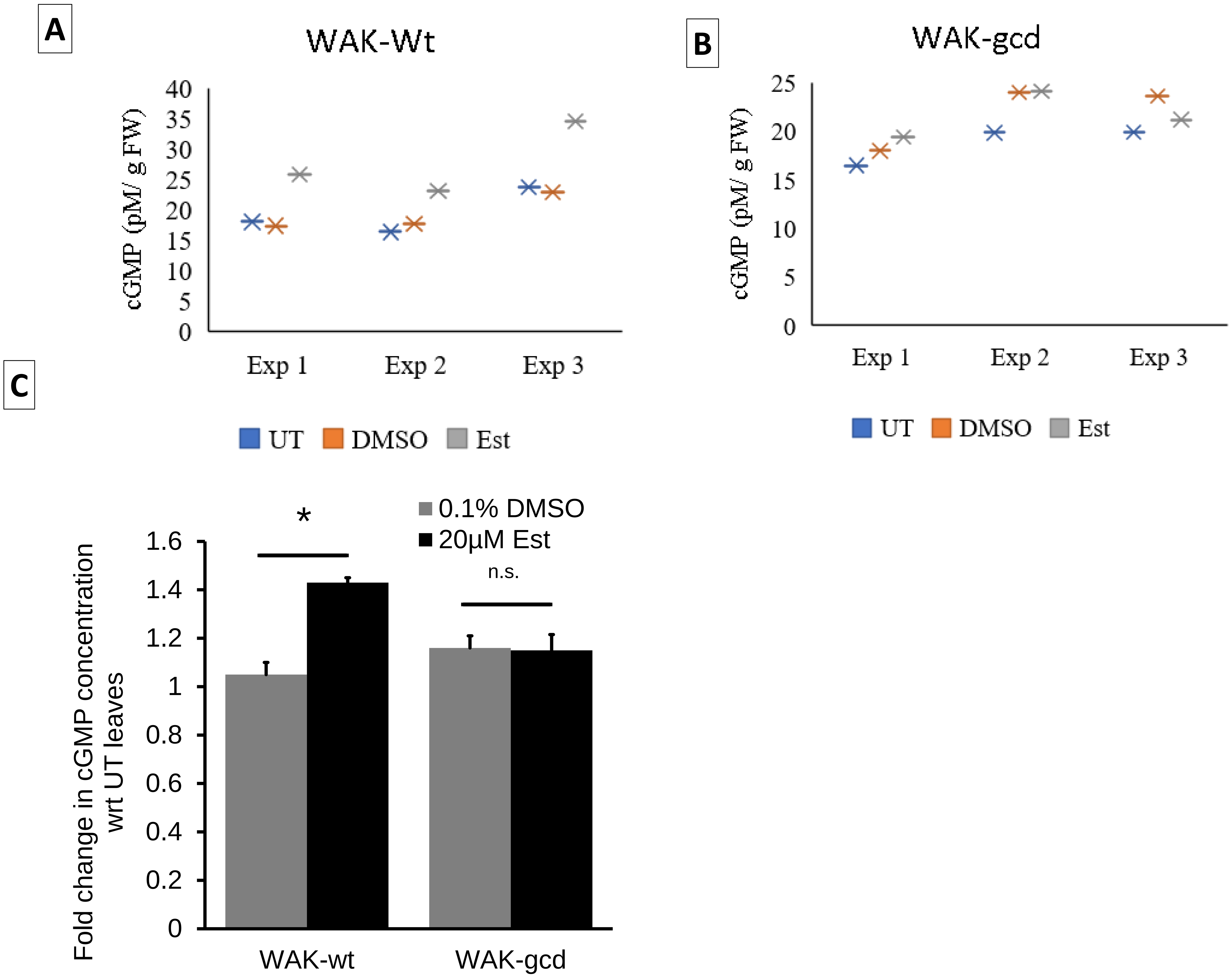

**Supplemental Figure S11: Ectopic expression of *OsWAKL21.2* in Arabidopsis enhances *in planta* cGMP level by its GC activity.**

(A) Concentration of cGMP in transgenic Arabidopsis plants ectopically expressing *OsWAKL21.2* after 3hr of treatment of 0.1% DMSO, 20µM estradiol or untreated (UT) control plants in three different experiments.

(B) Concentration of cGMP in transgenic Arabidopsis plants ectopically expressing *OsWAKL21.2-gcd* 3hr post treatment of 0.1% DMSO, 20µM estradiol or untreated (UT) control plants in three different experiments.

(C) Fold change in the in planta cGMP concentration 3hr post treatment of respective transgenic plants with either DMSO or estradiol with respect to untreated plants. In each experiment, cGMP concentration in untreated samples was considered 1 and fold change after 3hr of treatment with either 0.1% DMSO or 20µM estradiol was calculated with respect to it. n.s. indicate not significant difference in relative expression.

(D) Western blot from Arabidopsis leaves from sister lines carrying either *OsWAKL21.2*, *NahG* or both. Est+ indicate infiltration with 20µM estradiol while Est- indicate control that is infiltrated with 0.1% DMSO. Anti OsWAKL21.2$_{376\text{-}725}$ antibody (α-WAK) was used for Western blotting. CBB indicate Coomassie brilliant blue staining of gel ran parallelly for loading control.

### Supplemental Figure S12

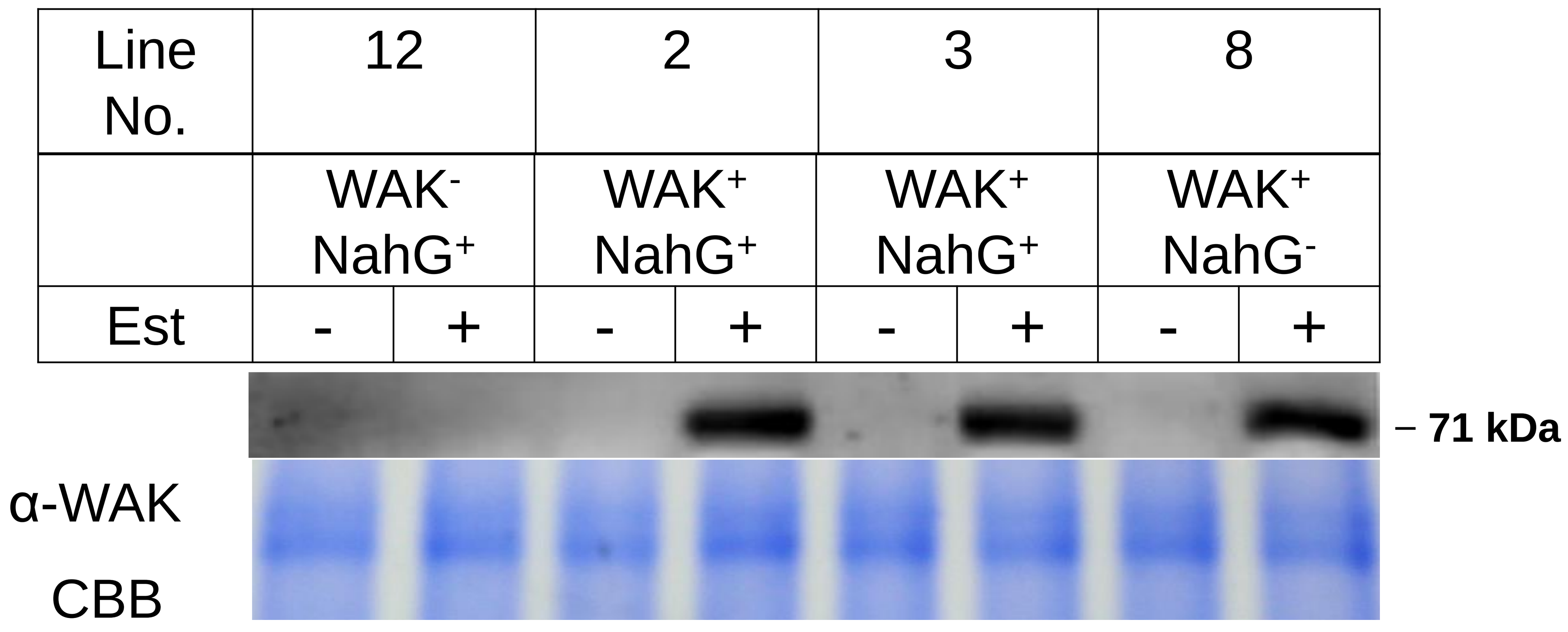

**Supplemental Figure S12: Western blot validation of ectopic expression of *OsWAKL21.2* in Arabidopsis transgenic lines generated after crossing with *NahG* lines.**
Western blot from Arabidopsis leaves from sister lines carrying either *OsWAKL21.2*, *NahG* or both. Est+ indicate infiltration with 20µM estradiol while Est- indicate control that is infiltrated with 0.1% DMSO. Anti OsWAKL21.2$_{376\text{-}725}$ antibody (α-WAK) was used for Western blotting. CBB indicate Coomassie brilliant blue staining of gel ran parallelly for loading control.
